# DNA branch migration across intact membranes powers an inside-out GPCR analog for lysis-free detection of intracellular RNA with on-membrane amplification

**DOI:** 10.1101/2025.10.29.685379

**Authors:** Ranjan Sasmal, Saanya Yadav, Gde Bimananda Mahardika Wisna, Nirbhik Acharya, Carter Swanson, Hao Yan, Himanshu Joshi, Rizal F. Hariadi

**Affiliations:** Center for Molecular Design and Biomimetics at the Biodesign Institute, Arizona State University, Tempe, Arizona, USA; Department of Biotechnology, Indian Institute of Technology, Hyderabad, India; Department of Physics, Arizona State University, Tempe, Arizona, USA; School of Molecular Sciences, Arizona State University, Tempe, Arizona, USA

**Keywords:** amphiphilic DNA, DNA nanotechnology, DNA branch migration, transmembrane signaling, synthetic receptor (GPCR mimic), hybridization chain reaction, live-cell biosensing, signal amplification

## Abstract

Interrogating the sequence-specific DNA/RNA content of a living cell without destroying it, is an elusive goal in biology and diagnostics with transformative potential. A DNA/RNA-based approach, however, must relay the recognition event across the intact lipid bilayer, a step that has long been considered impossible because hydrophilic, massively charged DNA/RNA is incompatible with the hydrophobic membrane interior. Here, we show that DNA branch migration can be driven across a lipid bilayer, converting an intracellular nucleic-acid sequence into an amplified, extracellular optical signal while preserving cell viability and compartmentalization, a synthetic transmembrane signal transduction analogous to natural receptors, such as G-protein-coupled receptors (GPCRs), but operating in reverse. We demonstrate this principle with Hybridization Across Lipid for Oligonucleotide Sensing (HALOS), an amphiphilic DNA hairpin comprising a toehold for recognition, a stem for stability, a loop, and cholesterols for transmembrane anchoring. Upon binding to a sequence-specific nucleic acid target, toehold-mediated strand invasion drives DNA branch migration across the bilayer, switching HALOS from a closed to an open conformation that relays the signal to the opposite face. Molecular dynamics simulations revealed that a cholesterol belt stabilizes the DNA stem within the membrane, preserving the hairpin structure necessary for transmembrane signaling. Notably, this branch migration proceeds despite the kinetic barrier of hybridization through the hydrophobic bilayer interior. By combining HALOS with an isothermal hybridization chain reaction (HCR), we established a platform that enables intracellular nucleic acid target detection and amplified fluorescent reporting from outside synthetic vesicles and live mammalian cells, achieving nanomolar-range sensitivity. These results establish DNA branch migration across lipid membranes as a design principle for programmable lysis-free molecular sensing in intact cells.

## Introduction

Interrogating which nucleic-acid sequences are present and expressed inside a living cell, without lysing, fixing, or genetically reprogramming it, is a foundational goal across cell biology, diagnostics, and synthetic biology. Living cells continuously move information in the opposite, outside-in direction, coupling extracellular recognition to an intracellular response without breaching the membrane. ^1–3^ Natural receptors, including G-protein-coupled receptors (GPCRs), achieve this input-output coupling through tightly regulated conformational and biochemical signaling cascades. ^4–7^ Although these receptors regulate diverse physiological processes and represent major therapeutic targets, ^4,8–10^ their native ligand specificity and structural complexity limit their reprogrammability toward arbitrary molecular targets. Engineering *de novo* transmembrane transducers for new targets is hard: it demands selective molecular recognition, membrane-spanning conformational switching, and an extracellular output, all while staying stable in the bilayer. Such synthetic transducers, reprogrammable to arbitrary intracellular targets, could enable a new class of biosensors for real-time, non-destructive monitoring of molecular activity in living cells, yet achieving this requires propagating a sequence-specific hybridization across a hydrophobic bilayer interior incompatible with charged nucleic acids.

Synthetic transmembrane signaling has been pursued along several routes: retargeting natural receptors such as GPCRs, ^11,12^ and building systems from engineered proteins and peptides, ^13–18^ lipid assemblies, and small molecules. ^19–21^ Protein- and peptide-based receptors can provide sophisticated signaling functions, but often require extensive optimization and may face stability or immunogenicity constraints in biological environments.^22–24^ Lipid and small-molecule systems can integrate efficiently into membranes, but are generally less suited for programmable sequence recognition and signal amplification. ^21,25^ None of these approaches, however, reads a sequence-specific signal from inside a living cell and reports it without disrupting it, motivating molecular architectures that combine membrane compatibility, chemical stability, programmable sequence recognition, and amenability to rational design.

DNA nanotechnology has emerged as a unique synthetic platform that integrates programmable molecular recognition, chemical stability, and nanoscale architectural control, making it especially suitable for *de novo* engineering of membrane-spanning functional nanodevices. ^26–32^ Versatile DNA assemblies spanning lipid bilayers have served as synthetic receptors and channels, enabling controlled transport of diverse cargo such as ions, ^33–36^ small molecules, ^37,38^ polymers, ^39^ DNA, ^40,41^ proteins, and lipids. ^42,43^ Furthermore, stimuli-responsive DNA nanopores with gated architectures have been demonstrated to mimic natural ion channels by reversible switching between open and closed states. ^44,45^ However, most DNA membrane systems have been built for molecular transport rather than non-invasive signal transduction. Recent DNA-based nanodevice approaches have used receptor-like switching at membranes, but none have transduced a sequence-specific hybridization across an intact bilayer, coupling recognition on one face to a structural output on the other. Instead, they are triggered by pH (a strand crosses, but the input is protonation, not sequence recognition), ^46–48^ or rely on surface-confined switching in which no strand crosses, with demanding delivery and poor live-cell performance. ^49^ Critically, they also lack the sensitivity to detect low-abundance intracellular targets, indicating a need for signal amplification. Whether such transduction, as opposed to mere strand transport, is achievable has thus remained unresolved.

Here, we report Hybridization Across Lipid for Oligonucleotide Sensing (HALOS), an amphiphilic DNA hairpin that integrates cholesterol-mediated membrane anchoring with programmable hairpin DNA to detect sequence-specific intracellular nucleic acid targets and transduce their recognition across the lipid bilayer in a lysis-free manner. Recognition of an intracellular target by a short single-stranded toehold (an overhang that nucleates strand exchange) triggers strand displacement and drives DNA branch migration (the progressive exchange of base pairs) across the membrane, a step we demonstrate directly. This switches HALOS from a closed hairpin to an open conformation that presents a reporter-binding domain on the opposite face of the membrane for a fluorescent readout. To our knowledge, HALOS is the first synthetic device to combine sequence-specific recognition of an intracellular target, transduction of that recognition across the lipid bilayer by DNA branch migration, and enzyme-free on-membrane signal amplification, and we demonstrated this integrated mechanism in living cells. By coupling the opened hairpin to a non-enzymatic fluorescent hybridization chain reaction (HCR), ^50,51^ we amplified the transduced signal up to 8.2-fold relative to a single non-amplifying hairpin reporter, enabling nanomolar-range detection of a transfected RNA target in live cells. Unlike previous DNA-based transmembrane signaling devices, ^47,49^ which often require pH modulation, vesicle fusion, or complex delivery strategies, HALOS inserts directly into lipid and live-cell membranes via cholesterol anchors and transduces intracellular recognition across the intact bilayer by DNA branch migration. Analogous to a GPCR but operating inside-out, HALOS demonstrates that DNA branch migration can transduce a sequence-specific recognition event across a lipid bilayer, offering a lysis-free strategy for detecting nucleic acid targets delivered into intact cells, with potential applications in non-destructive cellular diagnostics, synthetic and chemical biology, and therapeutic monitoring.

## Results

### HALOS design principle for transmembrane signaling

In GPCR signaling, extracellular ligand binding drives an allosteric change that relays the signal across the bilayer to intracellular cascades ^5,7^ (Fig. 1(a)). Inspired by natural GPCRs, we designed a *de novo* membrane-spanning signal transducer: an amphiphilic DNA hairpin (Fig. 1(b–d)). HALOS comprises three domains: a 10 nt toehold (a) for programmable target recognition, a stem (b-b*) bearing dual cholesterol anchors, and a 10 nt loop (c). The two cholesterols sit 5 nt apart on the stem, ∼180*^◦^* across the helix, favoring stable axial insertion across the bilayer (Fig. 1(c)). Two cholesterol anchors were chosen over other lipid anchors due to their superior membrane affinity, insertion, and stability.^43,52,53^ Analogous to GPCR signaling but functioning in an inside-out manner, HALOS identifies a single-stranded DNA/RNA targets located within the lumen of a lipid bilayer compartment via its toehold region (a). This interaction facilitates a hairpin conformational change across the lipid bilayer, initiating a cascade of DNA reporter strand binding events that amplifies the signal from the exterior of the compartment (Fig. 1(b–c)).

**Fig. 1.**
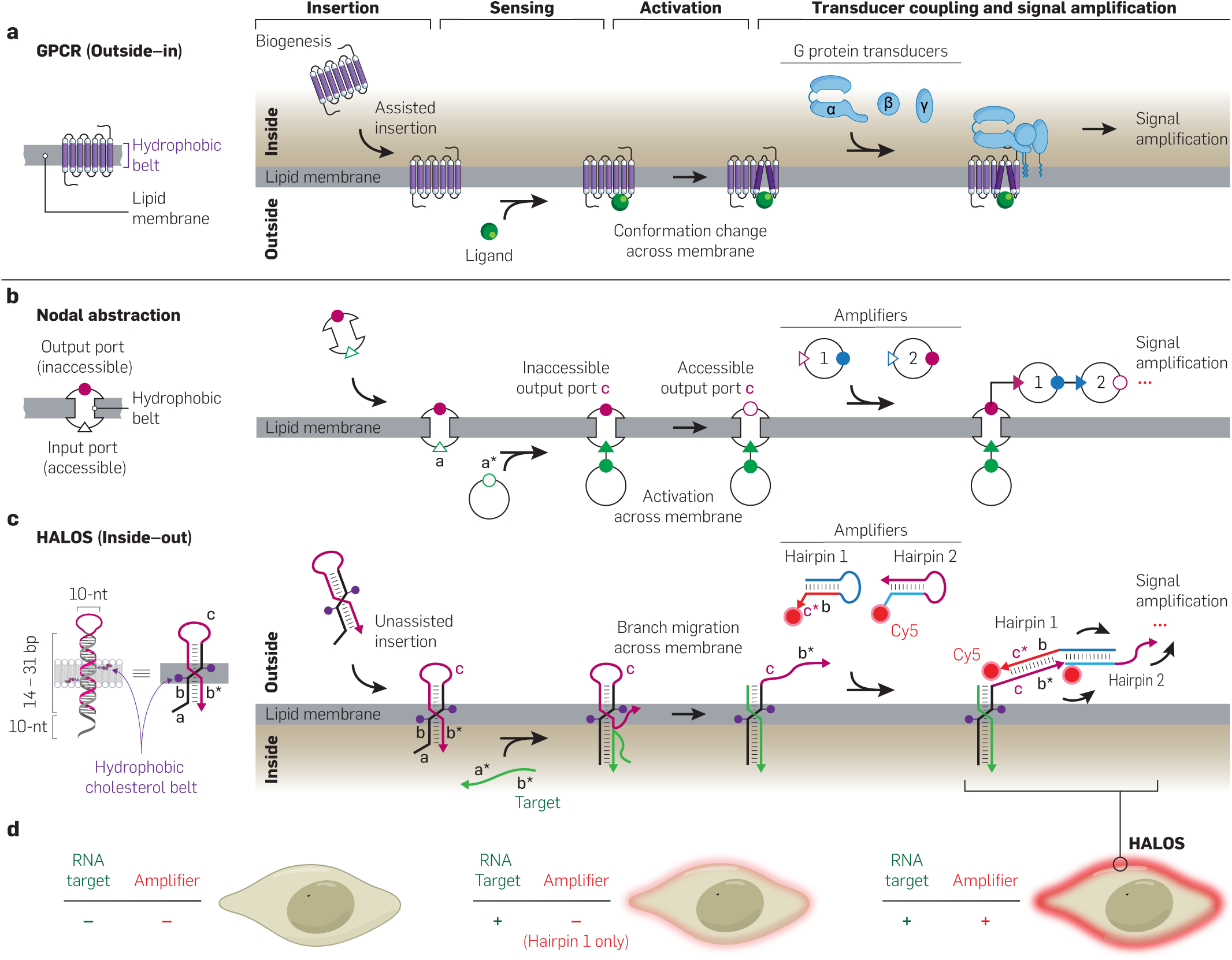
HALOS: a DNA-based synthetic transmembrane sensor that inverts natural receptor signaling to read intracellular RNA from outside the cell. HALOS recreates the input-output logic of a natural transmembrane receptor as a programmable DNA device that runs in reverse, sensing a nucleic-acid target on the intracellular face and reporting an amplified signal on the extracellular face of an intact cell. To maintain a consistent reading direction, each target species (ligand or nucleic-acid target) is placed at the bottom of the row, with signal amplification at the top. Thus, the GPCR in (**a**) is drawn with its intracellular side on top and its extracellular side at the bottom, opposite the conventional depiction. (**a**) A natural G-protein-coupled receptor (GPCR) signals *outside-in*: after biogenesis and machinery-assisted insertion, an extracellular ligand is sensed at the outer face; the resulting conformational change is relayed across the bilayer (activation) and couples an intracellular G-protein transducer (*α*, *β*, *γ*), triggering a downstream amplification cascade. (**b**) Nodal abstraction of a generic transmembrane receptor, independent of implementation: an accessible input port and an inaccessible output port, where target binding is transduced across the membrane to switch the output port to accessible, which then recruits metastable amplifiers (1, 2). Port labels a and c match the HALOS domains in (**c**). (**c**) DNA implementation, HALOS, running *inside-out*: an amphiphilic hairpin with a 10-nt toehold (a), a 14–31 bp stem (b-b*) bearing two cholesterol anchors, and a 10-nt loop (c) that inserts spontaneously. An intracellular target (a*-b*) sensed at the toehold drives branch migration of the hybridization junction across the membrane (activation), externalizing the b*-c domain, which nucleates a hybridization chain reaction between Cy5-labeled hairpin amplifiers (Hairpin 1, Hairpin 2). (**d**) Expected live-cell readout, without lysis, with the red glow reporting Cy5-labeled amplifiers recruited to the membrane: no target leaves the membrane dark; target recognition alone gives a detectable glow (+RNA target, −amplifier; H1 only); on-membrane HCR amplification makes live cells containing target RNA significantly brighter (+RNA target, +amplifier).

To summarize the signal transduction pathway, we depict the membrane-spanning HALOS reaction graph using a circular nodal abstraction consisting of a triangular input port and a circular output port (Fig. 1(b)). The state of each port is either accessible (open symbol) or inaccessible (filled symbol), reflecting whether its DNA recognition motif is available for binding. ^54^ Recognition of a complementary target nucleic acid by HALOS is represented by the input port switching from accessible to inaccessible; subsequent DNA hybridization across the membrane renders the output port accessible for reporter binding. A cascade of DNA chain reactions using metastable reporters at the accessible output port illustrates signal amplification on the membrane.

To our knowledge, HALOS is the first synthetic device to integrate sequence-specific recognition of an intracellular target, transduction of that recognition across the bilayer by DNA branch migration, and enzyme-free on-membrane amplification, without cell lysis or genetic modification (Fig. 1(c, d)). The proposed pathway initiates with cholesterol-mediated membrane anchoring in a lateral orientation, followed by insertion via axial reorientation, positioning the toehold domain for intracellular target access. Upon recognition of sequence-specific (a*-b*) Cy3-labeled targets (T-Cy3), toehold-mediated strand displacement initiates DNA hybridization and hairpin opening across the membrane, externalizing the reporter-binding domain (b*) and enabling Cy5-labeled reporter (R-Cy5) recruitment. Critically, the exposed b*-c segment can trigger a hybridization chain reaction (HCR) between Hairpin 1 and Hairpin 2, which have Cy5 labeling, to produce an amplified optical signal by creating a robust fluorescent polymer directly on the membrane surface. This integrated strategy enables the sensitive detection of intracellular targets without cell lysis.

### HALOS transmembrane stability is dictated by the stem length

Accurate transmembrane signaling by HALOS requires a hairpin stem that remains stable within the lipid bilayer, resisting hydrophobic interactions, membrane fluctuations, and base-pair disruption. Insufficient stem stability can cause spontaneous hairpin opening during membrane anchoring, prematurely exposing the b* domain and producing false-positive R-Cy5 labeling in the absence of target (Fig. 2(a)). To address this, we evaluated hairpin stems from 14 to 31 bp, each with dual mid-stem cholesterol anchors, by all-atom MD simulations (Figs. 2(b), S1 and Table S1). In these simulations, HALOS 14 broke base pairs more frequently than HALOS 31 in both aqueous and membrane environments (Fig. S1). This demonstrates that increasing the stem length from 14 to 31 bp confers greater structural integrity within the bilayer.

**Fig. 2.**
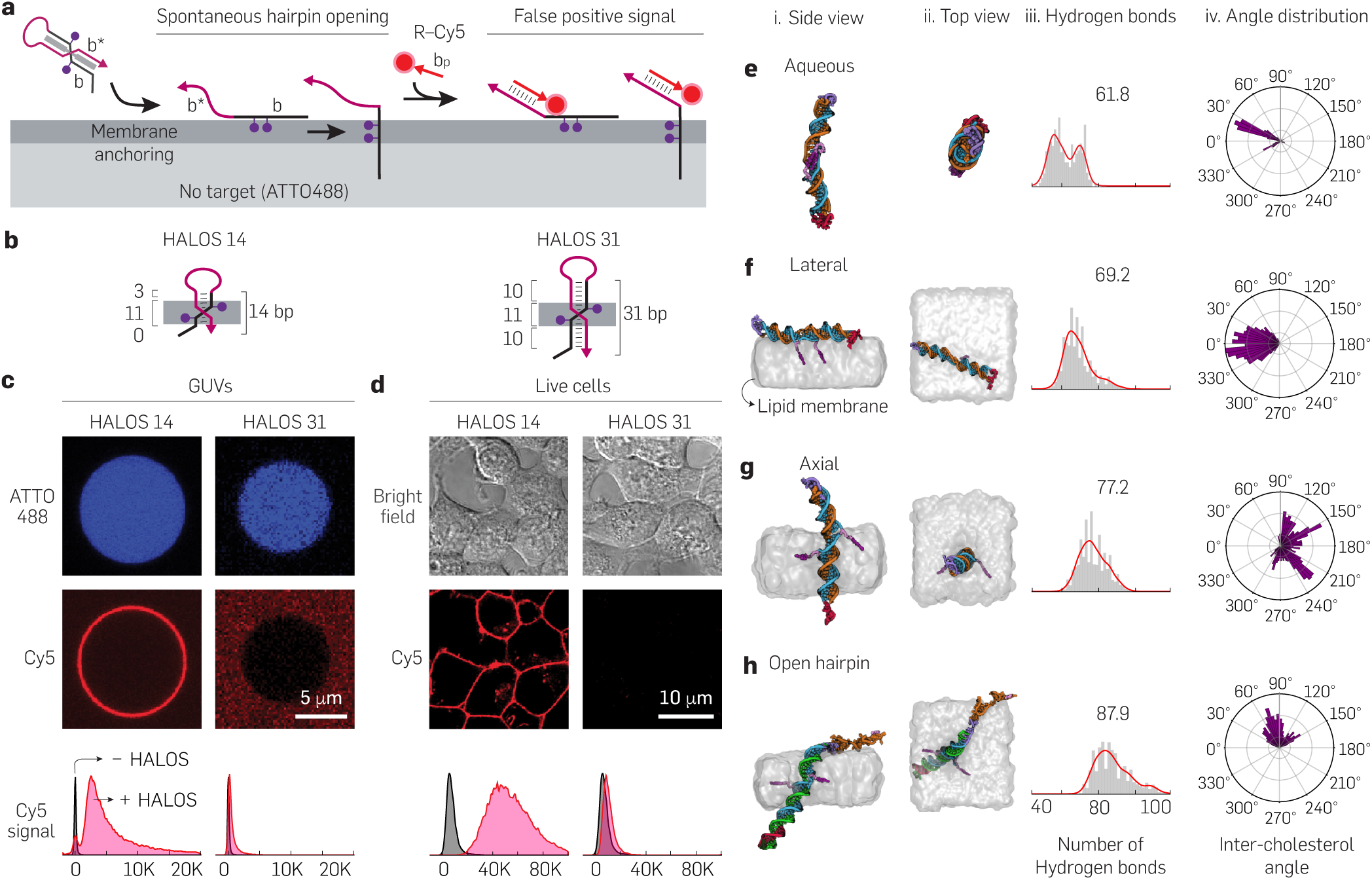
Stem-length optimization yields a stable HALOS variant that molecular dynamics predict is most stable in the membrane-inserted (axial) state. A systematic stem-length screen, corroborated by all-atom molecular dynamics, identifies HALOS 31 as the variant that resists spontaneous, target-independent hairpin opening and remains stably inserted in both synthetic vesicles and live-cell membranes. (**a**) Design rationale: an insufficiently long stem opens spontaneously during membrane anchoring, prematurely exposing the reporter-binding b* domain and producing a false, target-independent reporter signal, which motivates stem-length optimization. (**b**) The two representative constructs compared throughout, short-stemmed HALOS 14 and long-stemmed HALOS 31, shown in the axial membrane orientation. (**c** and **d**) Stem stability assays of HALOS 14 and HALOS 31 in GUVs (c; N_GUV_ ≥10,041 and N=3 independent experiments per condition) and HEK293T cells (d; N_cells_ ≥14,859 and N=3 independent experiments per condition): HALOS 31 remains dark (stable), whereas HALOS 14 produces a false reporter signal. (**e**–**h**) Side-view (i) and top-view (ii) snapshots from 1 µs all-atom MD simulations of HALOS 31 in the aqueous (**e**), lateral (**f**), axial (**g**), and open-hairpin (**h**) states, with per-state distributions of stem hydrogen bonds (iii) and the inter-cholesterol angle (iv); the membrane-inserted axial state retains the most intact base pairing among the states sampled. DNA backbone, tube; bases, spheres; lipid volume, transparent; water and ions omitted.

Stem length governed HALOS membrane stability in both synthetic giant unilamellar vesicles (GUVs) made from POPC (1-palmitoyl-2-oleoyl-glycero-3-phosphocholine) and live HEK293T cells, establishing the design parameters required for reliable cellular signaling applications. In the absence of target DNA (GUVs encapsulating only the small-molecule dye ATTO488 as a lumenal vesicle marker), we isolated the effects of stem-stability by monitoring R-Cy5 binding to the exposed b* domain on HALOS as an indicator of spontaneous hairpin destabilization (Fig. 2(a)). Retention of the lumenal ATTO488 signal throughout the assay confirmed that these control GUVs remained intact, sealed vesicles, so the R-Cy5 membrane labeling seen with short-stemmed variants reflected spontaneous hairpin opening rather than loss of vesicle integrity. We observed that HALOS constructs with 14 to 28 bp stems produced significant Cy5-positive membrane labeling on ATTO488 GUVs, diminishing from 14 to 28 bp as the stem length increased (Figs. 2(c) left and S2), suggesting that their stems are not sufficiently stable upon interaction with the membrane. Critically, HALOS 31 exhibited no detectable R-Cy5 labeling (Fig. 2(c) right), a finding corroborated by quantitative flow cytometry, which revealed negligible Cy5 fluorescence and confirmed its superior structural stability (Figs. 2(c) bottom and S2).

On live HEK293T cell membranes, HALOS variants with stem lengths of 25 bp or longer demonstrated robust stability. Cy5 fluorescence on HEK293T cell membranes revealed spontaneous destabilization exclusively in hairpin constructs with shorter 14–21 bp stems (Figs. 2(d) and S2). In contrast, HALOS variants with longer stems (25–31 bp) remained unlabeled (∼1% Cy5 positive cells), indicating a stable HALOS stem within the cell membrane. Quantitative flow cytometry was concordant with confocal microscopy results, detecting significant Cy5 signals for HALOS 14 and HALOS 21 only (*>*95% and ∼15% positive cells, respectively) on the membrane. The lower stability threshold in live cells (≥ 25 bp) compared to synthetic GUVs (≥ 31 bp) likely reflects the richer lipid and protein composition of biological plasma membranes, which may provide additional lateral support for the HALOS stem absent in simple phospholipid bilayers. Based on these comprehensive findings, HALOS 31 was selected for all subsequent studies, providing an optimized balance of structural integrity and minimal false-positive signal generation in both synthetic and cellular environments.

### Molecular dynamics reveal membrane anchoring and transmembrane stability of HALOS

All-atom MD simulations showed that HALOS 31 adopts a stable axial orientation within the lipid bilayer. In the target-bound, open conformation, the non-cholesterol stem (b*) resides as a free segment in the outer aqueous solution, consistent with its role in signal transduction. We systematically compared HALOS stability across various states, including aqueous, lateral, axial, intermediate, and open conformations, using hydrogen-bond analysis and quantified the angle between the two cholesterol anchors (Figs. 2(e–h) and S3). Side and top views from post-MD simulation snapshots demonstrated that HALOS 31 stably resided within the lipid bilayer while maintaining overall conformational integrity. Over 1 µs of simulation, the DNA hairpin exhibited substantial structural fluctuations, primarily in the flexible loop and single-stranded toehold domains. Notably, without the lipid bilayer, the double-stranded stem of HALOS remained mostly stable; however, the cholesterol moieties tended to stack and intercalate between DNA bases to avoid the aqueous environment, resulting in significant helical distortion (Movie S1).

Quantitative analysis of base-pair stability indicated that membrane incorporation reduces base-pair breaks, with the axial orientation the most stable of the states sampled. The time evolution of broken base pairs (bp), defined by hydrogen bond thresholds (A-T: *<*2 H-bonds; C-G: *<*3 H-bonds; distance *<*0.34 nm, angle *<*45*^◦^*), showed minimal disruption in axially anchored hairpins, while HALOS in aqueous solution exhibited increased base-pair breaks near the end of the simulation (Fig. S1). Hydrogen bond analysis showed the most intact base pairing in the axial state (mean±S.D.= 77.2±7.2 H-bonds), versus the lateral (69.2±7.5) and aqueous (61.8±7.3) states (Fig. 2(e–g)). The contour length distributions further indicated that laterally oriented or aqueous HALOS states were overstretched around hydrophobic modifications compared to the axial conformation (Fig. S3). Consistently, the root mean square fluctuations (RMSF) data showed that membrane incorporation stabilized the overall HALOS structure, whereas hairpin stems in aqueous solution experienced substantial conformational fluctuations (Fig. S3).

Since the cholesterol anchors are placed 180*^◦^* apart in HALOS design, the predicted angle between cholesterol anchors during MD simulations critically determines stem integrity and HALOS orientation, directly impacting target recognition function. In aqueous solution and in laterally oriented HALOS conformations, the inter-cholesterol angle is small (0–30*^◦^*), associated with cholesterol stacking, stem distortion, and an unfavorable helical conformation (Fig. 2(e–f) and Movie S1). Conversely, a large angle (100–210*^◦^*) promotes stable axial orientation, resulting in increased stability and positioning HALOS perpendicular to the membrane, thereby facilitating target recognition (Fig. 2(g) and Movie S1). MD simulations of target-bound HALOS in intermediate and open hairpin conformations confirmed the feasibility of signal transduction by membrane-anchored HALOS. The increased number of hydrogen bonds (87.9±9.5) compared to HALOS alone and a distinct large angle distribution between cholesterol anchors (60–150*^◦^*) confirm the stable residence of these constructs within the membrane (Figs. 2(h), S3, and Movie S2). These conformational changes, characterized by the altered H-bond network and cholesterol anchor angle, facilitate the exposure of the b* arm to the outer aqueous solution, enabling R-Cy5 recruitment on the opened HALOS domain, a configuration consistent with signal transduction across the membrane. We emphasize that these microsecond trajectories probe the conformational stability of each pre-assembled state (aqueous, lateral, axial, and target-bound) rather than the much slower kinetics of membrane insertion and transmembrane branch migration, which proceed over seconds to minutes in our experiments. The simulations therefore establish that the inserted and target-bound states are structurally stable once formed, while their formation and interconversion are demonstrated directly by the imaging and reporter assays (Figs. 3–7).

**Fig. 3.**
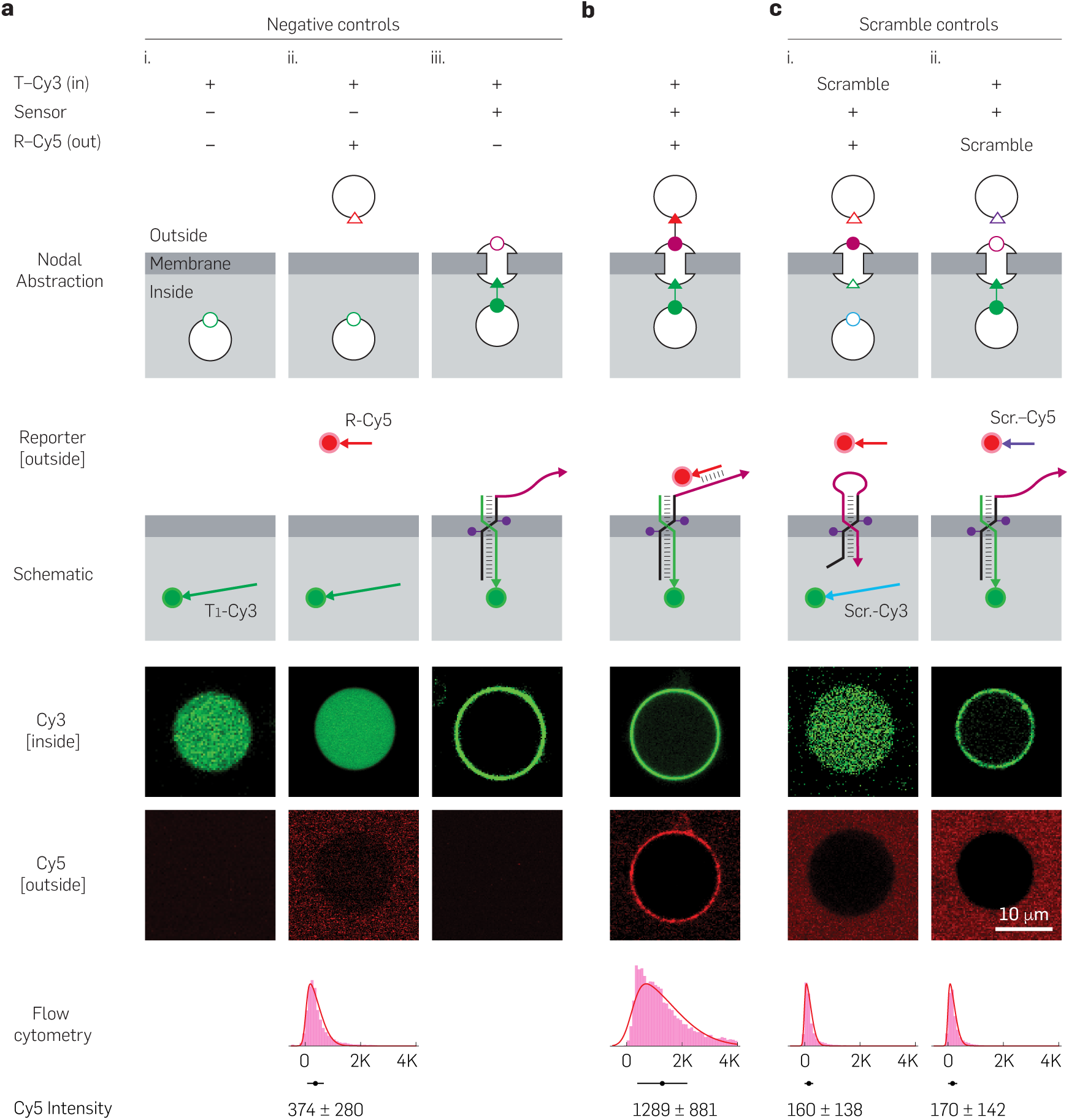
HALOS 31-mediated specific signal transduction in GUVs. Target detection by membrane-embedded HALOS 31 triggers a conformational switch that exposes the hairpin domain for reporter labeling, transducing an internal signal to the GUV exterior. (**a**) Negative controls establishing that signal transduction requires all components (nodal abstractions and schematics, top rows). Three conditions (i–iii) each omit one element: (i) target only, (ii) target and reporter without sensor, and (iii) target and HALOS 31 sensor without reporter. In the Cy3 (inside) channel, T-Cy3 remains in the GUV lumen when the sensor is absent (i, ii) and forms a membrane Cy3 ring only when HALOS 31 is present (iii); the Cy5 (outside) channel stays dark in all three, with mean Cy5 fluorescence of 374±280 a.u. (N_GUV_=22,663). (**b**) Positive validation (nodal abstraction and schematic): target binding triggers the HALOS 31 conformational change, exposing the open hairpin domain (b*) for recognition by a single 13-nt ssDNA reporter (R-Cy5); no chain reaction occurs here, as enzyme-free HCR amplification is introduced later (Fig. 6). Co-localized Cy3 and Cy5 signals and elevated Cy5 fluorescence (1,289±881 a.u., N_GUV_=16,669) confirm efficient transmembrane signal transfer. (**c**) Sequence-specificity controls: a scrambled target gives no Cy3 or Cy5, and a scrambled reporter binds neither despite correct target recognition (160±138 and 170±142 a.u.; N_GUV_=3,957 and 18,223, respectively). The transmembrane branch-migration step drawn in the schematics is validated independently in Fig. 7. N=3 independent experiments per condition. Error bars are ±SD.

**Fig. 4.**
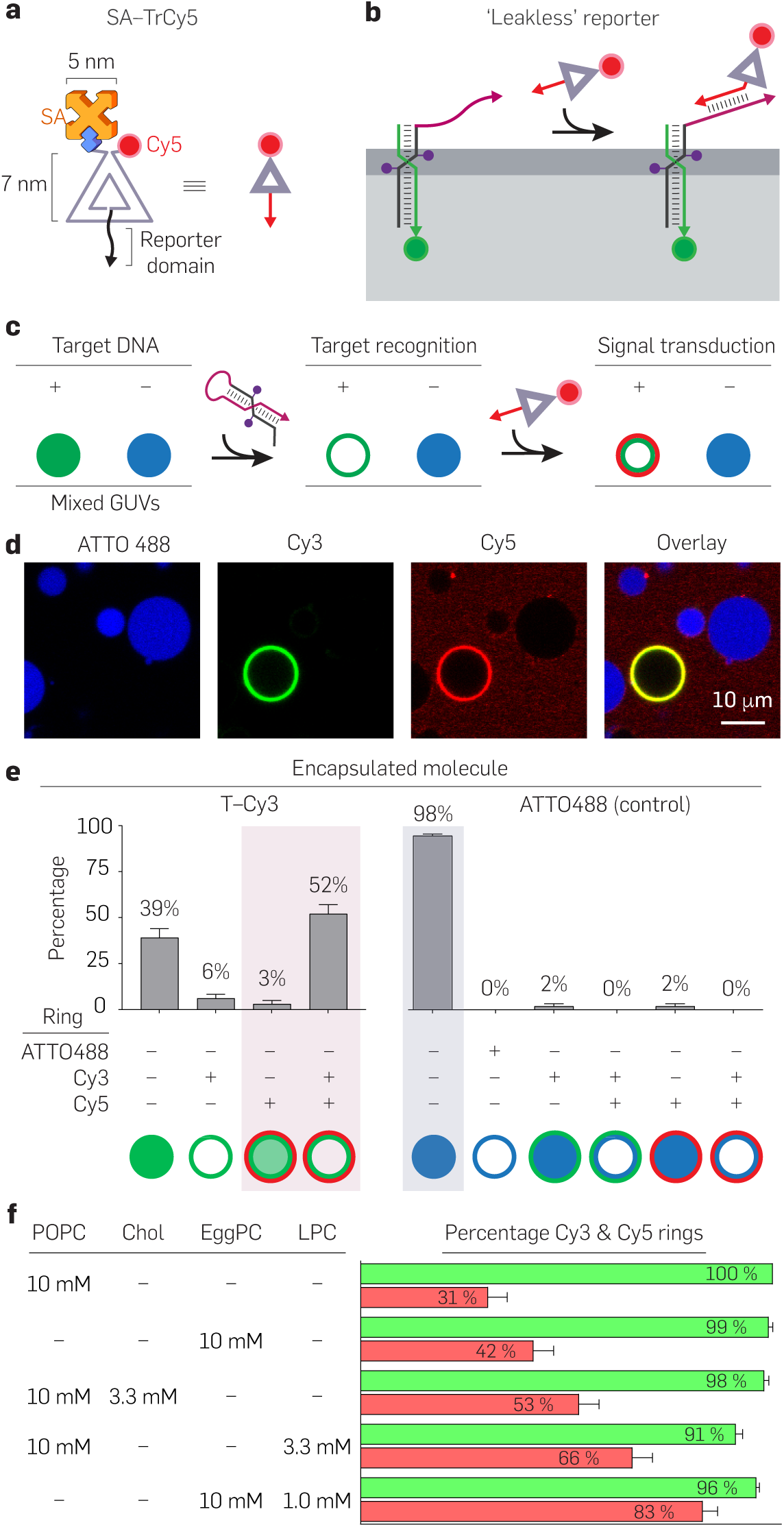
Leakless, target-specific and membrane-optimized signal transduction in GUVs. A bulky triangular reporter confines signal transduction to the outer membrane, enabling target-specific detection whose efficiency scales with membrane fluidity. (**a**) Triangular reporter design: 2 complementary DNA strands forming a 7-nm triangle bearing Cy5 and biotin labels, further enlarged by streptavidin (SA; 5 nm) to suppress leakage. (**b**) Bulky SA-TrCy5 reporter labeling target-recognized HALOS 31, preventing leakage across the membrane. (**c**) Schematic of the mixed-population assay: target-encapsulated GUVs combined with ATTO488 control GUVs, with signal transduction reported by SA-TrCy5. (**d**) Selective signaling from T-Cy3-encapsulated GUVs, showing membrane labeling, while ATTO488-GUV controls lacked Cy3 and Cy5 signals. (**e**) Quantitative detection analysis of T-Cy3 GUVs (N_GUV_=351). Signal transduction (Cy5-positive) occurred in 52±5% of GUVs with full and a further 3±2% with partial Cy3 recognition; 6±2% showed recognition without transduction (Cy3-positive, Cy5-negative); and 39±5% showed no recognition. Control ATTO488 GUVs (N_GUV_=420) showed 2±1.5% Cy3/Cy5 labeling, indicating negligible non-specific activation. (**f**) Transmembrane signaling efficiency varied with lipid composition, ranging from 31±4.6% in POPC to 83±3.6% in EggPC+LPC, with EggPC (42±4.9%), POPC+Chol (53±4.9%), and POPC+LPC (66±4.8%) intermediate. Error bars are ±SD from bootstrapping; N_GUV_ ≥846 and N=5 independent experiments per condition.

**Fig. 5.**
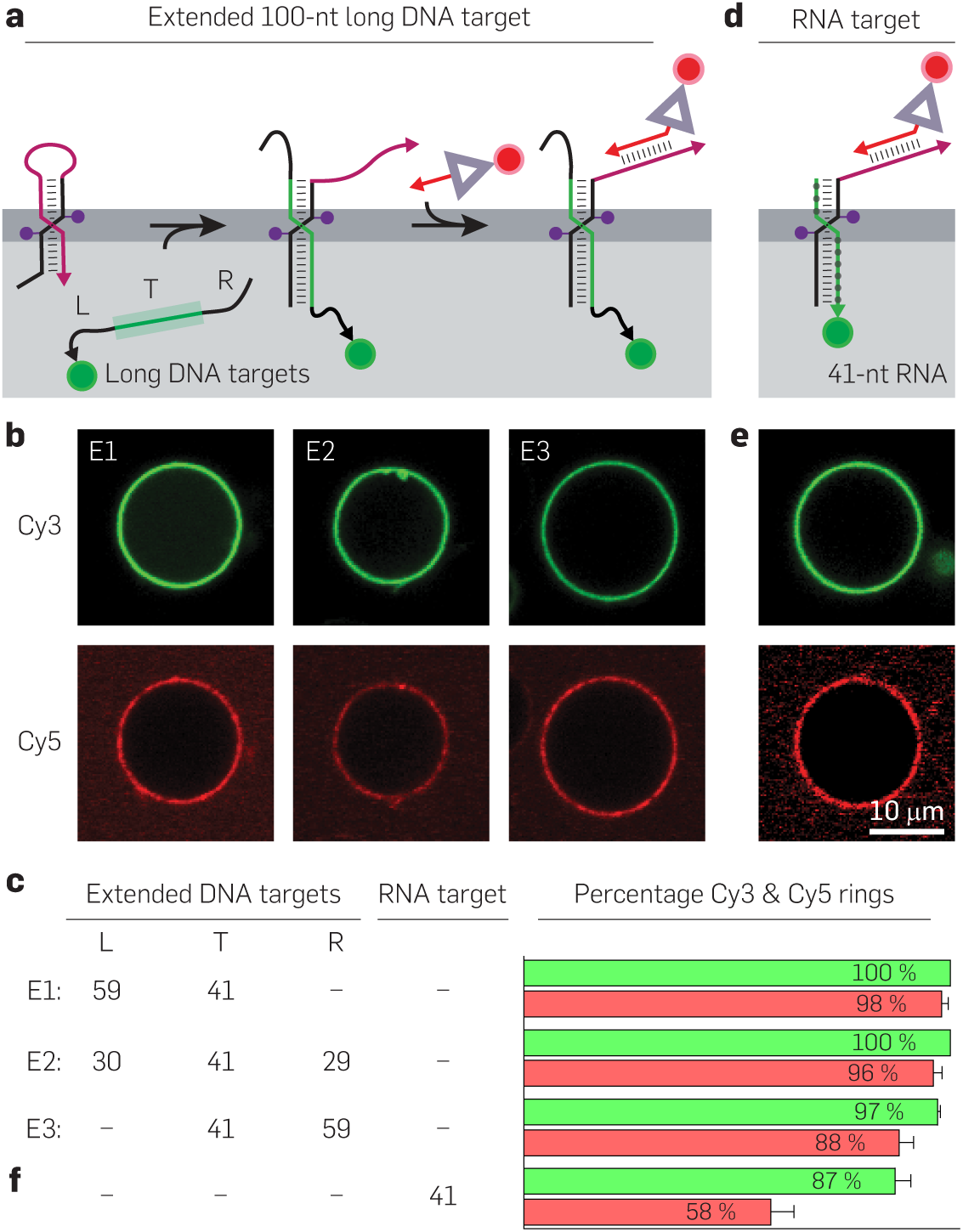
Transmembrane signal transduction upon detection of extended 100-nt DNA and modified RNA targets. HALOS 31 detects both extended 100-nt DNA and 2′-OMe RNA targets and recognizes its target site regardless of where it lies within a longer strand, extending sequence-programmable transmembrane signal transduction beyond short DNA. (**a**) Schematic illustrating the detection of 100-nt long DNA targets encapsulated within GUVs, with signal transduction mediated by SA-TrCy5 labeling. (**b**) Confocal images revealing Cy3 and Cy5 membrane labeling for HALOS-mediated target recognition and signal transduction, respectively. (**c**) Quantitative analysis displaying efficient signal transduction for three variants of the 100-nt DNA target with the 41-nt recognition domain at the 5′ end (E1), middle (E2), or 3′ end (E3): E1 (98±1.4%, N_GUV_=121), E2 (96±2%, N_GUV_=113), E3 (88±3.2%, N_GUV_=115). (**d**–**f**) Schematic representation (d), confocal images (e), and quantitative analysis (f) of 41-nt 2′-OMe modified RNA target detection (Cy3; 87±3.6%) and signal transduction (Cy5; 58±5.3%, N_GUV_=311), confirming HALOS 31-mediated detection of diverse encapsulated nucleic acid targets. Error bars are ±SD computed from bootstrapping analysis. N=3 independent experiments per condition.

**Fig. 6.**
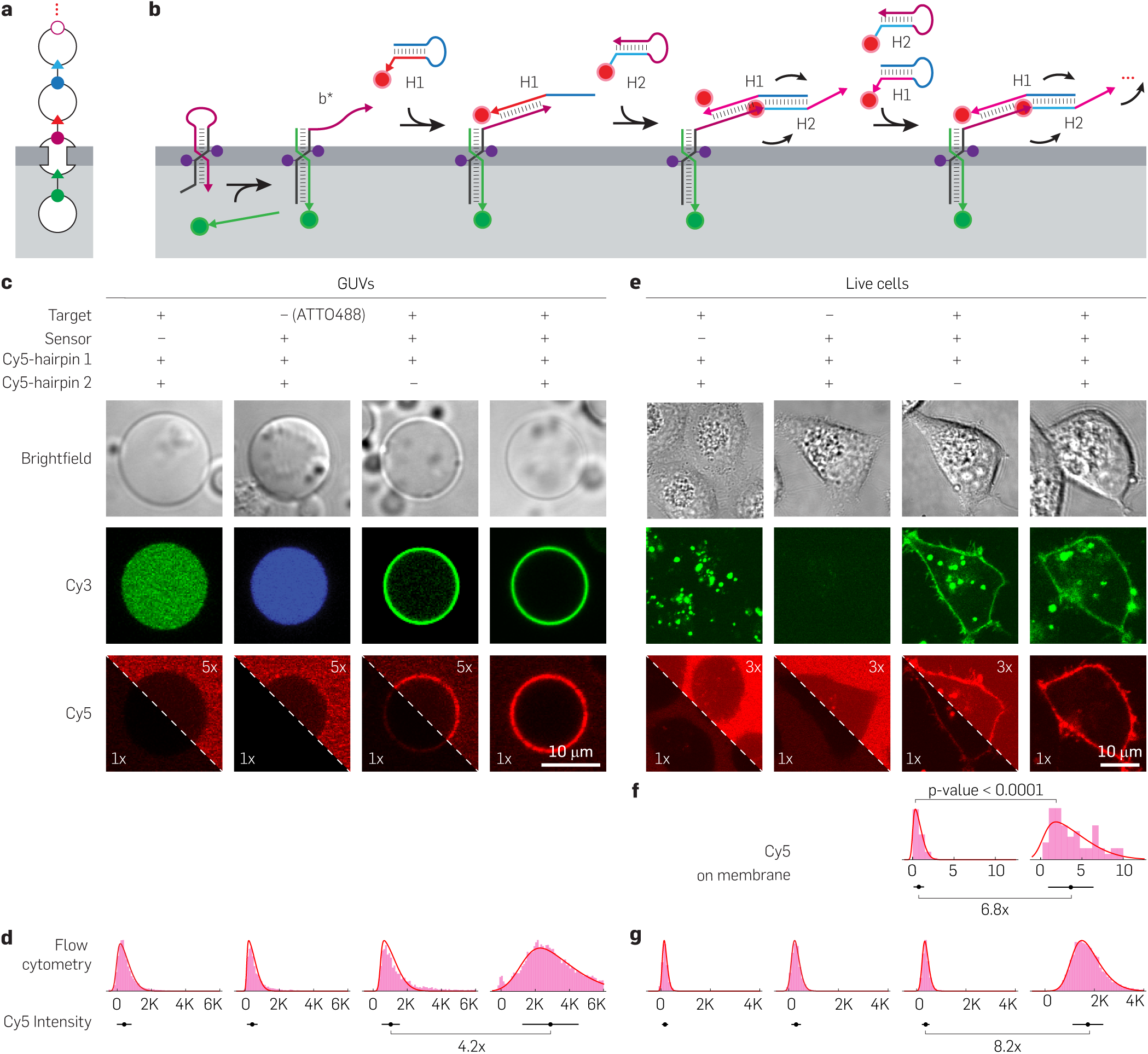
HCR on membrane facilitating background-corrected 8.2× enhancement over a single hairpin reporter in HALOS-mediated transmembrane signaling. Target-recognized HALOS initiates membrane-localized hybridization chain reaction (HCR) for amplified RNA detection in giant unilamellar vesicles (GUVs) and live cells. (**a**) Abstraction graph of target-recognized HALOS initiating extracellular HCR using two metastable hairpins (H1 and H2) for signal amplification. (**b**) Schematic of HALOS-mediated target recognition, generating an initiator for HCR on membranes, triggering stepwise fluorescence signal amplification. (**c**) Confocal images of GUV membranes (Cy5 channel): no Cy5 signal in controls lacking HALOS 31 or targets, minimal signal with hairpin 1 alone, and strong amplification with both hairpins. (**d**) Quantification reveals background-corrected ∼4.2× amplification in GUVs: background without HALOS 31 (450±394 a.u., N_GUV_=14,020) or targets (453±279 a.u., N_GUV_=3,623), detectable signal with H1 (1,058±491 a.u., N_GUV_=3,663), and significantly higher with HCR amplification (3,017±1,591 a.u., N_GUV_=14,984). (**e,f**) Live A549 cells detecting a synthetic RNA target with HCR amplification. Cy3 (RNA target recognition) and Cy5 (HCR amplification) membrane signals show background-corrected ∼6.8× enhancement over H1 alone (*P <* 0.0001, N_cells_ ≥55). Controls lacking HALOS 31 or targets show no membrane labeling. (**g**) Quantification on live HEK293T cell membranes confirms background-corrected ∼8.2× signal amplification: minimal background without HALOS 31 (168±103 a.u., N_cells_=8,320), without targets (296±171 a.u., N_cells_=11,124), moderate signal with hairpin 1 (360±134 a.u., N_cells_=7,402), and significantly higher with HCR (1,739±598 a.u., N_cells_=8,946). Data are mean±SD; N=3 independent experiments per condition; annotated P values were determined using the two-sided Mann–Whitney *U* test.

**Fig. 7.**
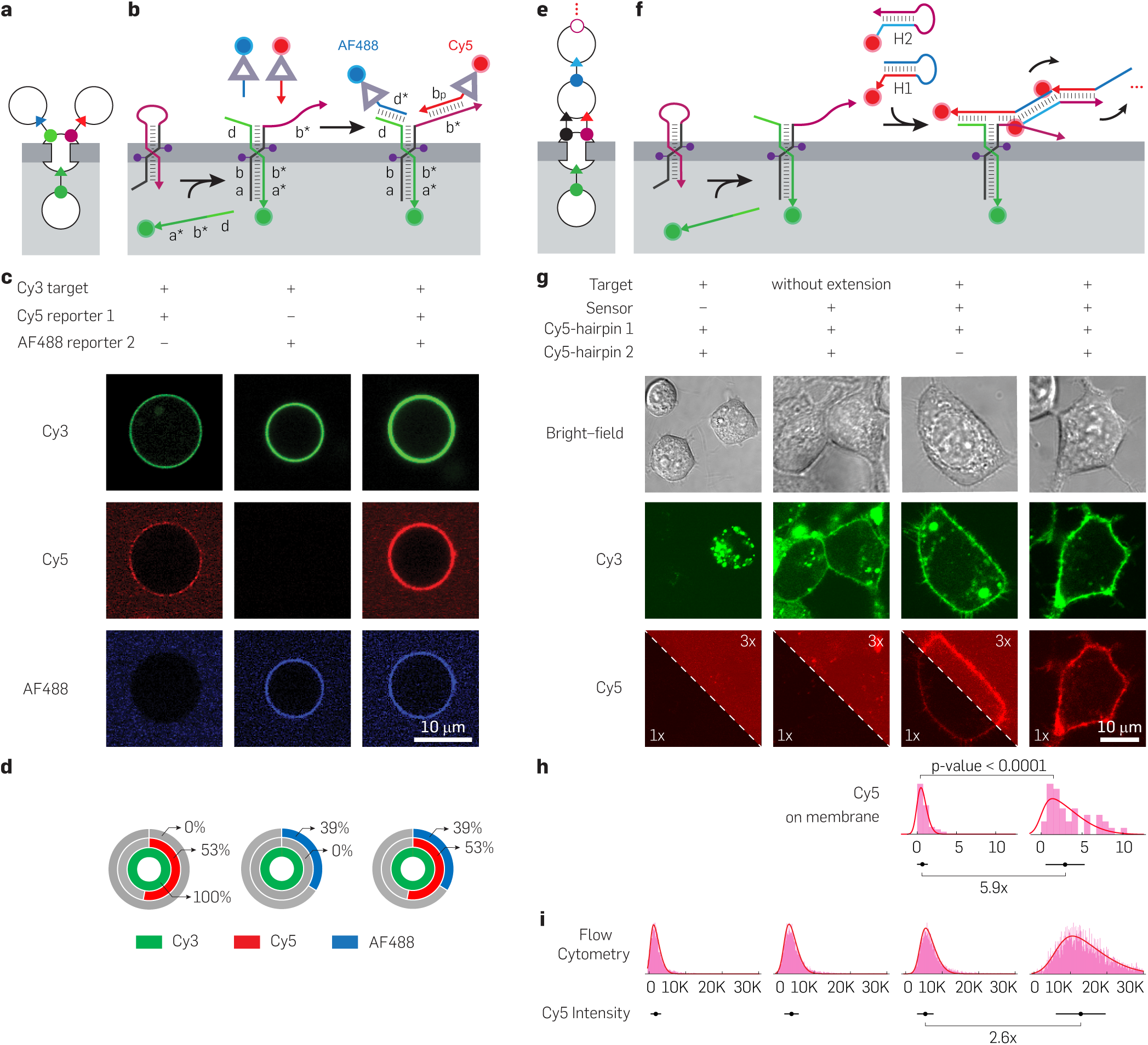
Mechanistic insights into DNA branch migration across membranes by dual reporter labeling and Proximity Split-Initiator HCR (ψ-HCR). Two orthogonal strategies, dual reporter labeling and proximity-based ψ-HCR, provide direct evidence for transmembrane DNA branch migration on GUVs and live cells. (**a**) Nodal abstraction of dual reporter labeling (SA-TrCy5, red; SA-TrAF488, blue) to T′-Cy3-recognized HALOS 31 on the membrane. (**b**) Schematic of T′-Cy3 detection by HALOS 31, branch migration of extended targets across the membrane, and orthogonal dual reporter labeling of the extended target domain (d) and HALOS open hairpin domain (b*). (**c**) Individual and simultaneous dual labeling by SA-TrCy5 (red) and SA-TrAF488 (blue), confirming the absence of cross-reactivity. (**d**) Quantitative dual-reporter labeling on T′-Cy3-recognized GUVs (N_GUV_=652): SA-TrCy5 (open hairpin, b*) labeled 53±5.0% and SA-TrAF488 (branch-migrated target domain, d) labeled 39±4.9%, the same whether applied individually or together. (**e**) Nodal abstraction of ψ-HCR on the membrane by T′-Cy3-recognized HALOS 31. (**f**) Target extension and hairpin opening of HALOS 31 trigger formation of the proximal split initiator, enabling ψ-HCR amplification using H1 and H2. (**g,h**) ψ-HCR-mediated signal enhancement on live A549 cell membranes (g); controls lacking HALOS 31 or the extended target domain showed no membrane Cy5 signal, whereas the extended target gave background-corrected 5.9× amplification over H1 alone (h, *P <* 0.0001, N_cells_ ≥39, N=3 independent experiments per condition). (**i**) Live HEK293T cell membranes: ψ-HCR (14,000±7,000 a.u.) shows a background-corrected 2.6× amplification over H1 alone (6,300±2,117 a.u.), with backgrounds measured without HALOS 31 (1,800±1,300 a.u.) or target (2,900±1,800 a.u.). N_cells_ ≥7,725 and N=3 independent experiments per condition. Error bars are ±SD.

### HALOS 31 enables specific transmembrane target detection and signal transduction in GUVs

Following the evaluation of HALOS 31 stability on the lipid membrane, we assessed its efficacy as a transmembrane sensor for target detection on GUVs. NUPACK design predictions ^55^ and gel electrophoresis confirmed the specificity of target recognition, hairpin opening, and subsequent reporter binding by HALOS 31 in solution (Fig. S4). To test membrane-embedded detection, T-Cy3 (41-nt) was encapsulated within POPC GUVs via electroformation, ^56^ validated by the formation of lipid spheres containing Cy3 dyes (Fig. 3(a) left). To stringently suppress false-positive signals from T-Cy3 outside the GUVs, non-encapsulated T-Cy3 was removed by repeated washing, and any residual external target was quenched with a ∼20× molar excess of complementary DNA (see Methods; Fig. S5). Upon incubating these GUVs with HALOS 31, pronounced Cy3 labeling appeared on the membranes after only 5 min of incubation, indicating efficient membrane insertion and target recognition (Figs. 3(a) right and S5). A progressive decrease in lumenal Cy3 fluorescence and a corresponding increase on the membrane indicated spatial redistribution of T-Cy3 targets from lumen to membrane, consistent with HALOS 31-mediated target capture. Quantification of sensor activity showed that 86±1.9% GUVs exhibited distinct Cy3-rings within 90–120 min (Fig. S5). Analysis of target recognition kinetics demonstrated heterogeneity in the rate of Cy3 accumulation at the membrane periphery over 120 mins (Fig. S6), likely reflecting variable lipid rafts (monolayer *vs.* multilayer), target encapsulation efficiencies based on GUV size variations, and HALOS 31 population on individual GUVs. Notably, no target detection was observed in either of the two controls: GUVs lacking HALOS 31 entirely, and GUVs treated with cholesterol-free HALOS variants (Figs. 3(a) middle and S7). Both controls confirm that membrane anchoring via cholesterol and the full HALOS 31 construct are each required for transmembrane signaling.

To assess the signal transduction capability of HALOS 31, we triggered its conformational switch with T-Cy3 detection (green), enabling R-Cy5 labeling (red) of the exposed extracellular domain (b*) from the outside (Fig. 3(b)). Confocal microscopy showed both Cy3 and Cy5 labeling only in GUVs containing HALOS 31, with controls lacking HALOS 31 and T-Cy3 displaying no Cy5 signal (Figs. 3(a–b), 2(c) and S7), indicating successful transmembrane signaling. Quantitative flow cytometry analysis demonstrated a significant increase in mean Cy5 fluorescence on GUVs containing HALOS 31 (1,289±881 a.u.) compared to its controls (374±280), confirming specific transmembrane signal transduction (Fig. 3(a–b) bottom). Consistently, Cy3 *vs.* Cy5 intensity plots showed that 26.9% of T-Cy3 GUVs with HALOS 31 displayed overlapping membrane labeling, compared to 1.8% without HALOS 31 (Fig. S7), highlighting the specificity of the signal transduction.

To demonstrate the reverse, an outside-in direction akin to canonical GPCR signaling, we performed an analogous assay on GUVs. Because HALOS 31 likely inserts in both orientations (toehold-in and toehold-out) during membrane spanning, a fraction should remain accessible for external target recognition. Upon externally added T-Cy3 binding, HALOS 31 opening was initiated from the outside, resulting in hybridization across the membrane to the inside of the GUVs, as evidenced by successful labeling with pre-encapsulated R-Cy5 GUVs (Fig. S8). No signal transduction was observed in the absence of external T-Cy3.

HALOS 31 exhibited high target selectivity, as scrambled DNA targets (Scr. T-Cy3) did not induce membrane labeling with Cy3 and Cy5 (Figs. 3(c) left and S9). Similarly, to assess the specificity of R-Cy5 labeling, we tested scrambled reporter sequences (Scr. R-Cy5) on GUVs in which HALOS 31 was correctly opened by its target and observed no non-specific Cy5 membrane labeling by confocal microscopy (Figs. 3(c) right and S9). Flow cytometry further confirmed negligible Cy5 labeling with both Scr. T-Cy3 (0.6%) and Scr. R-Cy5 (0.3%; Figs. 3(c) below and S9). These results collectively establish HALOS 31 as a robust and specific transmembrane sensor.

### Next-generation bulkier reporter design enables “leakless” signal transduction across the membrane

While monitoring HALOS-mediated signal transduction across synthetic GUVs (Fig. 3(b)), R-Cy5 leakage was observed in a minor population (*<*30%) of GUVs (Fig. S7). This leakage likely resulted from transient nanopore formation by DNA sensor duplexes with dual cholesterol anchors, ^36,48,57^ allowing 13-nt ssDNA R-Cy5 (radius of gyration ∼1.7 nm, supplementary methods) to diffuse through the membrane during the conformational change. This diffusion could lead to false positives by labeling partially anchored T-Cy3 within the GUV inner lumen (Fig. S10). Extending the R-Cy5 ssDNA strand length to 31-nt reduced the leakage but did not completely eliminate this undesired effect.

By engineering a reporter too bulky to cross the membrane, we achieved “leakless” signal transduction: the reporter remained confined to the outer membrane, with minimal diffusion across the bilayer into the vesicle lumen and hence minimal false-positive internal labeling. This reporter is a biotinylated, triangle-shaped DNA structure (Tr, ∼7 nm edge) bearing a 21-nt labeling domain (Figs. 4(a) and S10), large enough to be blocked from any such transient pores (Fig. 4(b)). Gel electrophoresis confirmed Tr formation and its specific labeling of T-Cy3-bound HALOS 31 in solution (Fig. S10). Loading the triangle with streptavidin (SA-TrCy5) enlarged it further and markedly reduced reporter leakage into the GUVs (Figs. 4(c–d) and S10); the ∼5 nm SA sterically obstructs the escape of any misfolded TrCy5 fragments.

Lysis-free detection of specific intracellular nucleic acid targets remains a significant challenge in biological research. To enable this, we performed signal transduction using a mixed population of T-Cy3 encapsulated and non-target control (only ATTO488 dyes) GUVs (Fig. 4(c)). Confocal imaging revealed membrane labeling by SA-TrCy5 exclusively on T-Cy3 GUVs (Figs. 4(d) and S10), confirming the high specificity of the signal transduction. Quantitative analysis demonstrated that either complete (52±5%) or partial (3±2%) T-Cy3 detected GUVs showed signal transduction via membrane Cy5 labeling. In the remaining GUVs, two scenarios were observed: some GUVs filled with Cy3 did not exhibit T-Cy3 recognition or signal transduction (39±5%), while others were recognized by T-Cy3 but showed no Cy5 labeling (6±2%, Fig. 4(e) left), possibly due to unfavorable HALOS orientation, steric occlusion of the b* domain, or incomplete hairpin opening within the experimental timeframe. Furthermore, minimal Cy3 and Cy5 labeling (2±1.5%) on control GUVs (ATTO488) ruled out false positives from external quenched targets (Fig. 4(d–e) right). The minimal false-positive signal on the control GUVs was attributed to sample handling and GUV fusion during probe incubation.

### Optimized membrane fluidity enhances HALOS-mediated signal transduction

Membrane fluidity, a critical physical property influenced by lipid composition, significantly impacts the function of membrane-associated nanodevices, particularly in the insertion and migration of charged DNA species. Using GUVs composed of POPC, a major mammalian membrane component, and the more biologically relevant EggPC, we systematically tuned the membrane fluidity by varying the core components.^58,59^ Cholesterol increases membrane order and rigidity, while 18:1 LysoPC (LPC) enhances fluidity and curvature, known to influence ion channel activity and signaling. ^58,60^ All-atom MD simulations revealed that the lipid compositional changes significantly altered the lipid bilayer thickness, head group density, lipid order parameters, and area per lipid, suggesting that the introduction of LPC increases the fluidity of the phospholipid membrane (Fig. S11). Experimentally, Laurdan generalized polarization (GP) calculation ^61^ and imaging demonstrated a significant alteration in membrane order and fluidity induced by cholesterol and LPC (Fig. S12).^62^ Specifically, 30% cholesterol in POPC was highly ordered (GP *>* 0.4), whereas adding LPC to POPC and EggPC demonstrated a highly disordered and fluidic membrane (GP *<* −0.15). Systematic studies on GUVs have revealed that HALOS-mediated signal transduction is dependent on the membrane liquid phase (ordered *vs.* disordered), which reflects its fluidity. Increased fluidity leads to increased lipid disorder, which boosts water penetration and the movement of amphiphilic molecules within the bilayer, thereby aiding DNA hybridization and effective transmembrane signaling. ^63^ The dynamic phase found in LPC-rich membranes improves lateral lipid diffusion and can lower the free energy barrier for DNA hybridization, allowing for swift signal transduction. Increasing fluidity, particularly via LPC addition, boosted signal transduction by SA-TrCy5 labeling from 31±4.6% in POPC-only to 66±4.8% in 30% (mol/mol) LPC in POPC GUVs, whereas it increased from 42±4.9% in EggPC to 83±3.6% in 10% (mol/mol) LPC in EggPC GUVs (Figs. 4(f) and S12). Cholesterol-enriched POPC showed intermediate efficiencies (53±4.9%). Notably, addition of HALOS 31 (250 nM) did not alter intrinsic membrane order across compositions (Fig. S12), confirming that signal enhancement derives from changes in lipid environment rather than sensor effects. These data establish membrane fluidity as a major but not the sole determinant of HALOS-mediated transduction efficiency, since the more ordered cholesterol-enriched POPC (POPC+Chol) supported higher transduction (53%) than the more fluid EggPC (42%) and POPC (31%) membranes.

### HALOS 31 enables detection and transmembrane signaling of longer nucleic acid targets

HALOS 31 efficiently detects long nucleic acid targets. Long cytosolic RNAs are critical disease biomarkers, but their detection in live cells without lysis or complex delivery methods remains challenging. Systematic validation using GUVs encapsulating synthetic 100-nt ssDNA with a target domain (41-nt) placed either at one of the termini or in the middle of the ssDNA demonstrated consistent ≥97% detection (Cy3) and 88–98% signal transduction (Cy5) efficiency across multiple target positions (Figs. 5(a–c) and S13). Detection remained essentially position-independent, with only a modest position dependence in signal transduction, confirming that HALOS reliably detects target sequences regardless of their position within a longer ssDNA molecule. The decreasing trend of signal transduction efficiency from E1 to E3 might be due to the ineffective DNA hybridization across the membrane by extended DNA domains from the 5′ end of the targets following HALOS opening. These results establish HALOS 31 as a proof-of-concept for lysis-free detection of longer nucleic acid targets, a step toward live-cell biomarker diagnostics.

Thermodynamically, an RNA target is at least as favored as a DNA target to invade and open the HALOS hairpin, because RNA-DNA hybrids are more stable than the corresponding DNA-DNA duplexes (RNA-RNA *>* RNA-DNA *>* DNA-DNA). To validate HALOS-mediated transmembrane signaling upon RNA-target recognition, we encapsulated a 2′-O-methyl RNA target (OMe-T-Cy3, 41 nt), a synthetic non-coding RNA, within GUVs and demonstrated successful HALOS 31-mediated detection with 87±3.6% efficiency (Figs. 5(d–f) and S13). Signal transduction efficiency of 58±5.3% confirms HALOS-mediated RNA detection; the lower efficiency relative to DNA targets likely reflects kinetic and geometric factors, such as the A-form geometry of RNA-DNA hybrids and slower branch migration, rather than reduced hybrid stability. This proof-of-concept for lysis-free RNA-target detection is a step toward sensing cellular RNAs, such as viral RNAs and mRNA biomarkers, in living cells.

### Membrane-localized HCR amplifies HALOS signaling up to 8.2-fold over a single hairpin reporter for lysis-free non-coding RNA detection in living cells

HALOS 31 enables membrane-localized HCR ^50,51^ for lysis-free RNA detection in living cells. The exposed b* domain of target-recognized HALOS 31 serves as an HCR initiator for enzyme-free signal amplification on the membrane, depicted as nodal-abstraction reaction graphs (Fig. 6(a–b)). Systematic validation using NUPACK and gel electrophoresis verified specific HCR product formation only when both metastable hairpin amplifiers were present (H1, H2), establishing target-dependent amplification (Fig. S14). The results showed a higher-order HCR assembly in solution exclusively in the presence of target-detected HALOS 31, with no amplification products detected in the negative controls.

Membrane-localized HCR produced robust, target-dependent signal amplification in GUVs. Confocal microscopy revealed Cy5 amplification exclusively in target-containing GUVs, with no signal detected in the negative controls lacking either HALOS 31 or T-Cy3 (Figs. 6(c) and S15). Comparing the HCR assembly images to those of H1 alone revealed a significant increase in the membrane Cy5 intensity. Quantitative flow cytometry demonstrated significant signal enhancement when both H1 and H2 monomers were present (3,017±1,591) compared to H1 alone (1,058±491), confirming the HCR efficiency on the membrane (Fig. 6(d)). The large standard deviations reflect the heterogeneity in the GUV size and target encapsulation efficiency, which is consistent with the single-vesicle measurements. After background subtraction using negative controls, without HALOS 31 (450±394), the net amplification was calculated to be ∼4.2× (2,567 *vs.* 608 a.u.). Three-dimensional imaging confirmed the spatial confinement of HCR products on the GUV peripheral membrane, establishing membrane-localized amplification architecture (Fig. S15). These results demonstrate that HALOS 31-initiated HCR overcomes the sensitivity limitations of single-reporter-based detection while maintaining high specificity for encapsulated targets.

Next, we extended HCR amplification to living cells, enabling lysis-free detection of intracellular synthetic non-coding RNA. Prior to detecting intracellular nucleic acid targets, the live-cell stability and compatibility of HALOS constructs were evaluated by performing signal transduction on live HEK293T cell membranes (Fig. S16). While the detection of endogenous mRNA requires target-specific HALOS redesign and extensive optimization, we validated transmembrane signaling and amplification using synthetic non-coding 2′-O-methyl RNA targets (2′-OMe-T-Cy3, 100 nM) as proof-of-concept. Lipofectamine-mediated delivery of synthetic RNA targets into live A549 and HEK293T cells, followed by HALOS-mediated HCR assembly on cell surfaces, produced Cy3 and Cy5 labeling, confirming successful RNA target detection and signal enhancement (Figs. 6(e) and S17). Comprehensive specificity controls showed no signal in cells lacking HALOS 31, target RNA, or containing scrambled HALOS sequences, establishing high selectivity with minimal background (Figs. 6(e) and S17). Confocal microscopy revealed HALOS-mediated target detection (Cy3) on the membrane, and the elevated Cy5 signal confirmed robust signal amplification by HCR compared to H1 alone (Figs. 6(e) and S17). Quantitative analysis revealed 6.8× amplification by confocal microscopy and 8.2× enhancement by flow cytometry, computed as background-corrected net signal (1,571 vs. 192 a.u.; raw 1,739 vs. 360 a.u., Fig. 6(g)) (Figs. 6(f–g) and S17). This strategy overcomes the sensitivity limitations of single-reporter detection by coupling target recognition to HCR amplification, enabling real-time monitoring of low-abundance intracellular RNA targets while preserving cellular viability. This establishes membrane-confined HCR as a proof-of-concept for lysis-free RNA detection in live cells, with potential applications in disease diagnosis, therapeutic monitoring, and fundamental research.

### Dual-reporter labeling validated transmembrane DNA branch migration across the membrane

To establish whether the recognized target genuinely traverses the bilayer, we sought direct, quantitative evidence for transmembrane DNA branch migration. HALOS 31 insertion and target recognition may induce toroidal nanopore formation, creating a hydrophilic microenvironment within the hydrophobic lipid bilayer, consistent with prior reports of DNA-induced toroidal lipid pores. ^36,48,57^ This microenvironment would enable T-Cy3 hybridization with HALOS across the bilayer and promote spontaneous hairpin opening outside the membrane, forming thermodynamically stable complexes with the target DNA. To test for transmembrane crossing, we designed an extended target (T′-Cy3) with an extended branch migration domain (d) for an orthogonal reporter labeling (R-AF488). Successful transmembrane migration yields both Cy5 and AF488-labeling on GUVs by recognizing both the open hairpin (b*) and extended target (d) domains (Figs. 7(a–b)). NUPACK analysis and gel electrophoresis established reporter orthogonality with no cross-reactivity, while solution-phase dual-labeling validated the experimental design (Fig. S18).

Individual reporter labeling of T′-Cy3-detected HALOS on GUVs established specificity, with confocal imaging showing no inter-channel fluorescence bleed-through (Figs. 7(c), left and middle, and S19). Subsequent incubation with both reporters (SA-TrCy5 and SA-TrAF488) produced simultaneous dual labeling, indicating transmembrane target hybridization with the HALOS stem during target detection and hairpin opening (Figs. 7(c) right and S19). Experiments with live HEK293T cells validated cellular compatibility, with co-localization of both reporters on the cell membrane following external HALOS 31 anchoring and T′-Cy3 detection (Fig. S19).

Quantitative analysis of dual reporter-labeled GUVs from the T′-Cy3 detected pool revealed 53±5.0% labeling with SA-TrCy5 and 39±4.9% with SA-TrAF488 (Fig. 7(d)). The higher rate of SA-TrCy5 labeling revealed that the hairpin opening process surpassed the extended target DNA branch migration events across the membrane. The lower AF488 labeling likely reflects incomplete T′-Cy3 migration events within the experimental time frame (t=120 min). These results provide mechanistic insights and quantitative evidence for transmembrane DNA branch migration using the HALOS system.

### Proximal split-initiator induced HCR (ψ-HCR) validates transmembrane branch migration with 5.9× signal amplification

Proximity-dependent ψ-HCR confirmed transmembrane DNA branch migration through proximity-induced initiator formation and activation of HCR. This approach combines sequences from the extended target (T′) and open hairpin to create split-initiator domains that enable ψ-HCR initiation only when both split-initiator sequences are present in close proximity, thereby minimizing the non-specific background signal. Absence of either initiator prevents amplification, as validated by gel electrophoresis, which showed signal amplification exclusively when both split initiators were present in the solution (Fig. S20). We then applied this principle to T′-Cy3 branch migration on HALOS 31, performing membrane-localized ψ-HCR (Fig. 7(e–f)).

GUV-based validation established proximity-dependent amplification following transmembrane branch migration. Confocal microscopy revealed enhanced Cy5 signal intensity compared to H1 labeling alone, establishing efficient ψ-HCR amplification on the membrane (Fig. S21). Control images with single initiator domains (T-Cy3) showed no HCR propagation, demonstrating minimal false-positive amplification and validating the proximity requirements of the initiator domains. We next validated ψ-HCR amplification in live A549 and HEK293T cells. Confocal microscopy showed successful 2′OMe-T′-Cy3 detection with HALOS 31 anchored on cell membranes (Cy3) and distinguished signal amplification (Cy5) compared to the H1 reporter alone (Figs. 7(g) and S22). Control experiments lacking the split-initiator domain produced only Cy3 labeling, validating specificity. Quantitative analysis revealed background-corrected amplification of ∼5.9× by confocal microscopy in A549 cells (Fig. 7(h)) and ∼2.6× by flow cytometry in HEK293T cells (Figs. 7(i) and S22), with differences reflecting the cell line and detection method. These results demonstrate that proximal split initiator formation on HALOS-mediated signaling and ψ-HCR enable specific transmembrane signal amplification with quantitative enhancement and minimal background interference.

## Discussion

Here, we present a synthetic transmembrane signal transducer using an amphiphilic DNA hairpin, which shows the ability to detect intra-vesicular and cellular nucleic acid targets without significant perturbation to membrane integrity. By covalently anchoring only two cholesterols into the internal HALOS domain and without any charge neutralization coupled to the DNA phosphate backbone, we introduce a transmembrane domain in HALOS 31 to hold the nanosensor within the hydrophobic bilayer of the lipid membrane. ^43,52,57^ MD simulations indicated incorporation and stability of several variants and conformations of HALOS within the membrane. The ∼180*^◦^* inter-cholesterol angle stabilizes an axial membrane orientation as the preferred conformation for both unbound and target-bound HALOS within the bilayer. Live-cell microscopy showed that even after repetitive washing and dilution cycles, HALOS 31 remained strongly adhered to the membrane and functionally inserted, detecting intracellular targets. Our studies revealed that target detection is followed by a conformational switch in HALOS 31 across the membrane, leading to signal transduction by reporter labeling.

Transmembrane proteins, especially GPCRs, transduce signals ‘outside-in’ across the membrane through extracellular ligand binding, a receptor conformational change, and activation of a heterotrimeric G protein by GDP-GTP exchange, triggering downstream intracellular signaling cascades. ^7,64^ Various GPCRs also participate in integrin ‘inside-out’ signaling, which directs cell adhesion, spreading, and mechanotransduction. ^65–67^ In this study, we present a synthetic analog of GPCRs which transduces signal ‘inside-out’ and vice versa based on the direction of ligand-target addition. To date, ∼35% of all FDA-approved drugs target GPCRs, ^4,10,68,69^ so a synthetic inside-out analog such as HALOS is of considerable interest as a programmable, sequence-specific receptor for molecular sensing and, prospectively, therapeutic monitoring.

The current HALOS-mediated nucleic acid detection system utilizes synthetic target sequences limited in length up to ∼100-nt due to synthesis constraints, and lacking the secondary structures typical of cytosolic short non-coding RNAs such as miRNAs or tRNAs. Detection of longer cytosolic RNAs, such as mRNAs, which possess highly complex secondary structures, was not pursued in this study. At present, each HALOS variant is capable of detecting only a single target sequence, with multiplexed detection yet to be explored in future studies. Temporal limitations of detection and signal amplification, on the scale of minutes to hours, are mainly attributed to the slow DNA branch migration across the membrane and the kinetics of in situ HCR on the live cell membrane. Enhancing multiplexed detection and temporal resolution will require HALOS variants with accelerated branch migration kinetics, combined with faster isothermal amplification strategies, to reduce response times currently limited by the rate of toehold-mediated strand displacement across the membrane. ^70^

In addition, incorporation of 18:1 LPC, a monoacyl phospholipid, into biologically relevant membranes (POPC and EggPC) substantially influences membrane fluidity, enhancing HALOS-mediated signal transduction events. LPC induces positive membrane curvature and disrupts lipid packing, ^71^ with bilayer integrity maintained up to approximately 30% LPC (molar ratio) in POPC and only about 10% in EggPC membranes, beyond which stable vesicle formation is precluded and coherent bilayer structure is lost. This disruption reflects greater disorder in EggPC compared to POPC bilayers. Modulation of membrane biophysical properties by LPC incorporation improves liposome-based therapeutic delivery by enhancing membrane permeability and enabling faster delivery. Moreover, LPC can alter cell membrane permeability to the extent of inducing necrotic cell death, underscoring its physiological and pathological significance in cell signaling and membrane dynamics. ^72^ Importantly, in our study LPC was incorporated only into synthetic GUVs to enhance transduction efficiency, while live-cell detection required no LPC, avoiding such cytotoxicity. Thus, LPC incorporation provides a strategic avenue to tune membrane characteristics for improved biointerfacing and therapeutic outcomes in membrane-based synthetic biology applications.

HALOS-mediated signal transduction in mixed synthetic cell populations demonstrates high molecular specificity, triggering inside-out signaling exclusively in cells containing the defined target nucleic acid sequence, while cells without this sequence remain inactive. This precise target recognition enables discrimination within heterogeneous cell populations, which could facilitate sequence-based cell sorting without affecting traditional cell surface receptor based isolation strategies or their biological functions. Such specific recognition could support cell-based therapies; for instance, engineered T-cells recognizing cancer-specific nucleic acid markers might be selectively isolated for adoptive cell therapies targeting tumors or hematologic malignancies. ^73^ Similarly, stem cell-based immunotherapies benefit from isolating populations modulated by defined molecular signals. ^74^ Thus, HALOS-mediated signaling offers a programmable basis for sequence-specific cell recognition, with potential for sequence-based cell purification that could enhance the purity, efficacy, and safety of cell-based therapeutics in oncology. This approach builds on the paradigm of artificial transmembrane signaling in synthetic cells, where membrane-anchored synthetic receptors mediate transmembrane signal transduction through direct molecular recognition rather than passive diffusion of signaling molecules across the membrane, analogous to natural receptor mechanisms.

Finally, sensitive lysis-free detection of intracellular targets coupled with HCR-based signal amplification offers substantial biological and diagnostic advantages. In situ HCR, which leverages programmable DNA hairpins, enables enzyme-free, isothermal, and highly specific amplification under physiological conditions, preserving cell viability and avoiding genetic manipulation. ^75,76^ Advances in branched HCR designs further enhance efficiency, sensitivity, and specificity in fixed cell mRNA imaging. ^77^ HALOS integrates membrane-anchored DNA nanodevices with HCR, enabling the amplified detection of low-abundance RNA in live cells, avoiding complex intracellular delivery or genetic engineering, and facilitating applications such as real-time imaging of gene expression and extracellular mRNA biomarkers at the single-cell level. Compared to traditional intracellular nucleic acid detection approaches, such as molecular beacons, ^78^ FISH, ^79^ nano-flare, ^80^ and genetically encoded sensors, ^81^ which require probe delivery, cell fixation, or genetic manipulation and often lack signal amplification, HALOS-HCR offers amplified signal transduction on live membranes with minimal perturbation. Looking forward, combining the lysis-free HALOS-HCR signal amplification platform on live cell membranes promises early biomarker detection, therapeutic monitoring, precise cell isolation, and advances in cell-based therapies, bridging synthetic biology to molecular diagnostics and translational medicine.

## Acknowledgments

The authors thank E. Winfree, L. Qian, A. Aksimentiev, D. Karna, P. Chopade, Y. Hassan and M. Sibouth for their insightful discussions and valuable comments on our work. H. Joshi thanks the supercomputer facility for providing access through the NSM PARAM SEVA at IIT Hyderabad, India.

## Methods

### DNA-cholesterol synthesis using DNA synthesizer

Cholesterol-conjugated DNA was either obtained from Integrated DNA Technology or synthesized in the laboratory using a DNA synthesizer (Applied Biosystems 3400 model). Synthesized DNA (Tables S1–S6) was cleaved from the solid CPG support using a 30% ammonia solution, in accordance with the manufacturer’s protocol. Crude DNA was then purified using an Agilent 1260 HPLC and analyzed using an Agilent 6530 Quadrupole Time-of-Flight ESI-MS (Table S7). Pure DNA-cholesterol conjugates were reconstituted in ultrapure water and stored at –20 *^◦^*C as 10–20 µL lyophilized aliquots.

### DNA-fluorophore conjugation

Both 3^1^ and 5^1^ modified DNA-fluorophores were synthesized either using a DNA synthesizer or by amine-NHS ester coupling methods. Cy3 and Cy5 fluorophore-conjugated DNA were synthesized using a DNA synthesizer, while AF488 fluor coupling with DNA was carried out using amine-modified DNA and AF488 NHS ester (Lumiprobe, 11820). In a typical conjugation method, an aqueous solution of amine-modified DNA in 0.1 M sodium bicarbonate buffer was mixed with AF488 NHS ester (5 equiv) in 10% DMSO (v/v) for 12 hours. Conjugates were then purified using HPLC and analyzed using ESI-MS (Tables S4, S5 and S7).

### Molecular dynamics simulations

All-atom MD simulations were performed using NAMD3^82^ with periodic boundary conditions and the particle mesh Ewald (PME) method implemented for long-range electrostatics. ^83^ Simulations were conducted in the isothermal-isobaric ensemble, with pressure and temperature regulated using the Nosé-Hoover Langevin piston ^84,85^ and Langevin thermostat, ^86^ respectively. For the membrane systems, anisotropic pressure coupling was applied to maintain a constant ratio between the X and Y dimensions, allowing the Z dimension (normal to the membrane) to fluctuate independently.

The CHARMM36 force field ^87^ was used to describe all bonded and non-bonded interactions among DNA, lipid bilayer membranes, water, and ions. Cholesterol and TEG linker parameters were adopted from previous studies. ^43,52,88^ A cutoff scheme of 8-10-12 Å was employed for van der Waals and short-range electrostatic interactions. All systems were simulated using a 2 fs time step for integrating the equations of motion. The SETTLE algorithm ^89^ constrained the water geometry, and all other covalent bonds involving hydrogen were constrained using the RATTLE algorithm. ^90^ Coordinates were saved every 19.2 ps. Simulation trajectories were analyzed and post-processed using VMD ^91^ and CPPTRAJ.^92^

Initial PDB structures of the DNA hairpins were created using the Alphafold3 server, ^93^ yielding plausible models with a pTM-score of 0.4. An additional phosphate group was incorporated into the DNA backbone to enable the covalent attachment of the cholesterol-TEG linker using the psfgen plugin in VMD. ^94^ In-house TCL scripts were used to construct models with DNA hairpins anchored in phospholipid membranes, positioning cholesterol-anchored HALOS in both axial and lateral orientations relative to POPC bilayers. All membrane lipids within 0.1 Å of DNA hairpin or cholesterol atoms were removed, and the number of lipids was balanced between the upper and lower leaflets. All the systems were solvated in a TIP3P water box. Potassium and chloride ions were added to a 0.15 M concentration of KCl in the system using the Autoionize plugin of VMD. Thus, the assembled all-atom models of all the simulated systems are summarized in Tables S8 and S9.

Following assembly, each system underwent 4,800 steps of energy minimization using the conjugate gradient method. After minimization, each system was equilibrated in the NPT ensemble (P = 1 bar, T = 300 K) for 50 ns with harmonic restraints (1 kcal mol*^−^*^1^ Å*^−^*^2^) applied to all non-hydrogen atoms of the DNA hairpins, referenced to their initial coordinates. For membrane-embedded and aqueous systems, restraints were maintained for 50 ns and 5 ns, respectively, followed by ∼1,000 ns of unrestrained equilibration. Because the accessible 1 µs timescale is many orders of magnitude shorter than the experimental insertion and signaling kinetics, each system was initialized in a defined conformational state (aqueous, lateral, axial, or target-bound) to assess its structural stability, rather than to capture spontaneous insertion or strand-displacement events.

### Hydrogen bond and cholesterol angle analysis

Hydrogen bond analysis was performed using a custom TCL script. A hydrogen bond was defined by a donor-acceptor distance cutoff of 3 Å and a donor-hydrogen-acceptor angle ≤ 45*^◦^*. The Watson-Crick (WC) hydrogen bonds between paired DNA bases were computed for each frame of the simulation. For visualization, the mean hydrogen bonds per frame were calculated for each frame of the simulation (1 ns each) and plotted as Gaussian distribution plots. For the broken base pair representation, the hydrogen bond counts were normalized by the total number of base pairs in each system and displayed as heat maps. Cholesterol angles were determined by considering all atoms of the cholesterol molecules and generating a vector with the DNA stem within the membrane as the perpendicular axis. The angle between two cholesterol vectors was measured per frame of simulation time (1 ns each) and plotted in a circular plot ranging from 0 to 360*^◦^*.

### Gel electrophoresis

Native polyacrylamide gel electrophoresis (PAGE) was conducted using an 8% (w/v) precast gel in 1× TBE supplemented with 12.5 mM MgCl_2_ for signal transduction and hybridization chain reaction (HCR). Electrophoresis employed a 1× TBE buffer composed of 89 mM Tris, 89 mM boric acid, and 2.0 mM EDTA at pH 8.4, with an additional 12.5 mM MgCl_2_. DNA strands were annealed using a standard protocol. Samples were initially heated to 90 *^◦^*C, eventually cooled to 76 *^◦^*C at a rate of 2 *^◦^*C/5 min, and then gradually cooled down to 24 *^◦^*C at a rate of 4 *^◦^*C/5 min, followed by storage at 4 *^◦^*C. DNA was loaded into a gel at a typical concentration of 10 pmol (1 µM×10 µL) and run at 100 V for 60 min. Gels were then post-stained with 0.5 µg/mL ethidium bromide (EtBr) solution for 10 min and imaged under a UV transilluminator. HCR was conducted using 1% (w/v) agarose gel electrophoresis in 1× Sodium Borate buffer (10 mM NaOH, pH adjusted to 8.5 with boric acid), supplemented with 12.5 mM MgCl_2_. Gels were prestained with SYBR gold (0.5 µg/mL), loaded with 10 pmol (1 µM×10 µL) of sample, and run at 100 V for 90 min under ice-cold conditions, then imaged using a UV transilluminator.

### GUV preparation

Cy3 conjugated targets (T-Cy3)-encapsulated GUVs were prepared using a standard electroformation protocol. ^56^ Briefly, in this study, 10 mM lipid stocks (∼5 µL) in chloroform (Avanti polar lipids: POPC, 850457; EggPC, 840041; 18:1 LPC, 845875; purity *>*99%) were homogeneously spread over the conductive side of the indium tin oxide (ITO) coated on two glass slides, and the solvent was evaporated under vacuum for 20 min. All lipid stocks were stored in chloroform at −20*^◦^*C for long-term (1–2 years) storage and allowed to warm to room temperature before every use. For the preparation of GUVs with mixed lipids, the lipid stocks were premixed in a molar ratio in a glass vial prior to deposition onto ITO glass slides. A rubber ring was then placed on the lipid-coated ITO slide and filled with 250 µL of 300 mM sucrose solution (∼300 mOsm/kg) containing 250 nM T-Cy3. Another ITO-coated glass slide was placed on top, and the chamber was connected to the electrode of the Vesicle Prep Pro (Nanion). An AC field (3 V, 10 Hz) was applied via the electrodes for 128 min while the solution was heated to 37 *^◦^*C. GUVs were collected immediately after electroformation, stored at 4 *^◦^*C, and used within one week.

### Confocal microscopy

Confocal images were captured using a Nikon AX R Ti2 laser scanning confocal microscope with a plan apochromat 60× oil OFN25 DIC N2 objective (NA 1.42). Four monochromatic lasers (405, 488, 561, and 640 nm) were used to excite the respective fluorophores. Emissions were filtered using highly sensitive GaAsP adjustable emission filters (503–530 nm for the 488 nm laser line, 571–625 nm for the 561 nm laser line, and 662–737 nm for the 640 nm laser line). The laser intensity and detector gain were maintained constant for each channel during the confocal scanning. Multichannel images were recorded using sequential scanning at 1024×1024 pixel resolution with 2× averaging and a 1 AU pinhole size. 3D confocal images were recorded at 512×512 pixels in the XY-direction and 500 nm in the Z-direction resolution. A total of 31 slices were recorded over a 15 µm depth of the GUVs sample, and 3D images were reconstructed using the ImageJ 3D viewer plugin.

### Membrane fluidity measurement

Membrane fluidity of each lipid composition was measured using a membrane-sensitive probe, Laurdan (MedChem Express, HY-D0080), and GP values were calculated using the following generalized polarization equation: GP= [I_B_–I_R_]/[I_B_+I_R_]. Laurdan (5 µM) was incubated with freshly prepared GUVs in a 300 mM sucrose solution for 15 min and subjected to a plate reader for fluorescence measurement across the wavelength range (430–600 nm with 1 nm steps) after excitation at 390 nm (Laurdan). GP values were calculated using I_B_ at 436 nm and I_R_ at 512 nm. For fluorescence spectral scanning imaging using confocal microscopy, the GUVs incubated with Laurdan were excited with a 405 nm laser, and spectral scanning was carried out using a narrow 5 nm emission window (420–600 nm). GP images were generated from I_B_ and I_R_ confocal spectral scanning images using a custom-built GP plugin in ImageJ.

### Sample preparation for signal transduction across the GUV membrane

Electroformed GUVs (∼100 µL) were centrifuged at 300 g for 5 min after dilution with 1 mL homo-osmotic (∼300 mOsm/kg) 1× imaging buffer (1× TAE + 150 mM NaCl+12.5 mM MgCl_2_) and the outside buffer was exchanged to remove the excess free T-Cy3. Post centrifugation, the supernatant was discarded, and the GUVs pellets were further suspended in 100 µL of fresh 300 mM sucrose solution and stored at 4 *^◦^*C for at least 7 days. Excess complementary DNA (5 µM, 20× excess) against T-Cy3 (250 nM) was added to the GUVs solution for 1 hour to completely quench the T-Cy3 traces outside the GUVs. For experiments with mixed GUVs, the GUVs were mixed thoroughly before adding complementary components. Then, annealed HALOS 31 (250 nM) was incubated with ∼1–2 µL of GUV in 100 µL of 1× imaging buffer within a coverslip bottom 96 well plate (Ibidi, 89621) for sufficient time (2–3 h) for the sensor insertion and DNA migration across the GUV membrane. R-Cy5 (100 nM) was then incubated with the GUVs for 60 min and subsequently diluted to ∼500 µL using 1× imaging buffer to reduce the background signal of R-Cy5 prior to proceeding with confocal microscopy.

### RNA transfection protocol for live-cell imaging and flow cytometry

HEK293T and A549 cells were plated at 70–90% confluency either in 8-well glass-bottom chamber slides (Ibidi, 80806) for confocal microscopy or in 6-well dishes (Greiner bio-one, 657160) for flow cytometry. For healthy cell populations, prior to transfection, the culture media was replaced with Opti-MEM (200 µL for confocal; 2 mL for flow cytometry) and incubated for 1 hour. For confocal imaging, 2′-OMe RNA target (100 nM) was mixed with 1.5 µL P3000 reagent and diluted in 50 µL Opti-MEM, while Lipofectamine (1 µL) was diluted separately in 50 µL Opti-MEM; both solutions were vortexed, centrifuged, combined by pipetting, and incubated for 15 min before dropwise addition to cells. For flow cytometry, 100 nM target was mixed with 7.5 µL P3000 reagent diluted in 75 µL Opti-MEM, and 5 µL Lipofectamine was diluted in 75 µL Opti-MEM; similarly mixed and incubated before addition. Transfections were conducted under standard culture conditions. After 60–90 min, the cells were washed three times with DPBS (1 min incubation per wash) to eliminate extracellular non-transfected RNA and then subjected to imaging or flow cytometry.

### HCR protocol

HCR was performed in solution using 1× imaging buffer for characterization using gel electrophoresis. 3 µL each of 1 µM H1, H2, and initiator (HALOS 31+T-Cy3) were mixed in a PCR tube for the HCR. In the absence of any reactant, 3 µL of 1× imaging buffer was added to adjust the total volume of 9 µL in the sample mixture. For completeness, the reaction was allowed to proceed at room temperature for 24 hours before characterization using gel electrophoresis.

To perform HCR on the GUVs surface, HALOS 31 (2.5 µM×10 µL) was incubated with T-Cy3 encapsulated GUVs solution in 100 µL of 1× imaging buffer for 3 hours to perform DNA branch migration across the membrane, followed by the addition of H1 and H2 (1 µM×10 µL each) to the GUVs solution. After 1 hour, confocal microscopy images were acquired after dilution to ∼500 µL using 1× imaging buffer.

For HCR on live cell membranes, 2′-OMe-RNA-Cy3 (2′-OMe-T-Cy3, 41-nt) transfected cells in 8-well coverslip bottom chamber slides (Ibidi, 80806) were subjected to removal of the extracellular targets by repetitive washing using DPBS (3 times), and treated with complementary DNA (4 µM, 40× excess) in opti-MEM media supplemented with Aurintricarboxylic acid (ATA, a nuclease inhibitor; 50 µM) and MgCl_2_ (12.5 mM) for 20 min in live cell culture conditions for complete quenching of extracellular non-transfected RNA targets. Cells were then washed with DPBS (3 times) and treated with HALOS 31 (250 nM) in Opti-MEM media supplemented with ATA (50 µM), MgCl_2_ (12.5 mM), and complementary DNA (2 µM) for 20 min at 37 *^◦^*C. H1 and H2 (250 nM each) diluted in opti-MEM (200 µL), supplemented with 50 µM ATA and 12.5 mM MgCl_2_ were then incubated with the cells for 30–60 min at 37 *^◦^*C. Similarly, ψ-HCR was performed by transfecting 2′-OMe-T′-Cy3 (60-nt) and choosing the respective H1 and H2 reporters. 2′-OMe-T-Cy3 (41-nt), lacking the extended split-initiator domain, was considered as a negative control for ψ-HCR. Cells were finally washed with DPBS, the Opti-MEM media exchanged, and confocal microscopy performed.

### Flow cytometry

2′-OMe RNA transfected cells from a 6 well dish were washed, trypsinized, centrifuged at 800 rpm for 4 min, and resuspended in opti-MEM media in a microcentrifuge tube. Cells were treated with complementary DNA (4 µM, 40× excess) for 20 min at 4 *^◦^*C. Cells were then washed out by centrifugation and 200 µL HALOS 31 (250 nM) was incubated for 20 min at 4 *^◦^*C. After excess unbound sensor removal by centrifugation, the reporters (H1 and H2, 250 nM each) diluted in opti-MEM (200 µL) were then incubated with the cells for 30–60 min at 4 *^◦^*C. Cells were finally centrifuged, excess unreacted reporters removed, and pellets resuspended in Opti-MEM before proceeding with flow cytometry. Similarly, ψ-HCR was performed by transfecting OMe-T′-Cy3 (60-nt) and selecting the respective H1 and H2 reporters. OMe-T-Cy3 (41-nt) was used as a negative control for ψ-HCR. All samples were diluted to 500 µL in Opti-MEM and subjected to flow cytometry analysis. 10,000 scattering events were collected upon excitation of Cy3 (561 nm) and Cy5 (642 nm) dyes, and the live cells were adequately gated from the scattered plot of single cells.

### Image analysis

Confocal images were post-processed in ImageJ with identical intensity windows across channels. GP images were generated using a custom-coded ImageJ macro plugin. High-throughput analysis of GUVs was performed by isolating the coordinates of each GUVs using DisGUVery ^95^ and performing an algorithm-based membrane intensity calculation using custom-coded Mathematica and Matlab programming. Intensity plots and standard deviations were generated using GraphPad Prism and Mathematica. High-throughput image analysis of live cells was performed by analyzing the kymograph of the cell membrane using a custom-coded ImageJ macro plugin and Mathematica programming.

### Quantification and data analysis

All graphs were created using GraphPad Prism 9 and Mathematica. Error bars represent the SD, unless otherwise noted. Membrane-associated fluorescence intensities for HCR and H1-only samples were quantified after background subtraction. For confocal data, background was estimated from cell-free regions of the field of view. For flow cytometry, background was determined using target-encapsulated cells treated with reporter dyes alone. The fluorescence-fold enhancement was calculated as the ratio of mean intensities between the two groups. The Statistical comparisons between two independent groups were performed using the two-sided Mann–Whitney *U* test, a non-parametric test chosen because the per-vesicle and per-cell fluorescence-intensity distributions were right-skewed and could not be assumed to be normally distributed. Throughout, N_GUV_ and N_cells_ denote the numbers of individual vesicles and cells analyzed, respectively, and N the number of independent experiments. For panels comparing more than two conditions, a single pre-specified comparison (HCR amplification versus the H1-only reporter) was evaluated per panel; no correction for multiple comparisons was therefore applied. Exact P values are reported in the figure legends; NS, not significant.

Bootstrap resampling was used to estimate uncertainty in the mean Cy3 and Cy5 ring counts for GUVs. Bootstrap calculations were performed using a custom MATLAB code. First, the Cy3 and Cy5 rings were assigned binary numbers 0 and 1 for negative and positive membrane labeling, respectively. For bootstrapping, N=1,000 simulated datasets were generated by random resampling with replacement from the original dataset, preserving the size in each iteration. Each resampled dataset was stored as an independent bootstrap realization, from which the mean value was computed iteratively to obtain the bootstrap distribution of the means. Based on this distribution, the final mean value and standard deviation were determined across all bootstrap realizations of the Cy3 and Cy5 ring contents, thereby quantifying the uncertainty of the estimated mean without assuming an underlying parametric distribution.

## Data Availability

The source data in this study have been deposited in the repository. https://doi.org/10.5281/zenodo.21014442

## Code Availability

Scripts and source code used for data processing are available at https://github.com/Ranjans04/HALOS-Signal-Transducer.

## Supplementary Information

### Supplementary Materials and Methods

Unmodified DNA was purchased from Integrated DNA Technologies (IDT). Modified DNA was synthesized in the laboratory using a solid-phase DNA synthesizer (Applied Biosystems 3400). Reagents and phosphoramidites were purchased from Glen Research, Sigma-Aldrich, and Lumiprobe. Synthesized DNA was purified by high-performance liquid chromatography (HPLC, Agilent 1260) using a c18 reverse-phase column. Solvents used as the mobile phase in HPLC were solvent A: 100 mM Triethylammonium Acetate buffer, pH 7.0 (TEAA; Sigma Aldrich, 625718) and solvent B: Acetonitrile (Sigma Aldrich, 439134). DNA conjugates were analyzed by mass spectrometry using electrospray ionization (ESI-MS) on an Agilent 6530 Quadrupole Time-of-Flight LC/MS System. The intact mass analysis of DNA was carried out by performing deconvolution using BioConfirm software (Agilent). Unmodified DNA purchased from IDT was filtered through a molecular weight cut-off (MWCO) filter to remove all traces of salts and reagents using ultrapure water. DNA stocks were adjusted to 100 µM in ultrapure water prior to the experiments. The molar extinction coefficient (*ε*) of all DNAs was obtained from the IDT DNA analysis website. Native PAGE analysis was performed using a Bio-Rad Mini-PROTEAN Tetra Vertical Electrophoresis Cell (Bio-Rad, 1658004), and agarose gel electrophoresis was performed using a Thermo Fisher Scientific Owl Easy Cast (B2-BP) setup. GUV formation was carried out using a Nan]i[on vesicle prep pro setup (Nanion Technologies GmbH). Confocal images were acquired using a Nikon AX R confocal setup equipped with a GaAsP detector, and image analysis was carried out using Fiji-ImageJ, DisGUVery, custom-coded Mathematica, and MATLAB.

#### In-house DNA synthesis, conjugation, and characterization

HALOS 14 and HALOS 31 were purchased from IDT, while HALOS 21–28 were synthesized in-house using an automated DNA synthesizer via standard solid-phase phosphoramidite chemistry in the 3^1^ to 5^1^ direction. Each synthesis cycle included sequential detritylation, coupling of phosphoramidite monomers, capping, and oxidation steps to ensure a high coupling efficiency. Following chain assembly, the oligonucleotides were cleaved from the solid support, deprotected, and purified using reverse-phase HPLC. Fluorophores (Cy3, Cy5, and ATTO488) and biotin modifications were introduced either during synthesis via pre-labeled phosphoramidites or post-synthetically using NHS-ester-amine conjugation chemistry. Purified conjugates were further analyzed by ESI-MS equipped with BioConfirm to confirm the intact molecular masses. For ESI-MS, ∼10 µM DNA samples (10 µL) were injected in ultrapure water, using 0.1% NH_4_OH in ultrapure water as the mobile phase.

#### Modeling membranes with varied lipid compositions

All simulated membrane systems contained 64 lipid molecules per leaflet. Initial bilayer configurations were built using the CHARMM-GUI membrane builder ^91^ with the following compositions: POPC (64 POPC per leaflet), EggPC (40 POPC and 24 SLPC per leaflet), and POPC:cholesterol (45 POPC and 19 cholesterol per leaflet). Parameters for LysoPC (18:1), which are not available in CHARMM36, were assembled by extracting the molecular structure from the Human Metabolome Database (HMDB ID: HMDB0010385) and generating parameters using CGENFF. ^96^ Reference LysoPC species (14:0 and 16:0) in CGENFF provided parameter and charge penalties of 1 and 0.491, respectively, supporting accurate simulation of LysoPC (18:1) in the present study.

#### Structure building and optimization of Cy3 dye

The initial structure of Cy3 dimethyl was obtained from PubChem (CID: 5705412) and subsequently modified by the addition of ethanol and ethane groups using Gaussian 16. Modified structures underwent geometry optimization and vibrational frequency analysis at the density functional theory (DFT) level, employing the B3LYP functional with the 6-31+G basis set to balance computational efficiency and polarization accuracy. Vibrational frequency calculations confirmed the optimized geometry as a true minimum by the absence of imaginary frequencies. The Gaussian input included iop(6/33=2,6/42=6) keywords to enable a detailed natural bond orbital analysis and ensure a proper charge distribution for subsequent simulations. Force field parameters for Cy3 were derived using the CHARMM General Force Field (CGenFF). ^96^

#### RMSF and contour length calculation

All RMSF analyses were performed over the entire simulation trajectory of 1 µs using CPPTRAJ. Calculations were performed by selecting backbone atoms of the DNA bases to assess the structural fluctuations.

Contour length distributions were obtained by summing the rise base pair parameters of all paired DNA bases. Rise values were extracted using the ‘nastruct’ module of CPPTRAJ, and the resulting distributions were used to evaluate DNA overstretching behavior around the hydrophobic modifications.

#### Sample preparation of HALOS stability studies in GUVs

1. ATTO488 dyes encapsulated (without target DNA) GUVs were prepared by electroformation and resuspended in imaging buffer (1× TAE+150 mM NaCl+12.5 mM MgCl_2_).
2. HALOS variants (250 nM) were thoroughly mixed with GUVs and incubated for 30 min at room temperature.
3. Reporter R-Cy5 (13-nt ssDNA, 100 nM) was added and the mixture incubated for 30 min at room temperature.
4. The GUV solution was diluted to ∼5× volume using imaging buffer and subjected to confocal microscopy.

#### Cell culture

HEK293T and A549 cells were cultured at 37 *^◦^*C in a humidified atmosphere containing 5% CO_2_ using Dulbecco’s Modified Eagle’s Medium (DMEM, high glucose) supplemented with 10% fetal bovine serum (FBS), 1% antibiotics (100 U/mL penicillin, 100 µg/mL streptomycin; Corning, 30-002-CI) and 1% MEM non-essential amino acids (Corning, 25-025-CI). At ∼80% confluence, the cells were washed using DPBS (pH 7.3), trypsinized, and suspended in a culture medium. For imaging experiments, cells were counted and seeded at a density of ∼10,000 cells per well in 200 µL of culture media into 8-well glass bottom plates (ibidi, 80806). Cells were incubated under the same conditions for 24 hours to reach 80% confluence before imaging.

#### Sample preparation of HALOS stability studies in live cells

1. HEK293T cells seeded on glass bottom chambers were washed with DPBS, incubated with HALOS variants (250 nM) in Opti-MEM media containing ATA (50 µM) and MgCl_2_ (12.5 mM) for 15 min under live cell culture conditions.
2. Unbound probes were removed using DPBS washes (3 times) with 1 min incubation each time.
3. Reporter R-Cy5 (13-nt ssDNA, 100 nM) was incubated with the cells for 15 min in Opti-MEM containing ATA (50 µM) and MgCl_2_ (12.5 mM) under live-cell culture conditions.
4. Unbound probes were removed using DPBS washes (3 times) with 1 min incubation each time, and cells were maintained in Opti-MEM prior to confocal microscopic imaging.

#### Stepwise sample preparation for signal transduction experiment in GUVs

1. 100 µL T-Cy3 (or ATTO488 dye as control) encapsulated (250 nM) GUVs prepared using electroformation were diluted with 900 µL homo-osmotic (∼300 mOsm/kg) 1× imaging buffer (1× TAE+150 mM NaCl+12.5 mM MgCl_2_) in a microcentrifuge tube and centrifuged at 300g for 5 min at room temperature.
2. GUVs settled down as a pellet-like residue, and the supernatant was exchanged carefully with 1 ml of fresh 1× imaging buffer.
3. After two repetitive washings, GUVs were resuspended in 100 µL homo-osmotic sucrose solution. GUV solutions can be stored at 4 *^◦^*C for up to 7 days.
4. The complementary DNA (5 µM, 20× excess) in homo-osmotic 1× imaging buffer (1× TAE+150 mM NaCl+12.5 mM MgCl_2_) was added to GUVs and incubated for 60 min at room temperature for complete quenching of extravesicular targets.
5. Annealed HALOS 31 (250 nM) was incubated with GUVs (1–2 µL) in 100 µL 1× imaging buffer within the coverslip-bottomed 96-well plates for 2–3 h at room temperature.
6. R-Cy5 (100 nM) was added and incubated with the GUVs for 60 min.
7. The GUVs suspension was diluted to ∼500 µL with 1× imaging buffer to reduce the background signal of R-Cy5 prior to proceeding to confocal microscopy.

#### Calculation of the radius of gyration of 13-nt ssDNA R-Cy5 in water

The radius of gyration (R*_g_*) of 13-nt ssDNA R-Cy5 was estimated using a worm-like chain (WLC) model approximation. For a semi flexible polymer of contour length *L* and persistence length *£_p_*, the mean-square radius of gyration is given by

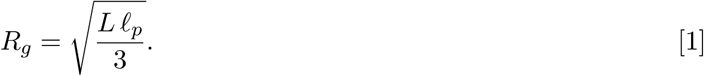

The contour length *L* was calculated from the number of nucleotides (*N*) and the rise per nucleotide (*a* = 0.59 nm) for ssDNA in an aqueous buffer:

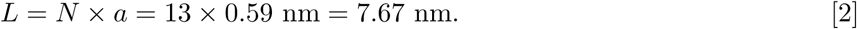

The persistence length of ssDNA under physiological ionic strength was assumed to be *l_p_* = 1.1 nm, consistent with previous single-molecule studies. ^97^ Substituting these values into Eq. 1 gives

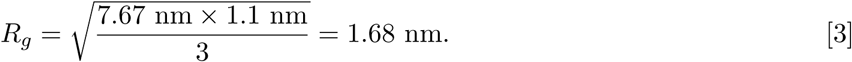

Thus, the radius of gyration of the 13-nt ssDNA-Cy5 conjugate in water was estimated as

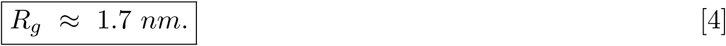

This value represents the equilibrium spatial extension of the ssDNA-Cy5 conjugate in an aqueous solution and is consistent with previous experimental and computational reports for short ssDNA oligomers.

#### Hairpin amplifier stands (H1, H2) design for ψ-HCR

Metastable DNA hairpin monomers (18 bp stem/6-nt loop) for ψ-HCR were designed using NUPACK simulations to optimize their thermodynamic and kinetic properties. Initially, the H1 strand was designed to utilize a toehold domain (9-nt) that initiates binding of the first split-initiator (SI-1) from extended target domain (d), followed by subsequent binding of a SI-2 from open hairpin through recognition of a proximal split-toehold, which would open the H1 hairpin stem and trigger HCR propagation from the opened state. However, gel electrophoresis analysis revealed that, although SI-1 and SI-2 bound the H1-hairpin, they failed to induce efficient stem opening, leading to minimal or no in situ hybridization. This limited propagation was attributed to insufficient thermodynamic stabilization upon binding of the second split initiator.

To enhance the HCR propagation, we designed a second class of H1-hairpin by distributing the initiator binding sites differently: the SI-1 from extended target domain (d) binds both the toehold (9-nt) and a partial stem region (8-bp) of H1, while the SI-2 from open hairpin (b*) binds the remaining stem segment (10-bp). This split-stem binding design increases overall binding affinity and significantly improves the thermodynamic stability of the opened H1 conformation. Consequently, this configuration facilitated more robust and faster ψ-HCR polymerization, as confirmed by pronounced polymerization bands in gel electrophoresis (Fig. S20).

#### Sample preparation for live-cell confocal microscopy

1. Live HEK293T and A549 cells were plated at 70–90% confluency into an 8-well glass-bottom dish using DMEM media.
2. The media was exchanged to opti-MEM prior to 1 h of RNA transfection process. Cells were allowed to remain healthy prior to transfection.
3. For each well, Cy3-labeled 2′-OMe modified RNA targets (100 nM) were premixed with 1 µL of P3000 reagent in a microcentrifuge tube and diluted with 50 µL opti-MEM media and mixed well. 1 µL Lipofectamine reagent was diluted with 50 µL opti-MEM media and mixed well. Both of these tubes were vortexed and centrifuged and combined into a single tube by vigorous mixing and allowed to stand for 15 min at room temperature.
4. The cocktail mixture was added dropwise to the live cells and incubated for 60–90 min to allow transfection.
5. The cells were then washed thoroughly (3 times) using DPBS to remove the excess non-transfected targets.
6. Complementary DNA (4 µM, 40× excess) against RNA targets supplemented with ATA (50 µM) and MgCl_2_ (12.5 mM) was incubated with the cells for 20 min in live-cell culture conditions for complete quenching of extracellular non-transfected RNA.
7. The cells were then washed with DPBS (3 times) and treated with HALOS (250 nM) supplemented with ATA (50 µM), MgCl_2_ (12.5 mM), and 2 µM complementary DNA for 20 min in live-cell culture conditions.
8. Unbound probes were removed by washing the cells using DPBS (3 times) and the reporter strands (Cy5 labeled H1 and H2, 250 nM each) diluted using opti-MEM (200 µL), supplemented with ATA (50 µM) and MgCl_2_ (12.5 mM) were incubated with the cells for 60 min in live-cell culture conditions.
9. Cells were finally washed with DPBS, the media was exchanged with opti-MEM, and the cells were processed for live-cell confocal microscopy.
10. The control studies were performed using cells lacking RNA-target transfection and sensors, with only the H1 reporter.
11. ø-HCR was performed by transfection of Cy3-labeled 2′-OMe modified extended RNA targets (T′-RNA, 60-nt) and choosing respective H1 and H2 reporters. A negative control for ψ-HCR was also performed by considering RNA target (T-RNA, 41-nt) lacking the extended split-initiator domain for HCR.

#### Sample preparation for flow cytometry

1. Live HEK293T and A549 cells were plated at 70–90% confluency into 6-well culture dish using DMEM media.
2. The media was exchanged to opti-MEM (2 mL) prior to 1 h of RNA transfection. Cells were allowed to remain healthy prior to transfection.
3. Cy3-labeled 2′-OMe modified RNA targets (100 nM) were mixed with 7.5 µL of P3000 reagent in a microcentrifuge tube and diluted with 75 µL opti-MEM media and mixed well. 5 µL Lipofectamine reagent was diluted with 75 µL opti-MEM media and mixed well. Both of these tubes were vortexed and centrifuged and combined into a single tube by vigorous mixing and allowed to stand for 15 min at room temperature.
4. The cocktail mixture was added dropwise to the live cells and incubated for 60–90 min to allow transfection.
5. The cells were then washed thoroughly (3 times) using DPBS to remove the excess non-transfected targets.
6. The cells were then trypsinized, centrifuged at 800 rpm for 4 min, and resuspended in opti-MEM (1 mL). This washing step was repeated twice; cells were resuspended in Opti-MEM (200 µL).
7. Complementary DNA (4 µM, 40× excess) against RNA targets supplemented with ATA (50 µM) and MgCl_2_ (12.5 mM) was incubated with the cells for 20 min at 4 *^◦^*C for the complete quenching of extracellular RNA targets.
8. The cells were then centrifuged, resuspended in opti-MEM (200 µL) and treated with HALOS (250 nM) supplemented with ATA (50 µM), MgCl_2_ (12.5 mM), and 2 µM complementary DNA for 20 min in live-cell culture conditions.
9. Unbound probes were removed by washing the cells using DPBS (3 times) and the reporter strands (Cy5 labeled H1 and H2, 250 nM each) diluted using opti-MEM (200 µL), supplemented with ATA (50 µM) and MgCl_2_ (12.5 mM) were incubated with the cells for 60 min in live-cell culture conditions.
10. The cells were then washed with DPBS (3 times) and resuspended in opti-MEM (500 µL). Cells were finally centrifuged, excess H1, H2, and media removed, and pellets resuspended in Opti-MEM (500 µL) before flow cytometry.
11. The control studies were performed using cells lacking RNA targets and HALOS, with only the H1 reporter, following the same HCR protocol described above.
12. (ψ-HCR) was performed by transfecting extended RNA targets (60-nt) and choosing respective H1 and H2. A negative control for ψ-HCR was also performed by considering the RNA target (T-RNA, 41-nt) lacking the extended split-initiator domain for HCR.

#### Flow cytometry protocol

The individually prepared samples were finally diluted to 500 µL opti-MEM media and subjected to flow cytometry studies. 10,000 events were collected from each sample and scanned using excitation of Cy3 (561 nm) and Cy5 (642 nm). Fluorescence scattering from single cells was collected using the respective band-pass filters. The live-cells were adequately gated from the single cell analysis and scattered plots of Cy3 vs. Cy5 intensity were generated.

#### Nuclease inhibition by ATA

Aurintricarboxylic acid (ATA, 50 µM) was employed in live-cell HCR experiments to inhibit extracellular nuclease activity and maintain the stability of the intracellular HALOS-inserted domain and metastable hairpin monomers (H1, H2). ATA is a broad-spectrum inhibitor of nucleases, including DNase I, exonucleases, and restriction endonucleases, acting by binding to positively charged regions of these enzymes and preventing access to DNA substrates, thereby blocking catalytic cleavage. ^98^ In vitro, ATA effectively inhibits DNase I activity at low micromolar concentrations (∼10 µM). During live-cell imaging, lower concentrations (∼10 µM) were insufficient to prevent degradation of single-stranded DNA reporters, necessitating an optimized concentration of 50 µM ATA. This concentration provides robust inhibition of both intra- and extracellular nucleases while minimizing cytotoxicity (which increases significantly above 100 µM), thereby preserving cellular function over the assay duration. The reversible electrostatic and hydrogen bonding interactions of ATA with nucleases confer protection of nucleic acid probes, enabling reliable signal transduction without adverse effects on live-cell viability.

### Supplementary Tables

**Table S1.**
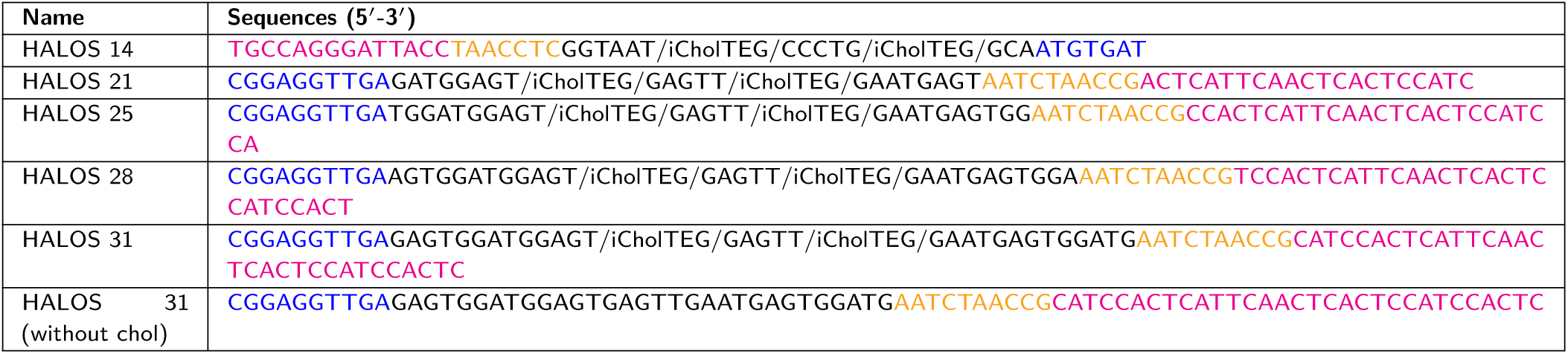
Cholesterol labeled DNA hairpins. Cholesterol labeled DNA hairpin sequences used in the study. Toehold domain (a) of HALOS is highlighted in blue, cholesterol anchored stem (b) in black, loop (c) in yellow-orange and reporter recognition stem (b*) in magenta. Internally modified Cholesterol is abbreviated as /iCholTEG/.

**Table S2.**
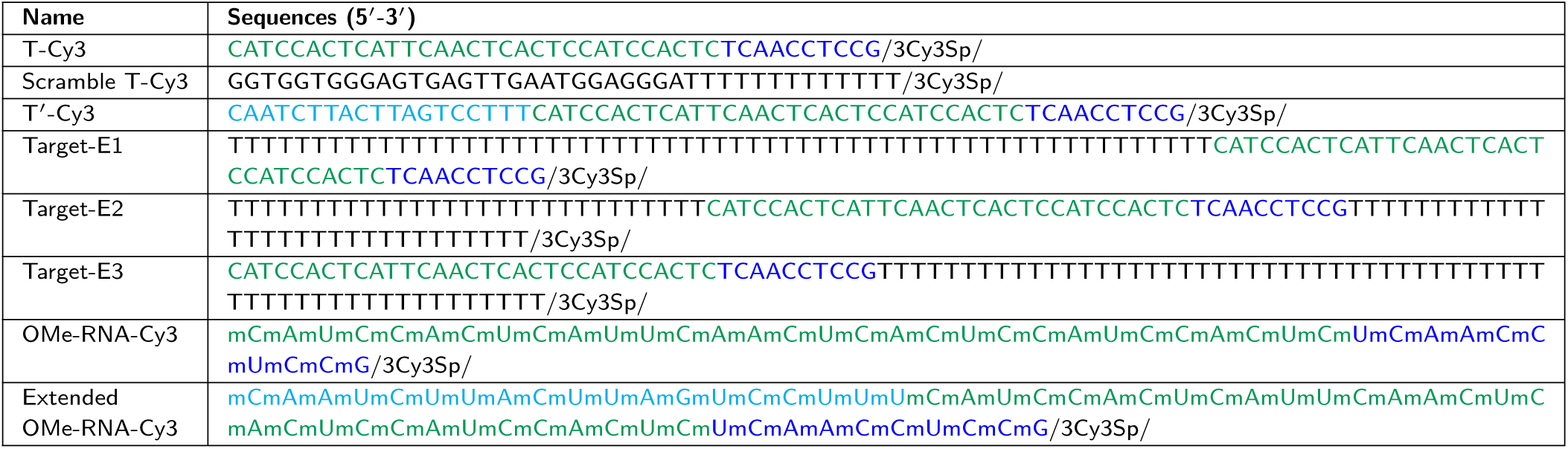
Cy3 labeled target DNA and RNA strands. Cy3 labeled DNA and OMe-modified RNA sequences used as target strands in the study. HALOS toehold binding domain (a*) in target strand is highlighted in blue, stem binding domain (b*) in forestgreen and extended target domain (d) for reporter labeling in cyan. Cy3 modification at 3′ end is abbreviated as /3Cy3Sp/.

**Table S3.**
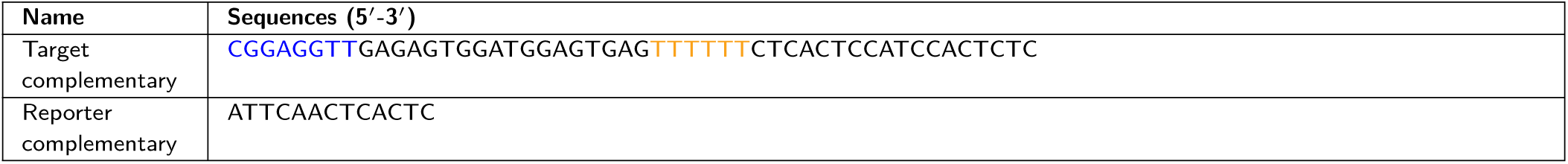
Complementary strands against target nucleic acids. Complementary strand against T-Cy3 and R-Cy5 strands used in the study. For hairpin complementary strand, toehold is highlighted in blue and loop in yellow-orange.

**Table S4.**
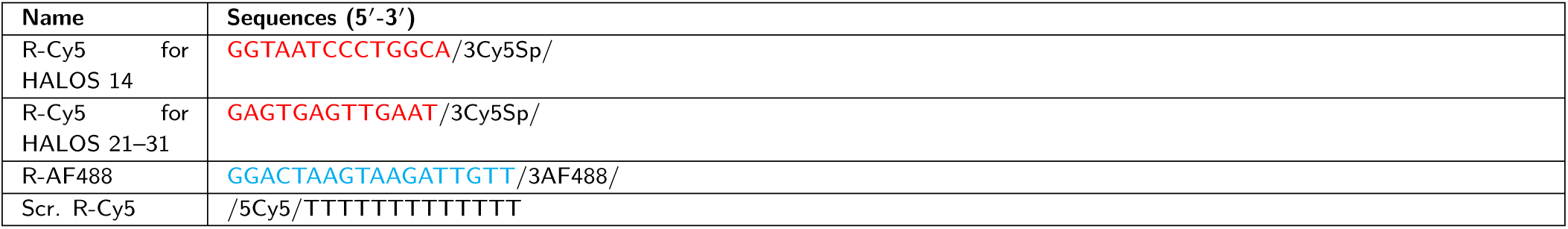
Cy5 labeled ssDNA reporters. Cy5-labeled ssDNA reporter sequences used in the study. Cy5 labeled reporters are highlighted in red and AF488 labeled reporter in cyan. Cy5 modification at 3′ and 5′ end are abbreviated as /3Cy5Sp/ and /5Cy5/ respectively.

**Table S5.**
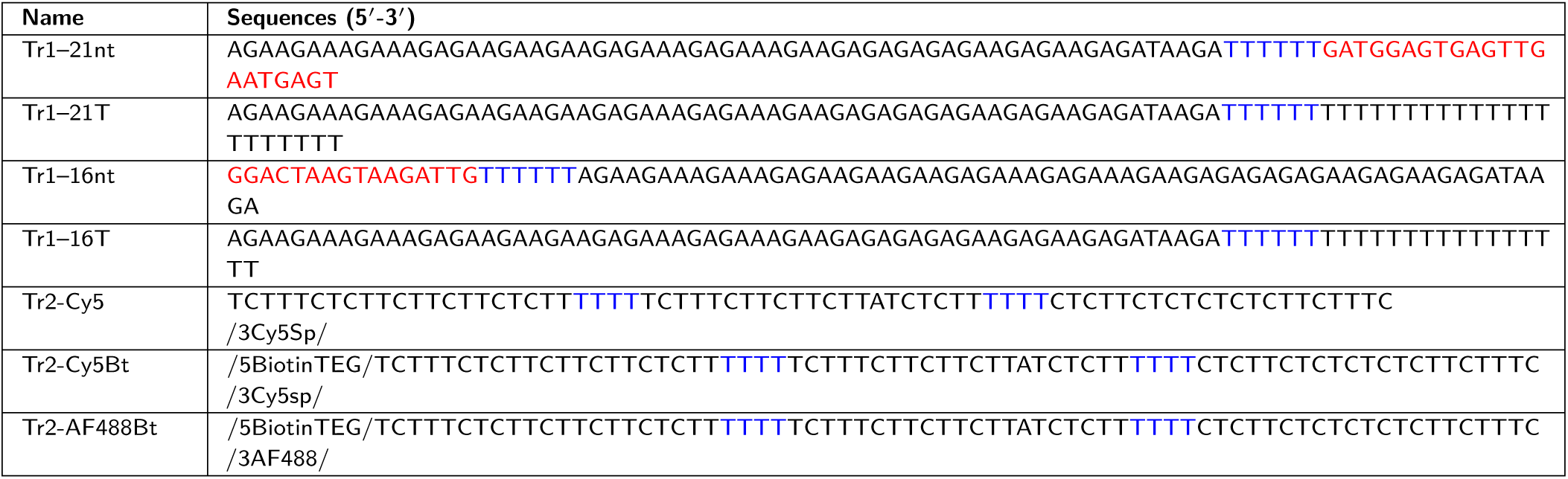
Triangle reporter strands. DNA sequences used for the formation of triangle reporters. Reporter domains are highlighted in red, polyT spacers at the triangle edges and linker between triangle and reporter recognition domain in blue. Cy5 modification at 3′ end is abbreviated as /3Cy5Sp/ and biotin modification at 5′ end is abbreviated as /5BiotinTEG/.

**Table S6.**
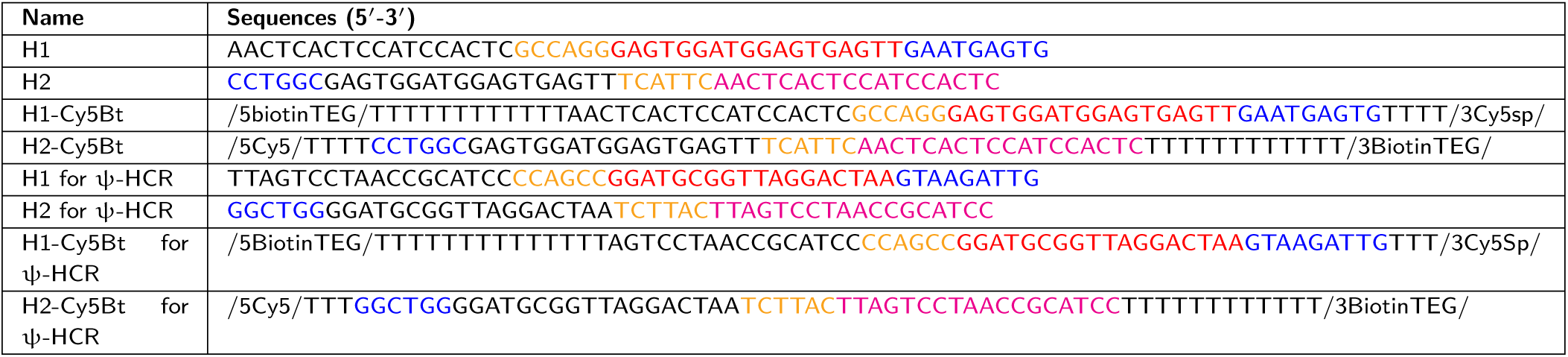
Metastable hairpin monomers. DNA sequences used for the formation of metastable hairpin monomers (H1, H2). Toehold of hairpin monomers are highlighted in blue, loop in yellow-orange, b*_p_* domain of H1 in red and b*_p_*\* of H2 in magenta, consistent with domain abbreviations of H1 and H2 in Fig. S14. Cy5 modification at 3′ end is abbreviated as /3Cy5Sp/ and biotin modification at 5′ end is abbreviated as /5BiotinTEG/.

**Table S7.** ESI-MS characterization of synthesized DNA and RNA conjugates. The in-house synthesized and HPLC purified DNA and RNA conjugates were characterized using ESI-MS using 0.1 % ammonium hydroxide buffer (v/v). Calculated and observed masses (Da) are presented in the tabular form.

| DNA strand name | Calculated mass (Da) | Observed mass (Da) |
| --- | --- | --- |
| HALOS 14 | Source: IDT |  |
| HALOS 21 | 20652.4 | 20652.6 |
| HALOS 25 | 23125.0 | 23125.1 |
| HALOS 28 | 24978.2 | 24979.5 |
| HALOS 31 | 26832.4 | 26832.7 |
| T-Cy3 | Source: IDT |  |
| T'-Cy3 | Source: IDT |  |
| Scr. T-Cy3 | Source: IDT |  |
| OMe-T-Cy3 (41-nt) | 13823.4 | 13822.2 |
| OMe-T' Cy3 (60-nt) | 20033.4 | 20031.9 |
| Target-E1 | 30680.2 | 30679.3 |
| Target-E2 | 30680.2 | 30680.5 |
| Target-E3 | 30680.2 | 30680.8 |
| R-Cy5 | Source: IDT |  |
| Scr. R-Cy5 | Source: IDT |  |
| R-AF488 | 6303.7 | 6303.8 |
| Tr2-Cy5 | 21748.4 | 21748.8 |
| Tr2-Cy5Bt | 22153.9 | 22154.5 |
| Tr2-AF488Bt | 22346.4 | 22346.2 |
| H1-Cy5Bt | 21610.6 | 21609.3 |
| H2-Cy5Bt | 21773.1 | 21773.6 |
| H1-Cy5Bt for ψ-HCR | 21201.3 | 21201.7 |
| H2-Cy5Bt for ψ-HCR | 20548.9 | 20549.6 |

**Table S8.** Compositions and number of atoms to construct the membrane. Table of lipid compositions studied for membrane modeling and measuring their physical parameters using MD simulations.

| Serial | Lipid composition | Total atoms |
| --- | --- | --- |
| System 1 | POPC | 30870 |
| System 2 | POPC + 30% Cholesterol | 29726 |
| System 3 | EGGPC | 31578 |
| System 4 | POPC + 30% LysoPC | 28188 |
| System 5 | EggPC + 10 % LysoPC | 28582 |

**Table S9.** HALOS variants and conformations studies using all-atom MD simulations. Table of HALOS variants and their conformations used for stability studies and signal transduction experiments using all-atom MD simulations.

| Serial | System composition | Total atoms |
| --- | --- | --- |
| System 1 | HALOS 14 in water | 47812 |
| System 2 | Axial orientation of HALOS 14 in POPC | 112539 |
| System 3 | HALOS 31 in water | 182795 |
| System 4 | Lateral orientation of HALOS 31 in POPC | 331478 |
| System 5 | Axial orientation of HALOS 31 in POPC | 199014 |
| System 6 | Intermediate conformation of HALOS 31 with target detection in POPC | 317758 |
| System 7 | Opened hairpin conformation of HALOS 31 with target detection in POPC | 425894 |

### Supplementary Figures

**Fig. S1.**
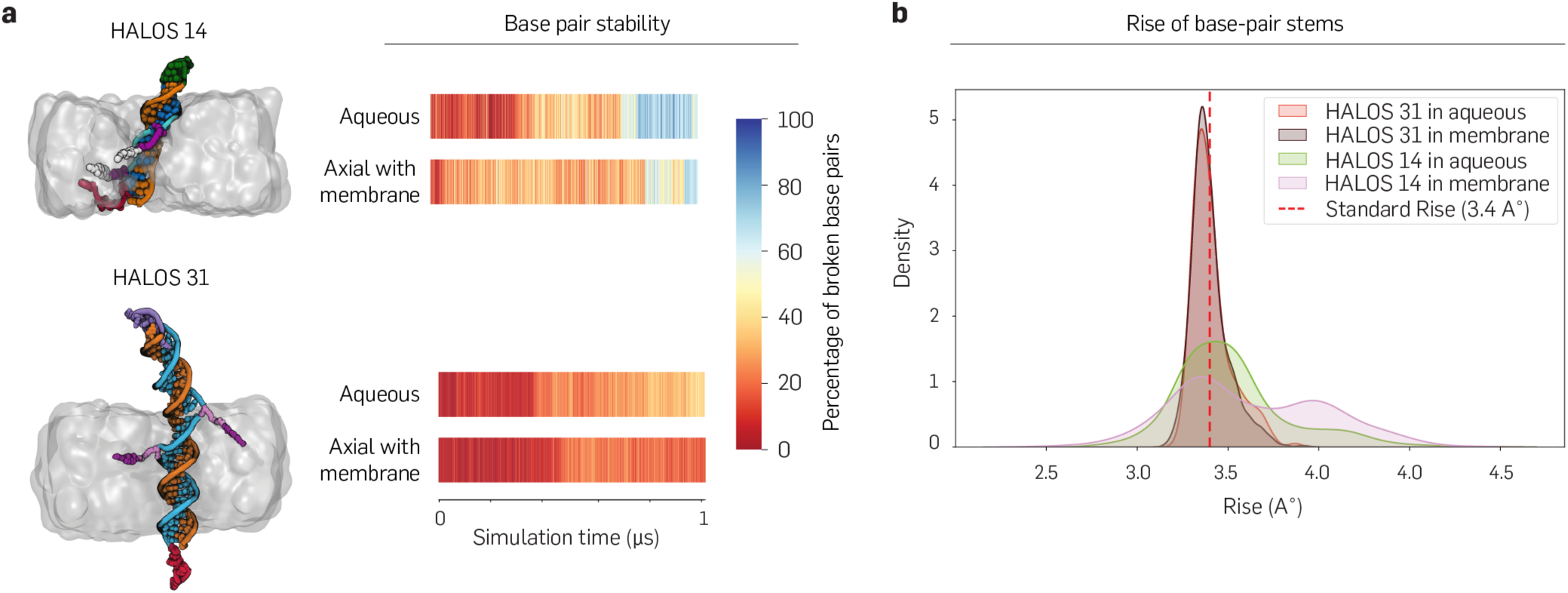
Stability assessment of HALOS-14 and HALOS-31 inserted in membrane using all-atom MD simulations. (**a**) Axial orientations of HALOS 14 and HALOS 31 within a POPC membrane. Heatmaps show the percentage of broken Watson-Crick base pairs in the paired stem domain during ∼1 µs simulations in aqueous solution and membrane environments. Base-pair disruption was quantified over time using hydrogen bond criteria (A-T: *<*2 H-bonds; C-G: *<*3 H-bonds; distance cutoff 3.4 Å, angle cutoff 45*^◦^*). HALOS 14 demonstrated greater base-pair disruption, indicating reduced structural stability compared to HALOS 31, suggesting HALOS 14 as a less suitable sensor variant. (**b**) Density distributions of the Rise base-pair step parameter reveal HALOS 31 maintains a sharp peak near 3.4 Å, consistent with canonical DNA spacing, while HALOS 14 deviates, reflecting structural distortion.

**Fig. S2.**
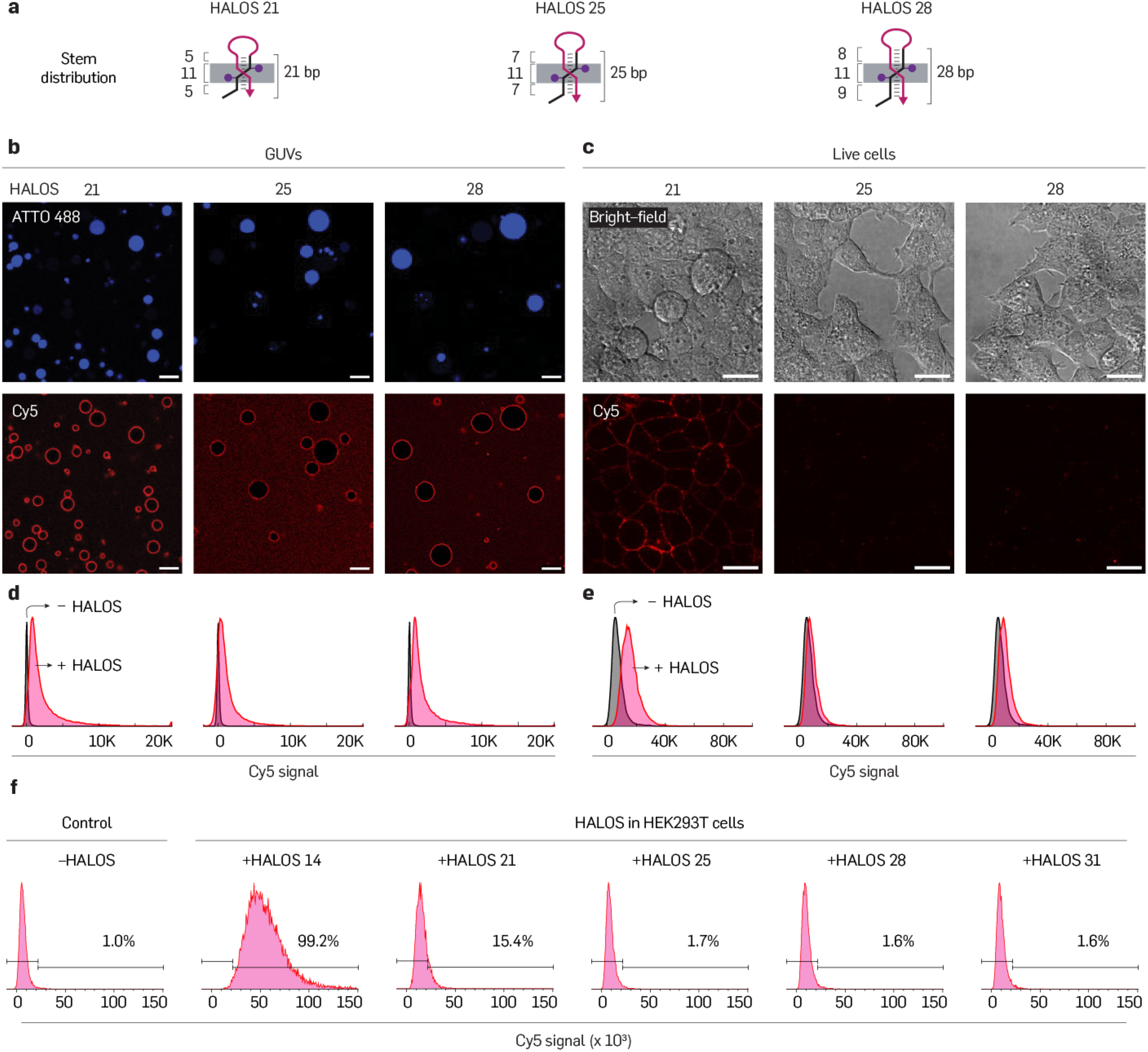
Additional confocal and flow cytometry analysis for HALOS variants in GUVs and live cells. In addition to Fig. 2**(c–d)**, the membrane stability of HALOS variants with varying stem-length (21–28 bp) was evaluated in GUVs and live HEK293T cells. (**a**) Schematic representation of HALOS variants differing in stem length and their membrane distribution in the axial orientation. (**b,c**) Confocal microscopy of HALOS stability in GUVs and live cells. HALOS (250 nM) was incubated with GUVs and live cells in the absence of an internal target (T-Cy3), followed by addition of reporter (R-Cy5, 100 nM) for 30 min. For imaging, GUVs were diluted five-fold in imaging buffer (1× TAE, 150 mM NaCl and 12.5 mM MgCl_2_), and cells were washed with Opti-MEM medium containing 12.5 mM MgCl_2_ and 50 µM ATA to remove unbound probes. ATTO488-encapsulated GUVs (in which the lumenal small-molecule dye marks target-free, intact vesicles) exhibited a gradual decrease in membrane Cy5 fluorescence with increasing stem length (21–28 bp), although HALOS 28 retained detectable signal intensity, indicating partial stem destabilization (b). In contrast, live cells showed significant Cy5 fluorescence for HALOS 21 but no detectable signal for HALOS 25–28, confirming improved membrane stability of longer-stem variants (c). Confocal images were acquired using 488 nm (ATTO488), 642 nm laser (Cy5) and halogen lamp (bright-field) under live-cell conditions (37 *^◦^*C, 95% humidity, and 5% CO_2_). The variation in the background of the confocal images in (b) and (c) arises from differences in the washing of excess reporter molecules and does not reflect the membrane-bound signal. (**d,e**) Flow cytometry of respective GUVs (d) and HEK293T cells (e) with and without (control) HALOS indicating similar trend of false-positive signal generation. This variation in the background does not affect the flow cytometry data, since flow cytometry measures the fluorescence associated with individual vesicles or cells and is not influenced by the fluorescence intensity of the surrounding media or buffer. (**f**) Quantification of HEK293T cells labeled with R-Cy5 (13-nt) upon HALOS treatment, depicting false-positive signals. Scale bar: 10 µm (b) and 20 µm (c).

**Fig. S3.**
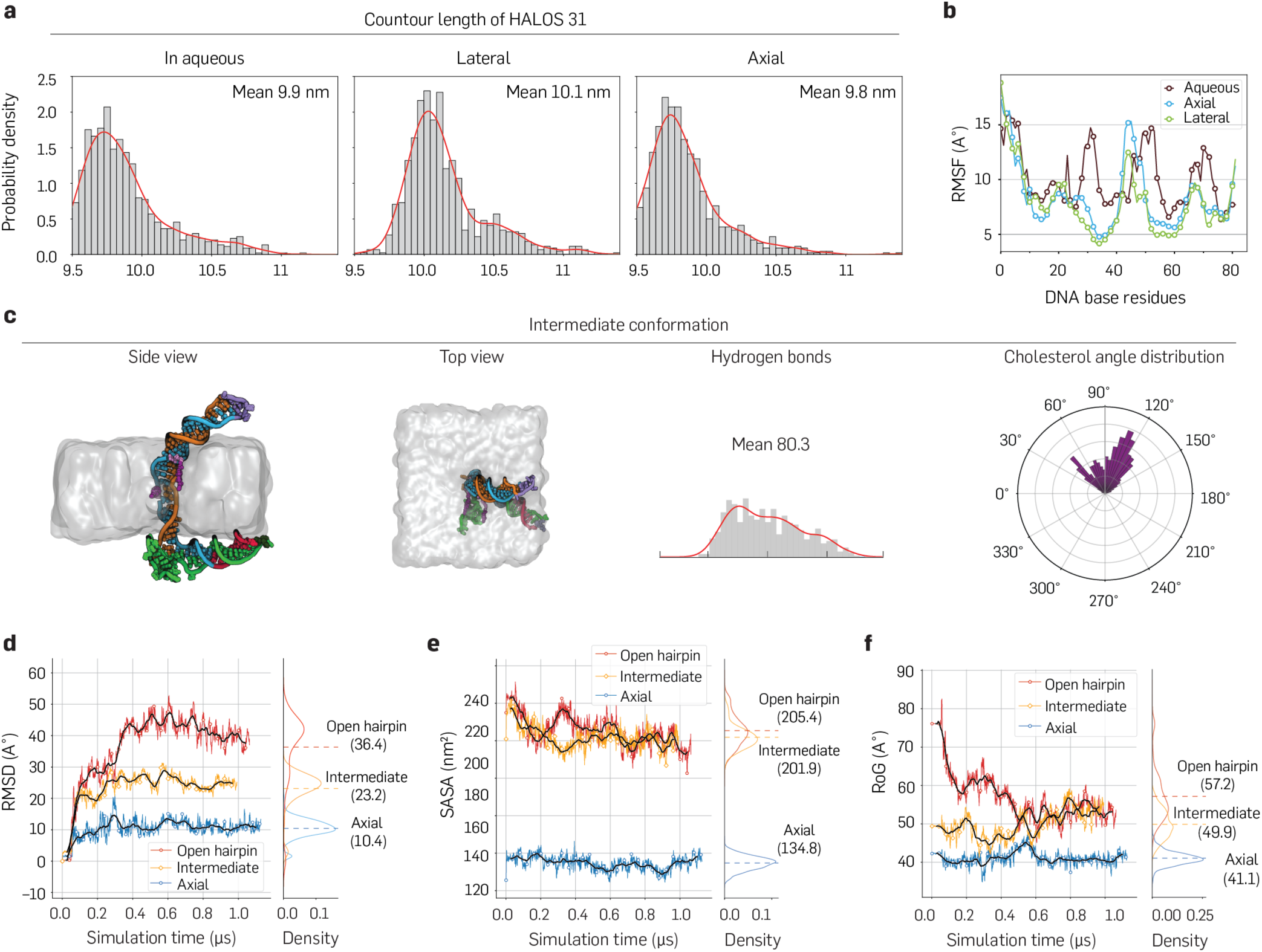
Additional MD simulations reveal membrane stability and signal transduction capability of HALOS and intermediate conformations. (**a**) Effect of the surrounding environment on the contour length of the paired region of the HALOS 31 stem, indicating less distortion in base pair stability for axially anchored HALOS compared to lateral and in aqueous solution. (**b**) Per-residue RMSF profiles of HALOS 31 demonstrate higher stability in the membrane compared to aqueous solution. (**c**) Representative intermediate structure shown from side and top views, along with Watson-Crick hydrogen bond patterns and cholesterol orientation distribution. (**d**–**f**) Comparative structural metrics for Axial, Intermediate, and Open hairpin conformations: RMSD (d), solvent-accessible surface area (SASA) (e), and radius of gyration (R*_g_*) (f). The Axial conformation, with the fewest unpaired bases, exhibits the lowest RMSD, SASA, and R*_g_* values, indicating the least deviations and compact structure in comparison to the other two.

**Fig. S4.**
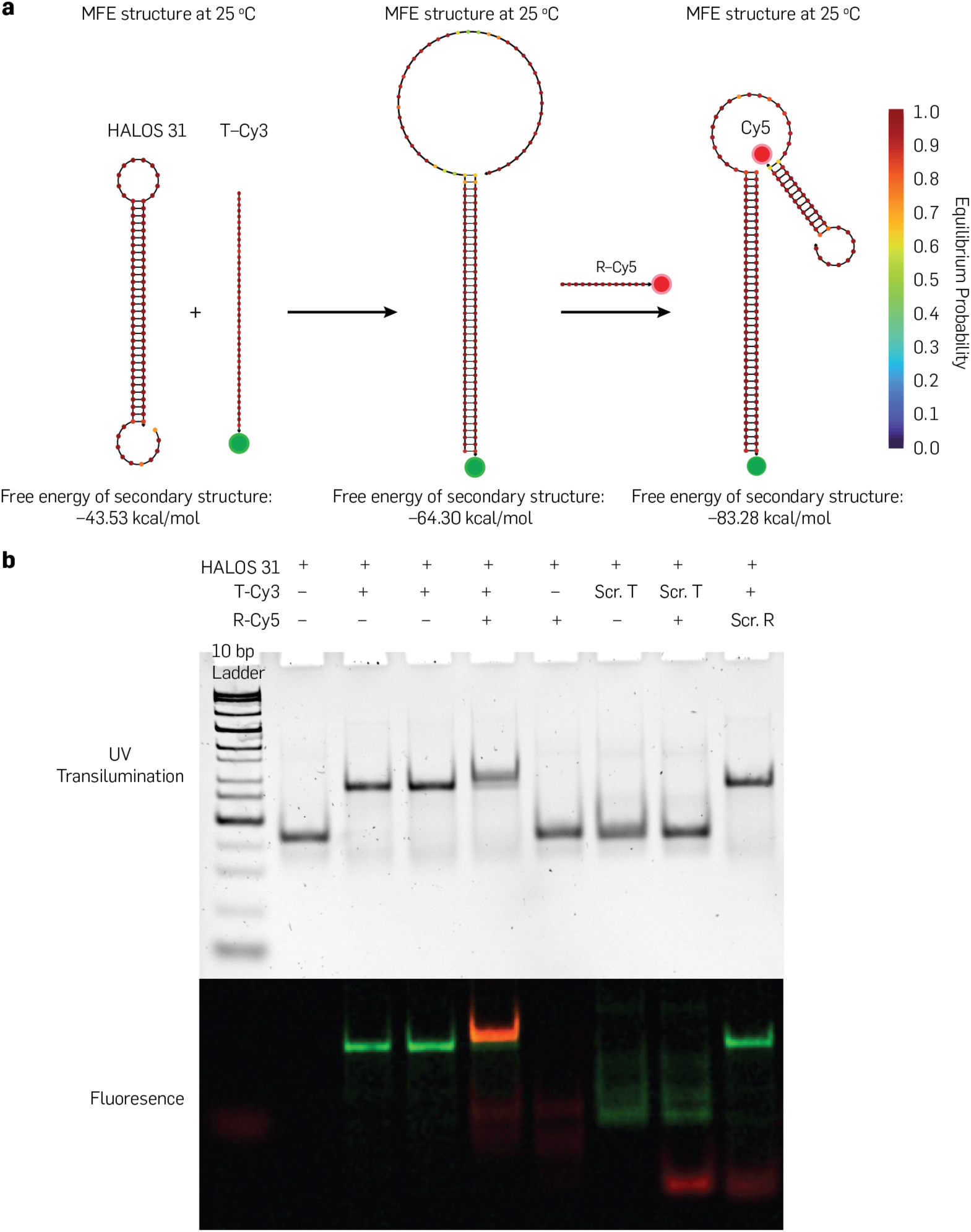
NUPACK simulation and native PAGE analysis of HALOS-mediated signal transduction in solution. HALOS-mediated target recognition, hairpin opening, and reporter binding in solution were examined using NUPACK simulations ^55^ and native PAGE. (**a**) NUPACK-simulated structures of HALOS 31 resulting from toehold-mediated T-Cy3 target binding, hairpin opening, and subsequent R-Cy5 recognition at 25 *^◦^*C. Sequences were designed for thermodynamic stability. Target (T-Cy3) binding initiated at the toehold domain of HALOS and resulted in complete hairpin unfolding, enabling reporter (R-Cy5) hybridization. (**b**) HALOS (without cholesterol anchor) was annealed and incubated with an equimolar T-Cy3 for 15 min at room temperature, followed by R-Cy5 addition. A 10% native PAGE was run in 1× TBE buffer with 12.5 mM MgCl_2_ at 100 V for 45 min at room temperature. Post-staining with EtBr (0.5 µg/mL, 10 min) and UV-imaging revealed T-Cy3 binding as a mobility shift versus HALOS alone (lane 1), for both pre-annealed complexes (lane 3) and in situ detection (lane 4). R-Cy5 binding induced a further mobility shift with Cy5 fluorescence (lane 5). Without target (lane 6) or with scrambled target sequences (lane 7-8), no stem opening or reporter labeling was observed. Scramble reporter also failed to bind with correctly targeted open hairpin in solution (lanes 9).

**Fig. S5.**
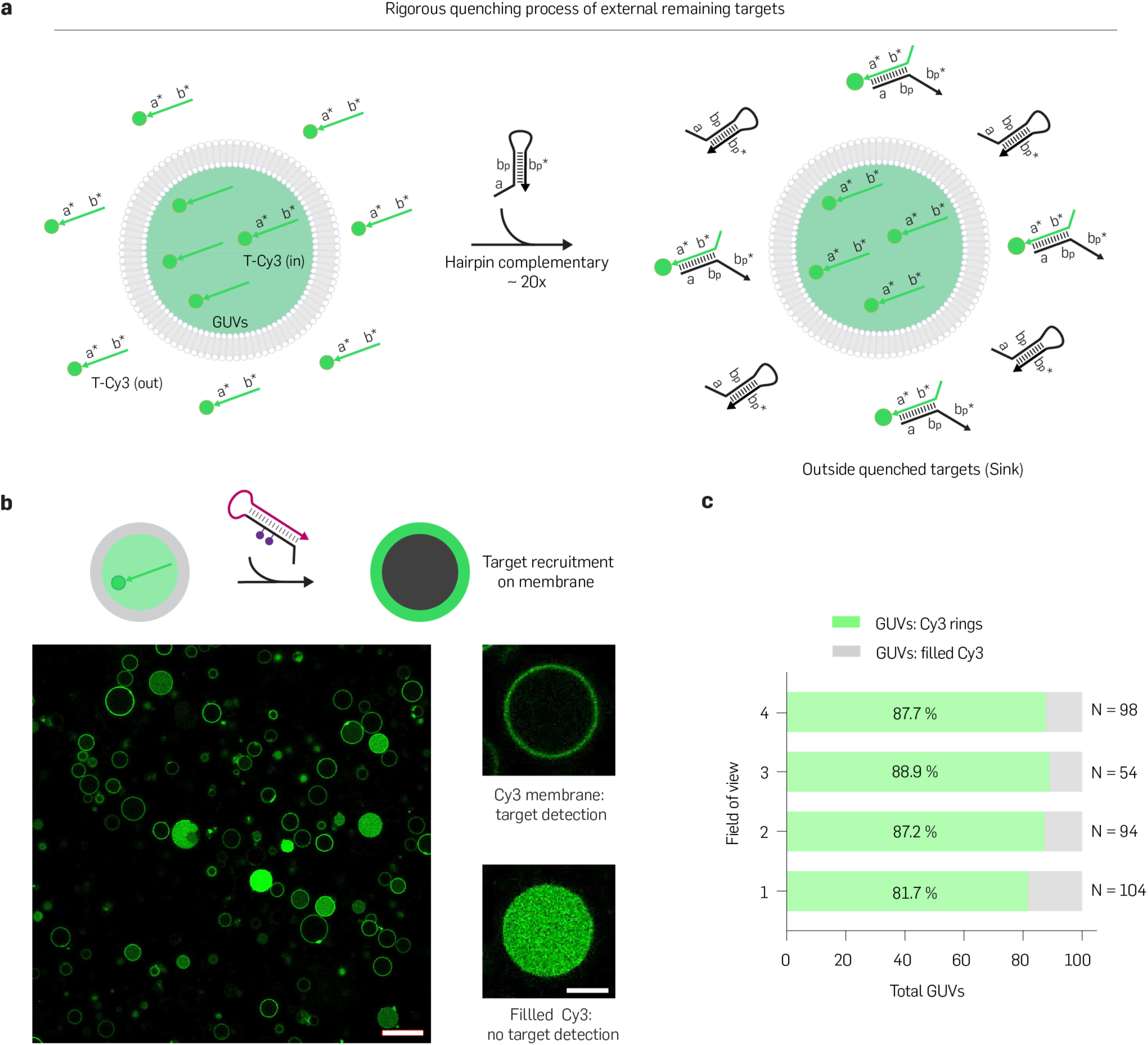
Quenching of extravesicular targets and quantification of HALOS mediated target recognition. (**a**) Schematic illustration of quenching free extravesicular targets by incubation with complementary hairpin strands (∼20× molar excess) for 45–60 min at room temperature. (**b,c**) Quantification of HALOS 31-mediated target detection across GUV membranes. T-Cy3 (250 nM) was encapsulated within GUVs (POPC) using electroformation, and excess external T-Cy3 was removed by centrifugation followed by treatment with complementary DNA (5 µM, ∼20× excess). Resulting GUVs were incubated with HALOS 31 (250 nM) for 120 min at room temperature and analyzed by confocal microscopy (561 nm laser, Cy3). (b) The confocal images show Cy3 fluorescence localized to the membrane upon successful HALOS 31 target recognition, while failed recognition resulted in uniformly fluorescent interiors. (c) Detection efficiency was quantified as the percentage of Cy3 ring-like fluorescence patterns (86 ± 1.9%, N_GUV_=350 across four fields of view). Data are mean±SD, Error bars are from bootstrapping. Scale bar: 20 µm (b), 5 µm (cropped images from b).

**Fig. S6.**
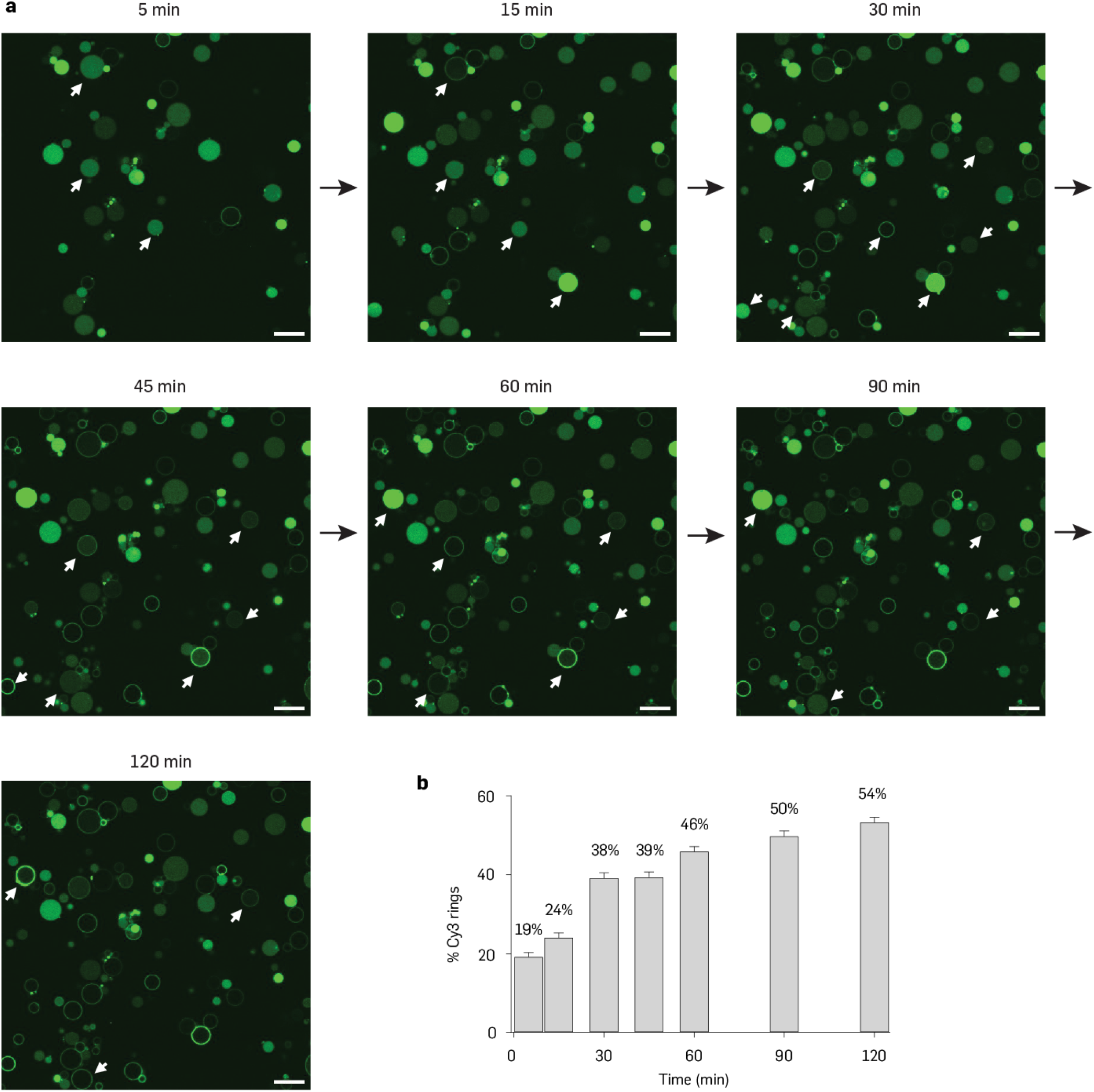
Kinetics of HALOS 31-mediated target detection across GUV membranes. (**a**) Cy3-labeled target detection using HALOS 31 was monitored by tracking Cy3 ring formation over time across lipid bilayers. Complementary DNA treated GUVs were plated onto microscopy dishes, and time-lapse confocal imaging was initiated immediately after HALOS 31 addition. Time-lapse images were acquired for up to 120 min, and the percentage of GUVs exhibiting Cy3 ring patterns was calculated at each time point. Target detection appeared in a fraction of GUVs within 5 min, with additional events accumulating over time. (**b**) Quantification across over three fields of view showed a gradual increase in Cy3 ring formation from 19% at 5 min to 54% at 120 min. Error bars are from bootstrapping. Scale bar: 20 µm (a).

**Fig. S7.**
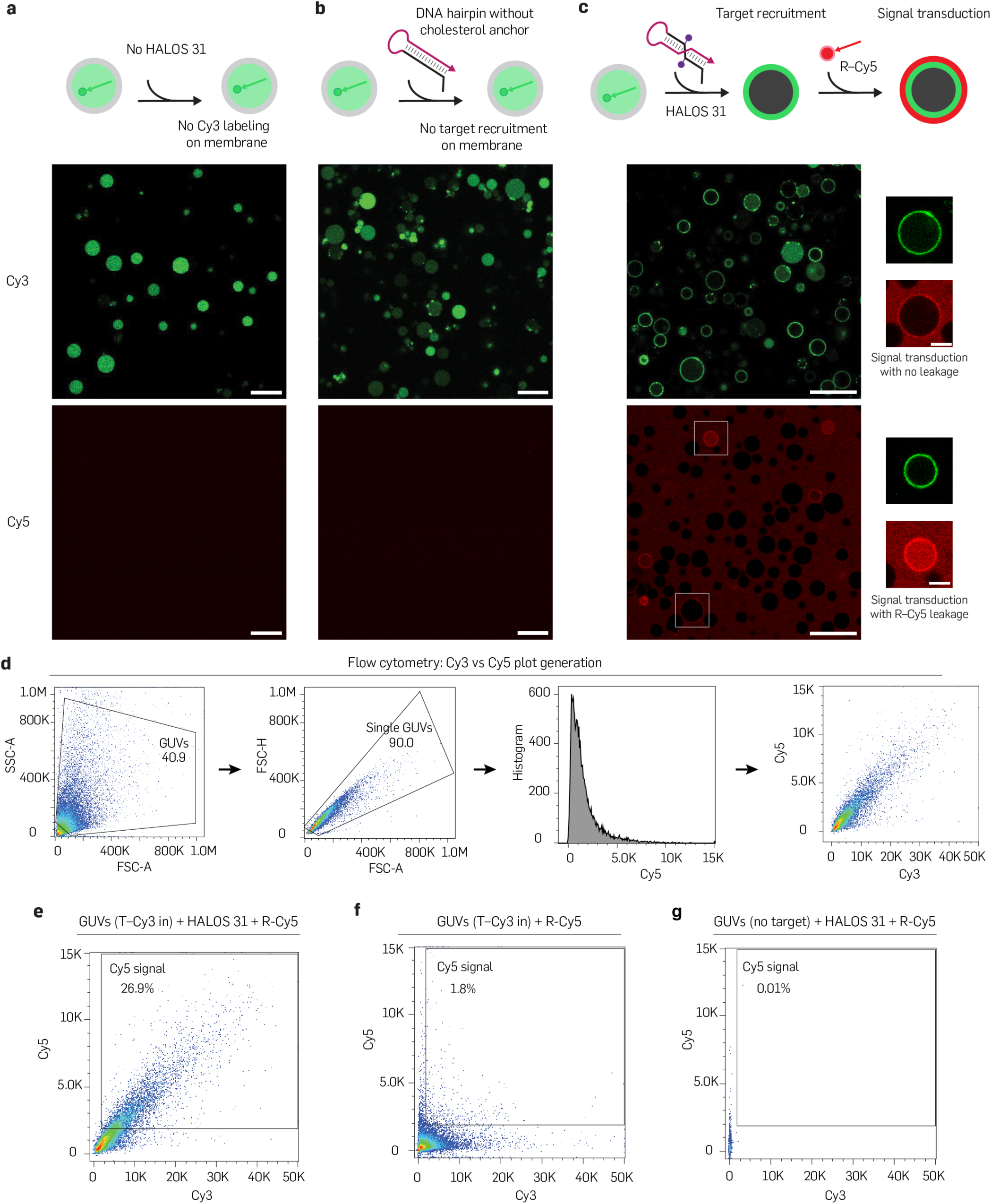
Additional large field of view confocal images and flow cytometry workflow for HALOS-mediated signal transduction. (**a,b**) Schematic and confocal images of controls lacking HALOS (a) or containing HALOS without cholesterol anchors (b), showing no Cy3 membrane labeling of T-Cy3 encapsulated GUVs and absence of reporter signal on membrane. (**c**) Extended confocal view corresponding to Fig. 3b, showing HALOS 31-mediated target recognition (Cy3) and signal transduction (Cy5). Most GUVs displayed membrane-associated Cy5 fluorescence consistent with signal transduction, while a minor population (*<*30%) exhibited R-Cy5 (13-nt ssDNA) diffusion across the membrane. (**d**) Schematic of flow cytometry analysis workflow used to generate histograms and Cy3 vs. Cy5 scatter plots. (**e**–**g**) Representative Cy3 vs. Cy5 plots quantifying signal transduction efficiency: 26.9% for HALOS-mediated samples (e), 1.8% for samples without HALOS (b), and 0.01% for GUVs lacking encapsulated targets (g). Scale bar: 20 µm (a–c), 5 µm (cropped images from c).

**Fig. S8.**
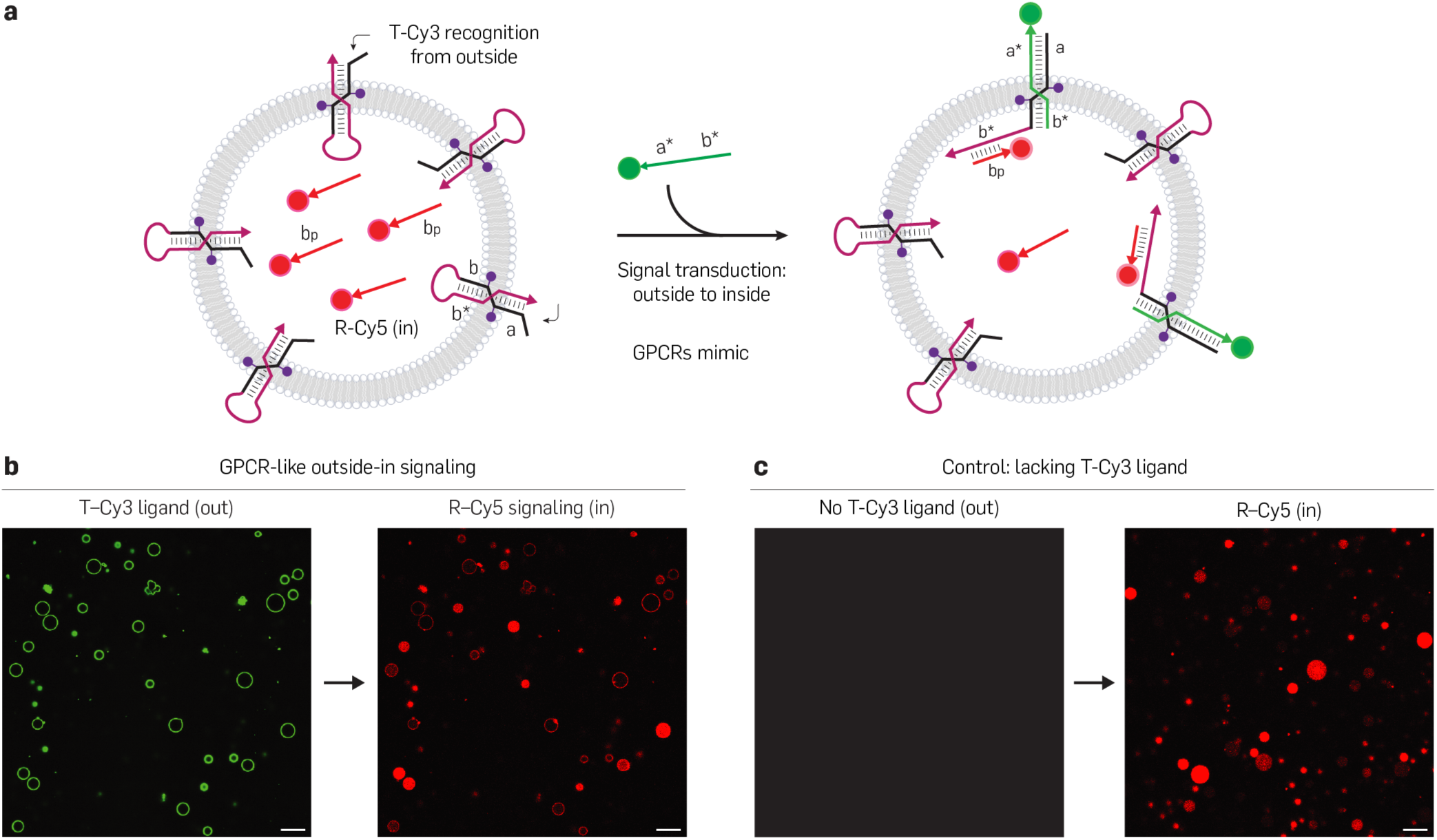
HALOS-mediated outside-in signaling analogous to GPCR signaling. (**a**) To recapitulate the outside-in directionality of GPCR signaling on synthetic vesicles, outside-in signal transduction was performed using HALOS 31. GUVs encapsulated with R-Cy5 were treated with HALOS 31 following quenching of external R-Cy5 by complementary DNA. Further, addition of T-Cy3 analogous to a GPCR ligand to the external medium allowed recruitment to the outward-facing toehold of HALOS 31, triggering hairpin opening and subsequent signal transduction by internal recognition of intravesicular R-Cy5. (**b**) Formation of Cy3 and Cy5 rings at the GUV membrane confirmed T-Cy3 recognition by HALOS, indicating successful signal transduction. (**c**) In the absence of external ligand, no Cy5 ring formation was observed, demonstrating no signal transmission. Scale bar: 20 µm (b–c).

**Fig. S9.**
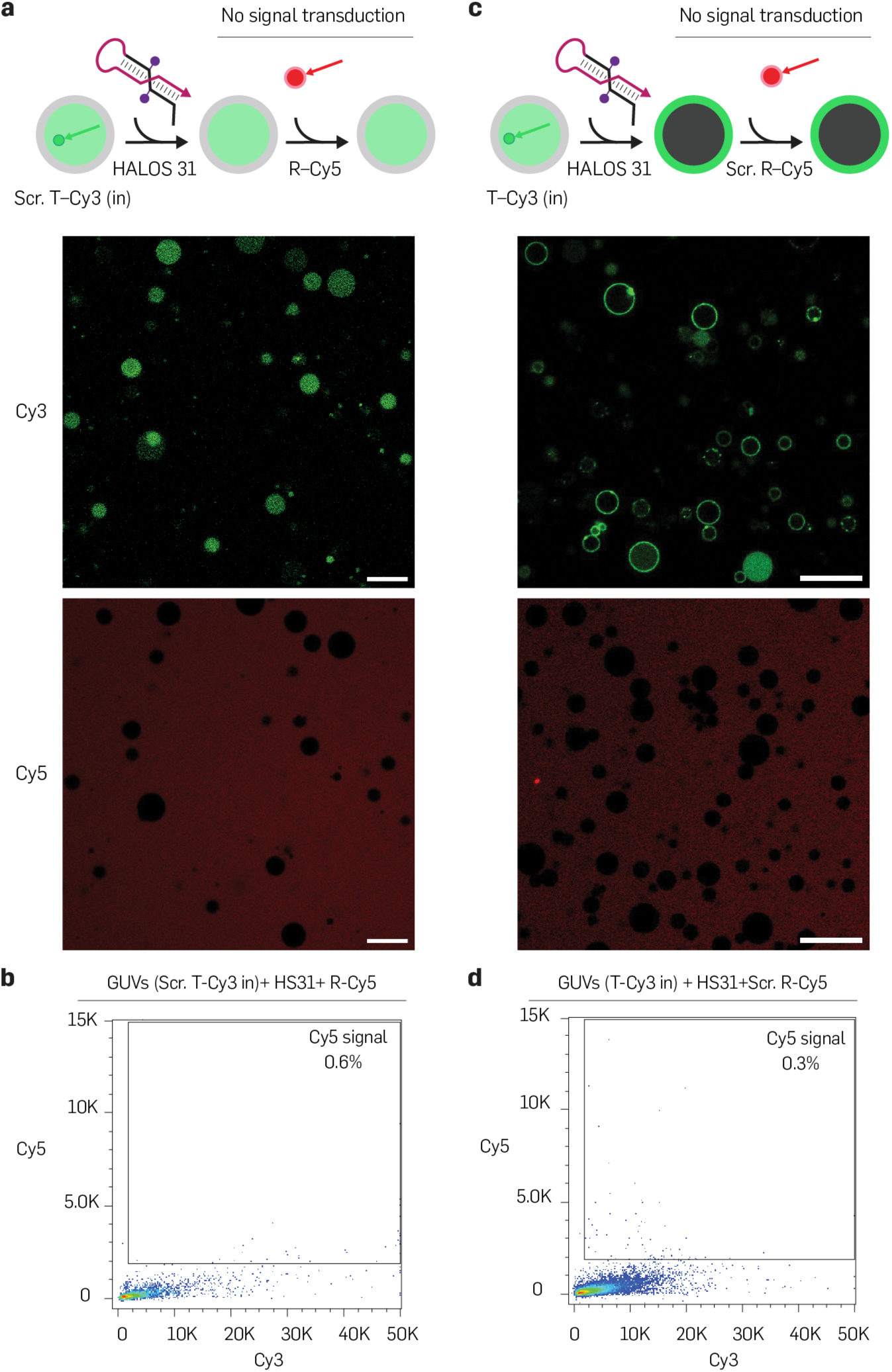
Additional large-field-of-view images of scramble controls. (**a**) Schematic and confocal images of GUVs containing scramble targets (Scr. T-Cy3, 250 nM) treated with HALOS 31 show no Cy3 ring formation on membrane. Addition of 13-nt ssDNA reporter (R-Cy5, 100 nM) resulted in no Cy5 ring formation, confirming no signal transduction across the membrane. (**b**) Cy3 vs. Cy5 flow cytometry plots reveal minimal signal transduction efficiency (0.6%) across the membrane for scramble controls. (**c,d**) GUVs encapsulating sequence-specific targets (T-Cy3, 250 nM) form Cy3 rings upon HALOS 31 treatment, but display no membrane Cy5 labeling in confocal microscopy and minimal efficiency (0.3%) in flow cytometry with polyT scramble reporter (100 nM), indicating sequence-specificity in HALOS-mediated signal transduction. Scale bar: 20 µm (a,c).

**Fig. S10.**
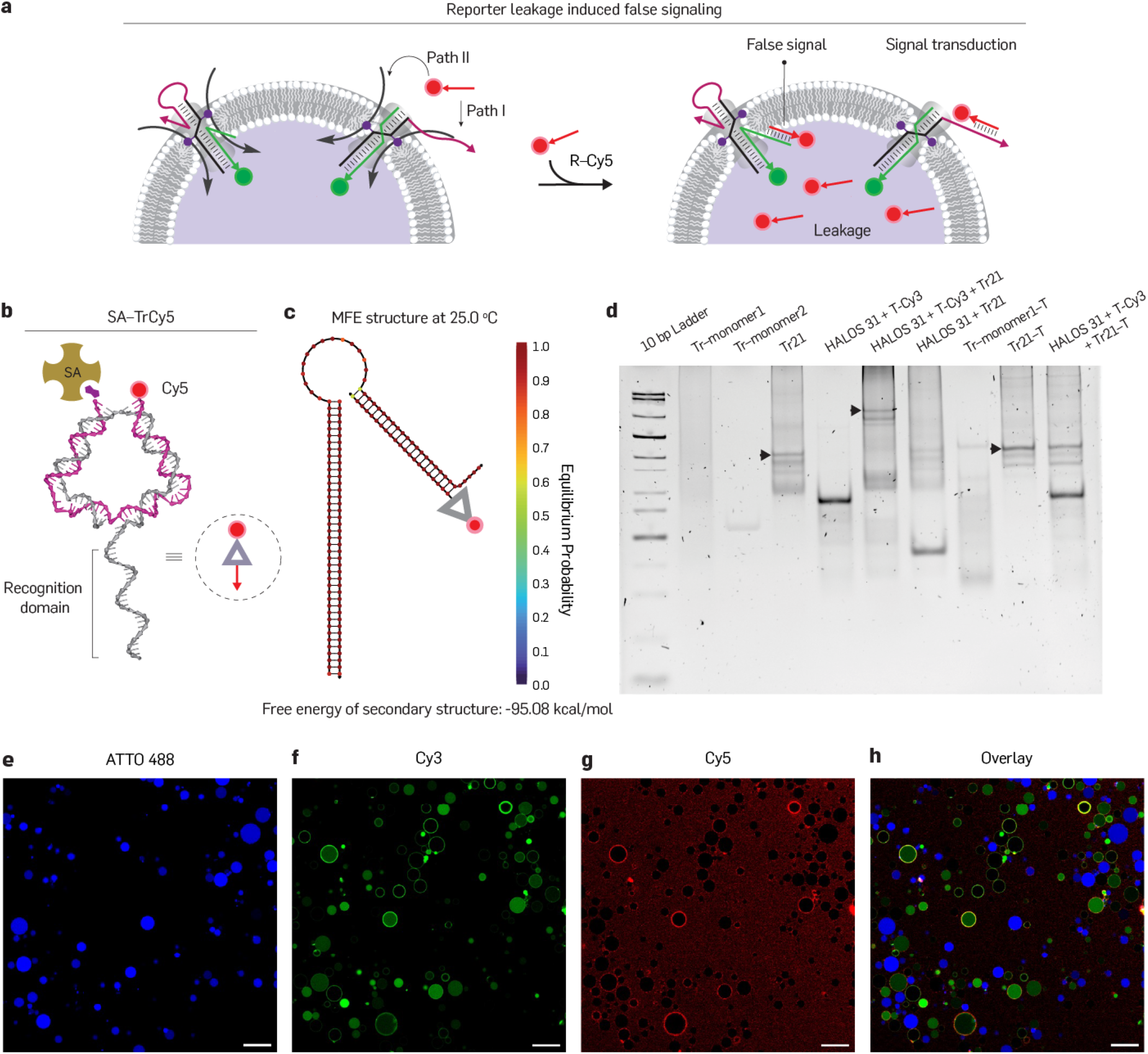
Bulkier reporter design for leakless signal transduction. (**a**) Schematic of gradient-driven leakage of the 13-nt ssDNA reporter (R-Cy5) across the membrane during signal transduction, where a transient, HALOS-induced membrane defect–which we attribute to a putative toroidal lipid nanopore, by analogy to previously reported cholesterol-anchored DNA nanostructures ^36,48,57^–can permit reporter diffusion and false-positive nonspecific labeling of overhanging T-Cy3. The size-dependent suppression of this leakage by progressively bulkier reporters (panels b–h) is consistent with diffusion through a transient, size-restricted aperture, independent of its precise geometry. (**b**) Design of a bulkier triangular reporter (Tr) formed by two complementary DNA strands extending reporter sequence to 21-nt, labeled with Cy5 and biotin for streptavidin (SA) anchoring to increase steric bulkiness. (**c**) NUPACK simulation confirms thermodynamic stability of Tr binding to open HALOS conformation. (**d**) Native 10% PAGE analysis shows formation of Tr reporter complexes with slower mobility than monomers and further shifted bands upon binding to open conformation of HALOS. Specificity in binding was confirmed by control interactions with HALOS alone and scramble reporter sequences. (**e**–**h**) Additional large-field confocal images (presented in Fig. 4d) of SA-TrCy5 signaling from T-Cy3 encapsulated GUVs show no reporter leakage inside GUVs. Control GUVs (ATTO488) show no signal transduction, confirming labeling specificity. Scale bar: 20 µm (e–h).

**Fig. S11.**
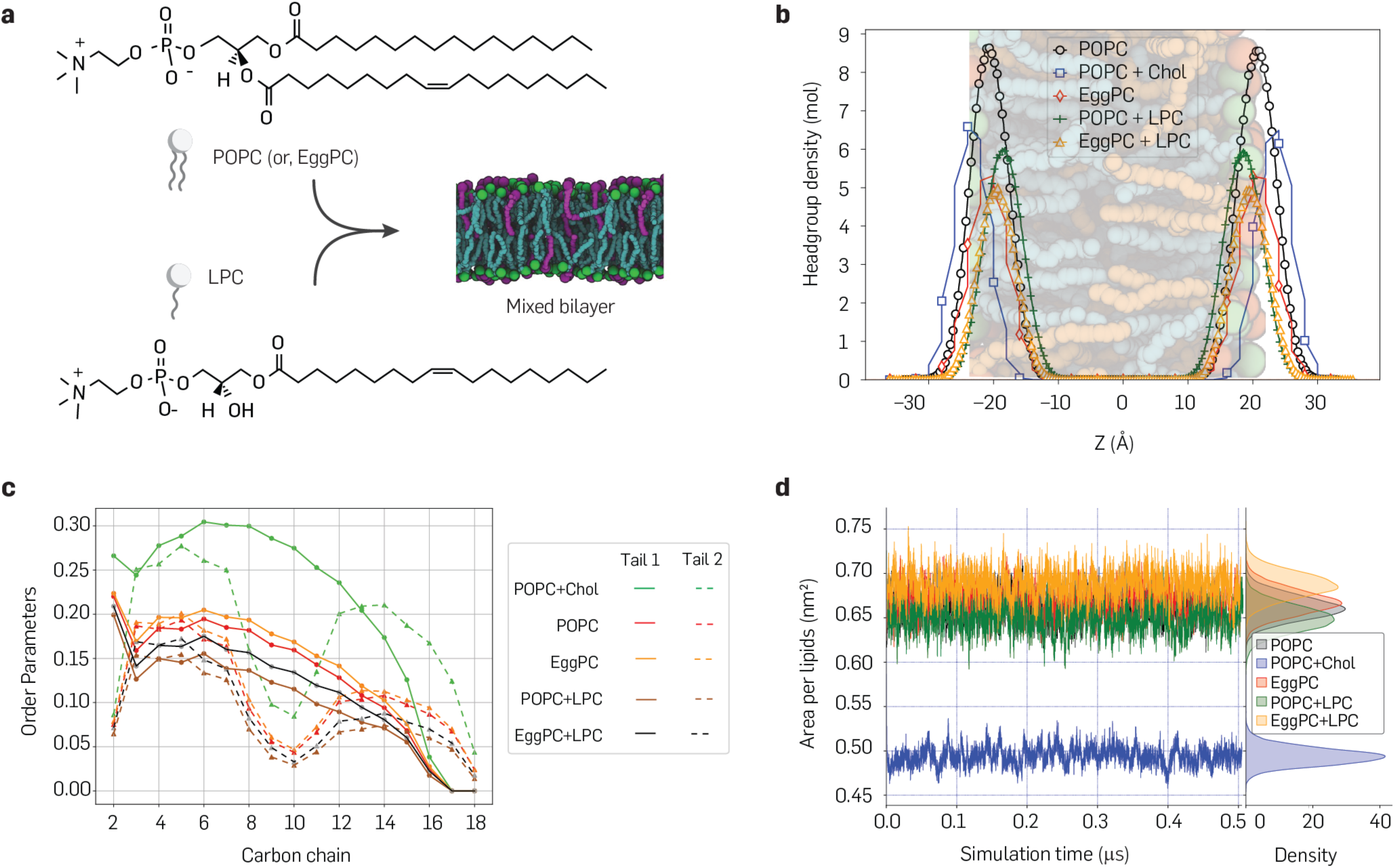
All-atom MD simulations reveal physical properties of lipid membranes. (**a**) Chemical structures of POPC (and/or, EggPC) and LPC, alongside an MD simulation snapshot of mixed POPC+LPC membrane. (**b**) Head group density profiles along the Z-axis of all lipid compositions studied, indicating membrane thickness. (**c**) Deuterium order parameters for the two acyl chains across all five models, reflecting membrane fluidity. (**d**) Area per lipid measurements for each membrane model, indicating membrane packing.

**Fig. S12.**
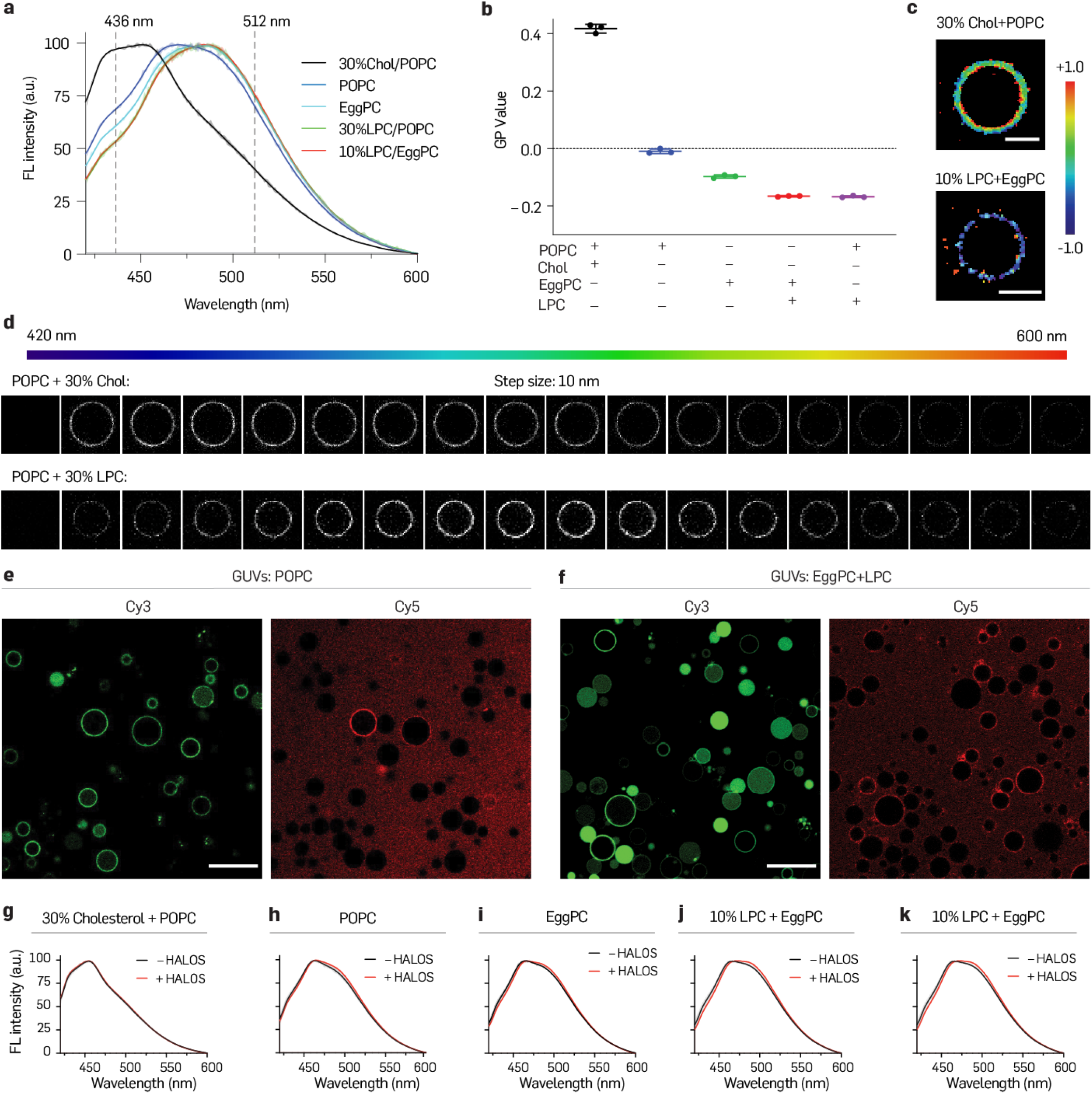
Membrane fluidity affects HALOS-mediated signal transduction in GUVs. (**a,b**) Membrane polarity was assessed by calculating Laurdan generalized polarization (GP) values. GUVs of various lipid compositions were incubated with Laurdan (5 µM for 15 min) and fluorescence spectra (a) were recorded to calculate GP values (b) by considering the fluorescence intensity at 436 nm (I_B_) and 512 nm (I_R_) upon excitation at 390 nm. Three replicates were measured for each lipid composition in this study. (**c**) GP images were generated from confocal images at these wavelengths using a custom ImageJ plugin, showing lipid-composition dependent GP shifts. (**d**) Spectral scanning of Laurdan indicated fluidity changes between (POPC+30% Chol) and (30% LPC + POPC) membranes; excitation: 405 nm, emission: 420–600 nm with 10 nm step size. (**e,f**) Confocal imaging of target recognition (Cy3) and signal transduction (Cy5) in GUVs of different lipid compositions demonstrated increased signal transduction with elevated membrane fluidity by higher LPC component. (**g**–**k**) Laurdan emission spectra of GUVs with and without HALOS (250 nM) confirmed membrane properties remain largely unchanged upon HALOS addition. Scale bar: 5 µm (c–d), 20 µm (e–f).

**Fig. S13.**
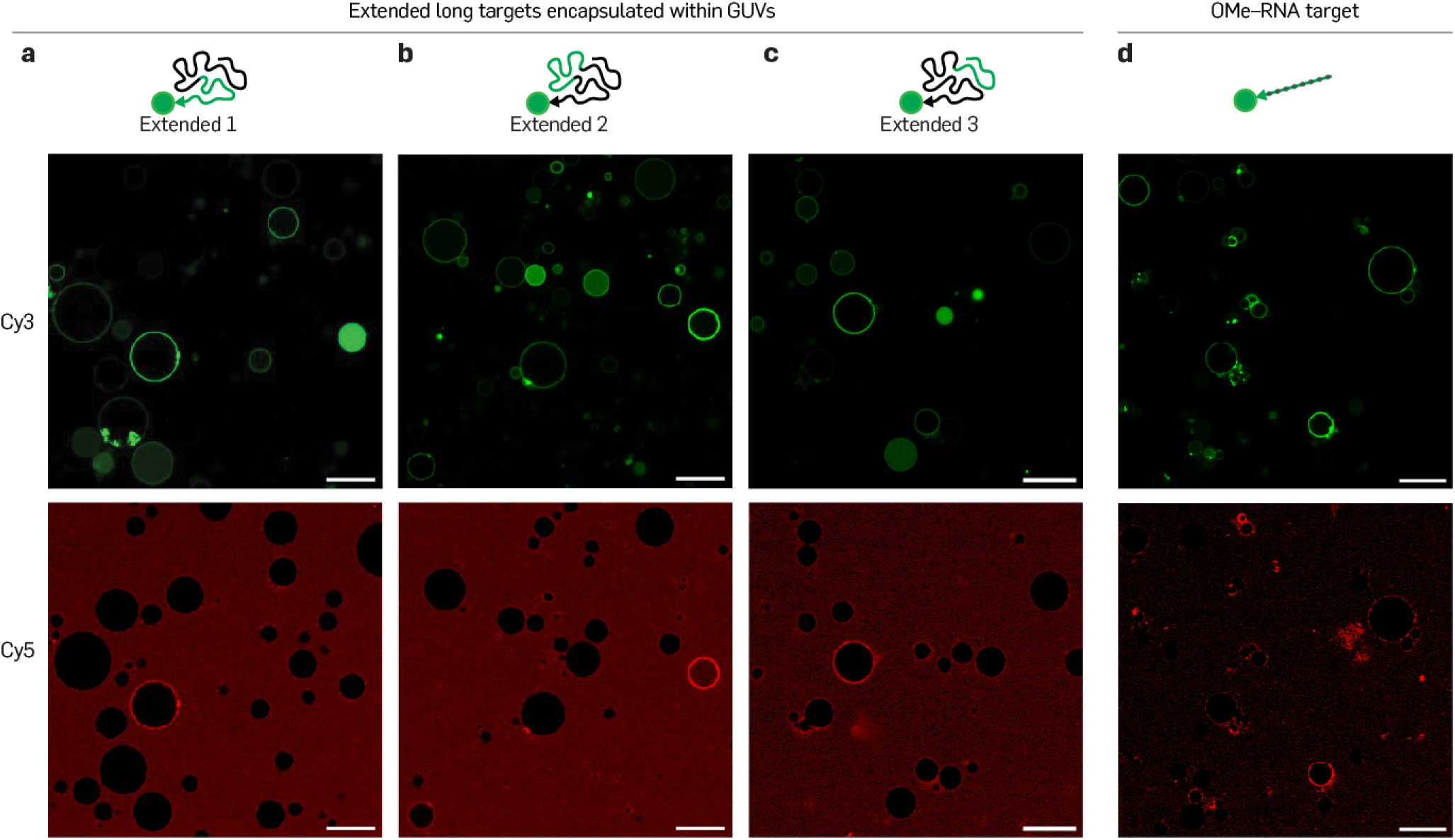
Additional large Field-of-view images of HALOS-mediated long extended-DNA and RNA-target recognition in GUVs. (**a**–**c**) Extended confocal-view corresponding to Fig. 5b. GUVs were encapsulated with longer DNA targets (100-nt, 250 nM) while keeping the recognition domain (41-nt) with polyT overhangs. Upon treatment with HALOS 31 (250 nM), all three isomers of long extended-targets (extended 1–3) showed Cy3 ring formation indicating target recognition independent of the position of the target recognition domain. Subsequent addition of Cy5-TrStp reporters (100 nM) yielded Cy5 ring formation, suggesting the signal-transduction ability of these long-DNA targets. (**d**) Encapsulation and detection of synthetic non-coding 2′-OMe-RNA (41-nt, 250 nM) in GUVs by HALOS 31, confirmed by Cy3 and Cy5 membrane ring formation, verifying efficient RNA target detection across the membrane. Scale bar: 20 µm (a–d).

**Fig. S14.**
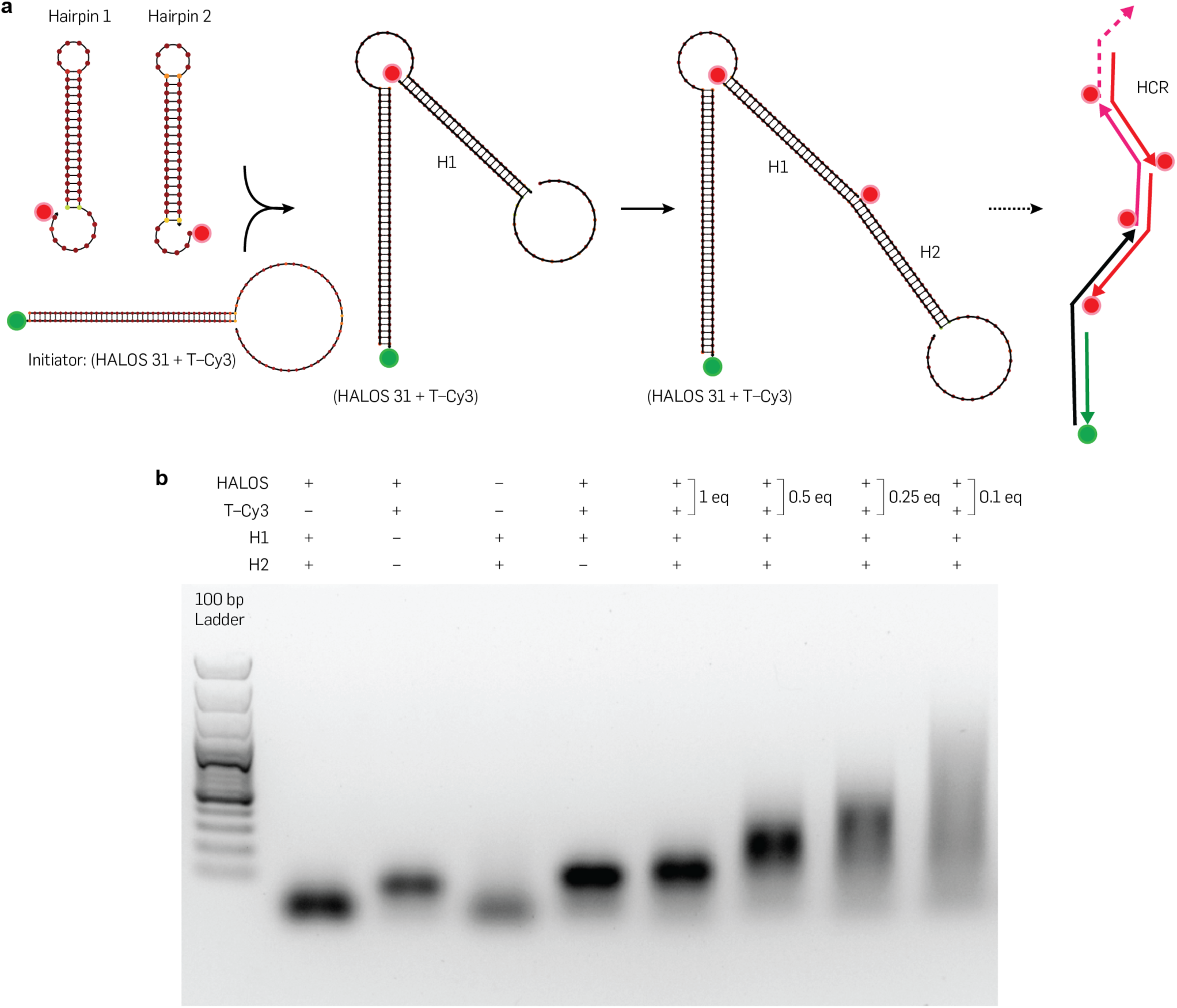
NUPACK simulation and Agarose gel analysis of HALOS-mediated hybridization chain reaction in solution. (**a**) Target-triggered hairpin opening in HALOS and initiation of hybridization chain reaction on open hairpin (initiator). NUPACK-simulated structure of the metastable hairpin monomers (H1, H2) coupled on HALOS, depicting spontaneous opening of the hairpin monomers in the presence of initiator and thermodynamic stability at 25 *^◦^*C. Sequences of H1 and H2 were designed with maximum thermodynamic energy gain with no cross-reactivity. (**b**) Agarose gel electrophoresis of HCR in solution (1× TAE+150 mM NaCl+12.5 mM MgCl_2_) supports the initiation of HCR with initiator (HALOS 31+T-Cy3). HCR progress was monitored with varied initiator ratios (1 to 0.1 equiv.), while no HCR was observed in the absence of T-Cy3 and HALOS 31. 1% (w/v) Agarose gel electrophoresis was performed using 1× Sodium Borate buffer (10 mM NaOH, pH 8.5) supplemented with 12.5 mM MgCl_2_. Gels were prestained with SYBR gold (0.5 µg/mL), loaded with 10 pmol (1 µM × 10 µL) sample and run with 100 V for 90 min under ice-cold condition and imaged by UV transillumination.

**Fig. S15.**
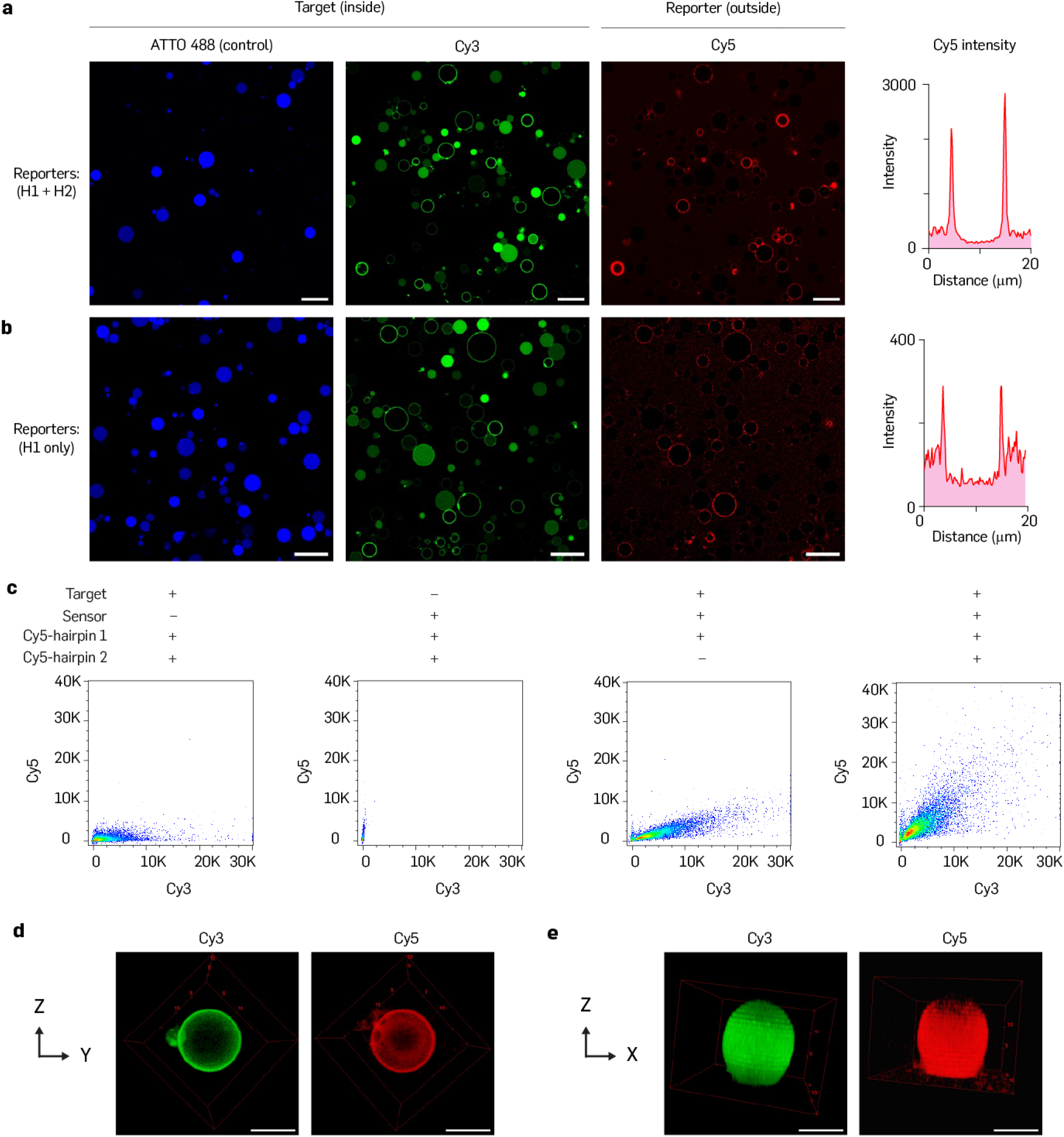
Hybridization chain reaction amplifies target-specific signal in GUVs. (**a** and **b**) Extended confocal views demonstrate HCR-based signal amplification from target-encapsulated (T-Cy3) GUVs, while control GUVs (ATTO488) show no amplification (presented in Fig. 6c). (a). Reporter H1 yields low signal-to-noise compared to HCR as illustrated by confocal images and Cy5 intensity profiles (b). (**c**) Cy3 vs. Cy5 scatter plots compare HCR reactions with H1 only and controls lacking HALOS 31 and target (T-Cy3), revealing higher Cy5 signal for HCR, compared to H1 alone and negative controls. (**d**–**e**) 3D view of GUVs after HCR on the membrane. GUVs containing Cy3 target, treated with HALOS 31 and HCR monomers, were subjected to 3D confocal microscopy over 15 µm depth with 500 nm z-stack resolution. 3D images were reconstructed using the ImageJ 3D viewer plugin. The homogeneous distribution of Cy3 and Cy5 labeling on GUVs was confirmed by different angles of view (YZ and XZ planes), confirming no non-specific labeling of HCR monomers to GUVs. Scale bar: 20 µm (a–b) and 5 µm (d–e).

**Fig. S16.**
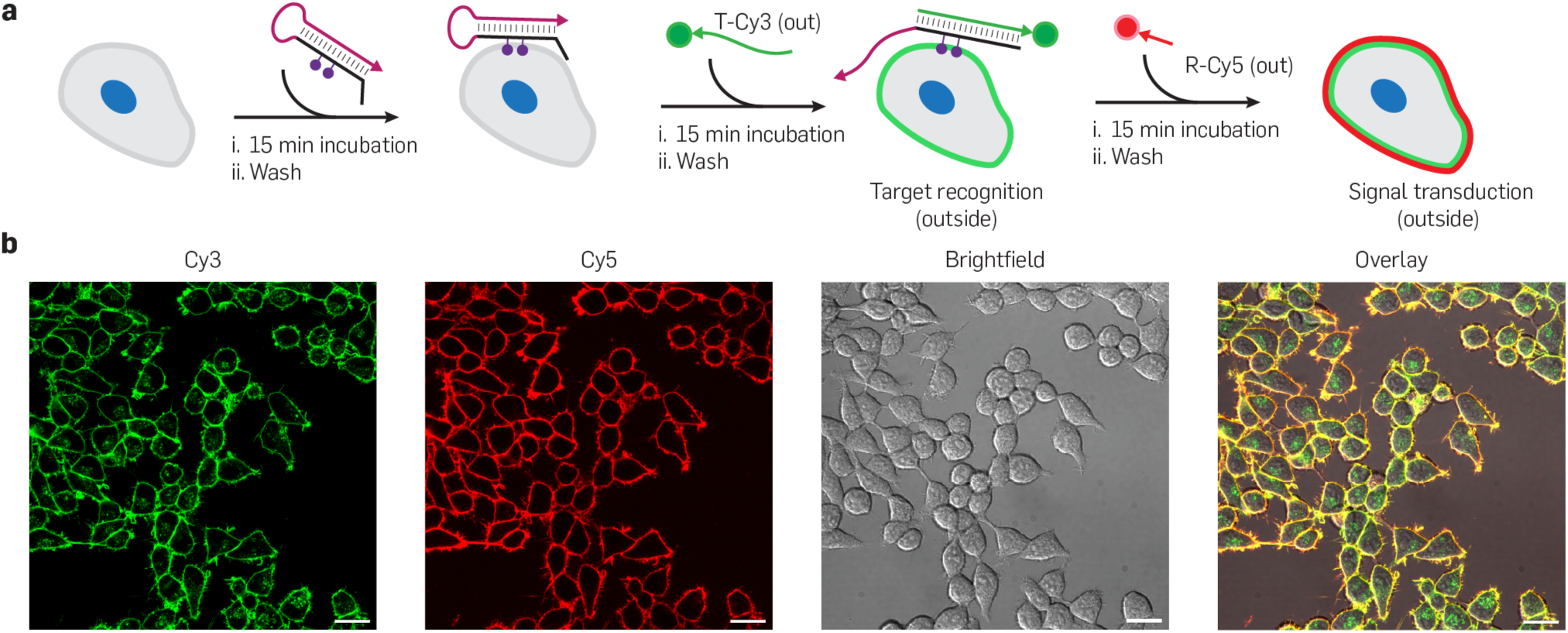
Evaluation of the live-cell compatibility of HALOS-mediated signal transduction on HEK293T cell membranes. (**a**) The compatibility of HALOS 31 constructs was assessed on live HEK293T cell membranes by performing signal transduction. Schematics illustrating the HALOS 31 anchoring (250 nM) in Opti-MEM containing 50 µM ATA and 12.5 mM MgCl_2_ for 15 min in a live-cell culture environment, followed by extracellular T-Cy3 recognition (250 nM), HALOS hairpin opening, and signal transduction by R-Cy5 labeling (250 nM) on the exposed opened hairpin domain on the cell membrane, with washing steps between each addition. (**b**) Confocal images showing overlaying Cy3 and Cy5 fluorescence at the cell membrane, demonstrating compatibility of HALOS-mediated target detection and signal transduction on the live-cell membrane. Scale bar: 20 µm (b).

**Fig. S17.**
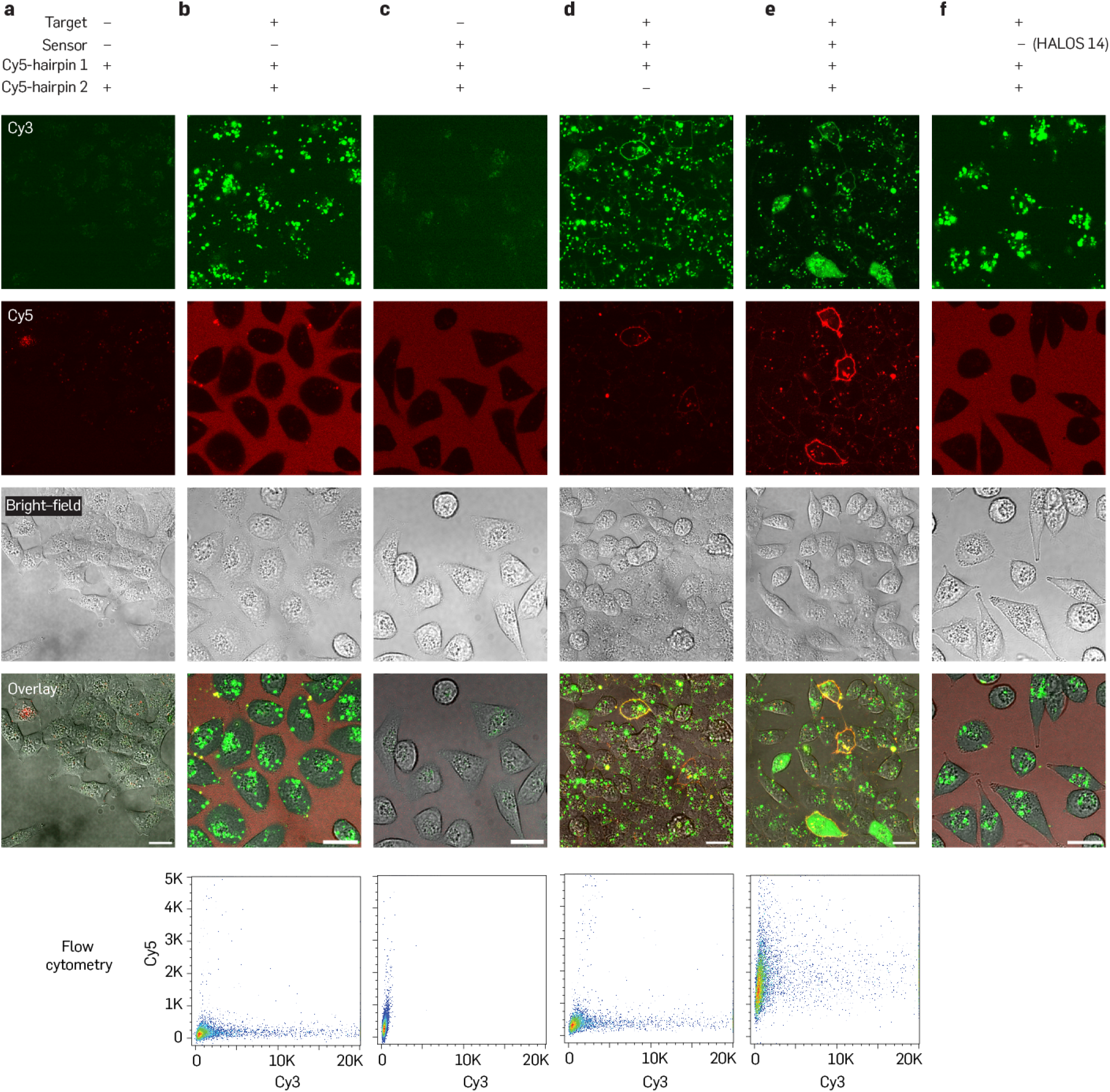
Signal amplification of HALOS-mediated RNA detection in live A549 cells. (**a**–**f**) Additional confocal microscopy images for signal amplification for HALOS-mediated RNA detection and amplification in live A549 cells (presented in Fig. 6e). Cells were transfected with synthetic 2′-OMe-RNA target (41-nt) using lipofectamine-based transfection. Negative controls lacking HALOS, target RNA, or both displayed no membrane Cy3 and Cy5 fluorescence (a–c). Target RNA-transfected cells treated with HALOS 31 exhibited a bright Cy5 signal upon performing HCR reaction (60 min in cell-culture environment at 37 *^◦^*C) with substantially higher Cy5 intensity for HCR versus H1 alone (d–e). HALOS having mismatched sequence (HALOS 14) with target RNA resulted in no signal transduction and amplification (f). Flow cytometry Cy3 vs. Cy5 scatter plots corroborate amplified Cy5 surface signal for HCR compared to H1 and negative controls. Scale bar: 20 µm (a–f).

**Fig. S18.**
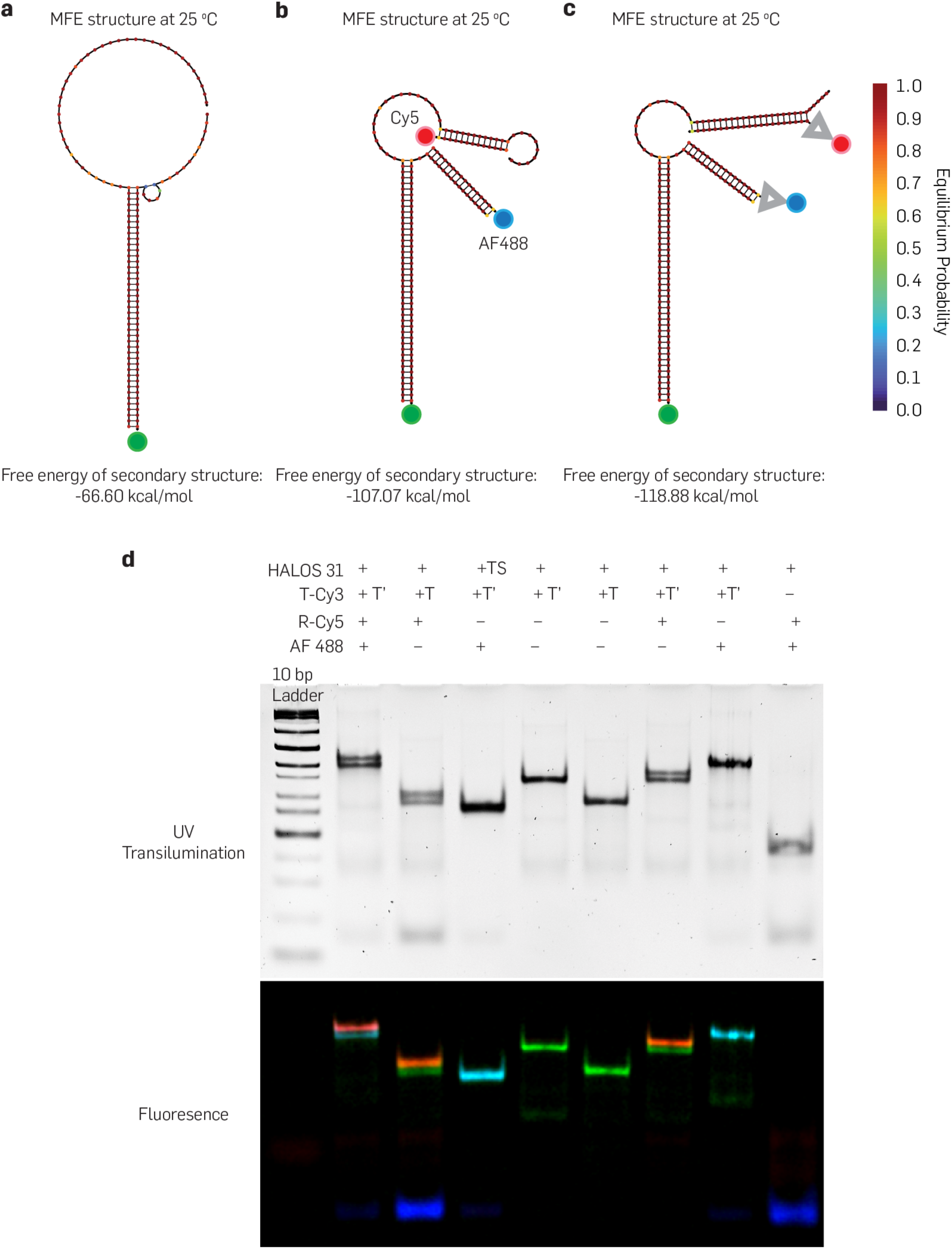
NUPACK simulation and native PAGE confirm dual reporter labeling of extended-target-treated HALOS 31 in solution. (**a**–**c**) NUPACK simulated structure of extended target (T′-Cy3, 60-nt) bound to HALOS 31 (a), linear and SA-Tr dual-reporter (Cy5 and AF488) labeling to the opened hairpins (b–c), confirming thermodynamic stability and no cross reactivity between reporters. (**d**) Native PAGE of dual reporter labeling to open HALOS conformation in solution. 10% Native PAGE was run using 1× TBE buffer + 12.5 mM MgCl_2_ at 100 V for 45 min at room temperature. Gels were post-stained with EtBr (0.5 µg/mL) for 10 min and imaged using UV-transillumination. Fluorescence images were scanned using an ImageQuant 800 gel imager. Addition of both R-Cy5 and R-AF488 resulted in dual labeling of T′-Cy3 (60-nt) treated HALOS 31 (lane 2). Absence of either the extended domain from the target (T-Cy3, 41-nt) or HALOS (lacking open hairpin domain) showed no labeling of R-AF488 and R-Cy5, respectively (lanes 3–4 vs lane 6), confirming no cross reactivity between R-Cy5 and AF488 reporters to open hairpin. The single color reporter (either R-Cy5 or R-AF488) labeling to T′-Cy3 (60-nt) treated HALOS 31 resulted in specific reporter labeling as depicted by the fluorescence image (lane 5 vs lanes 7–8). In the absence of target (T′-Cy3), HALOS 31 remained unreactive to both the reporters (lane 9).

**Fig. S19.**
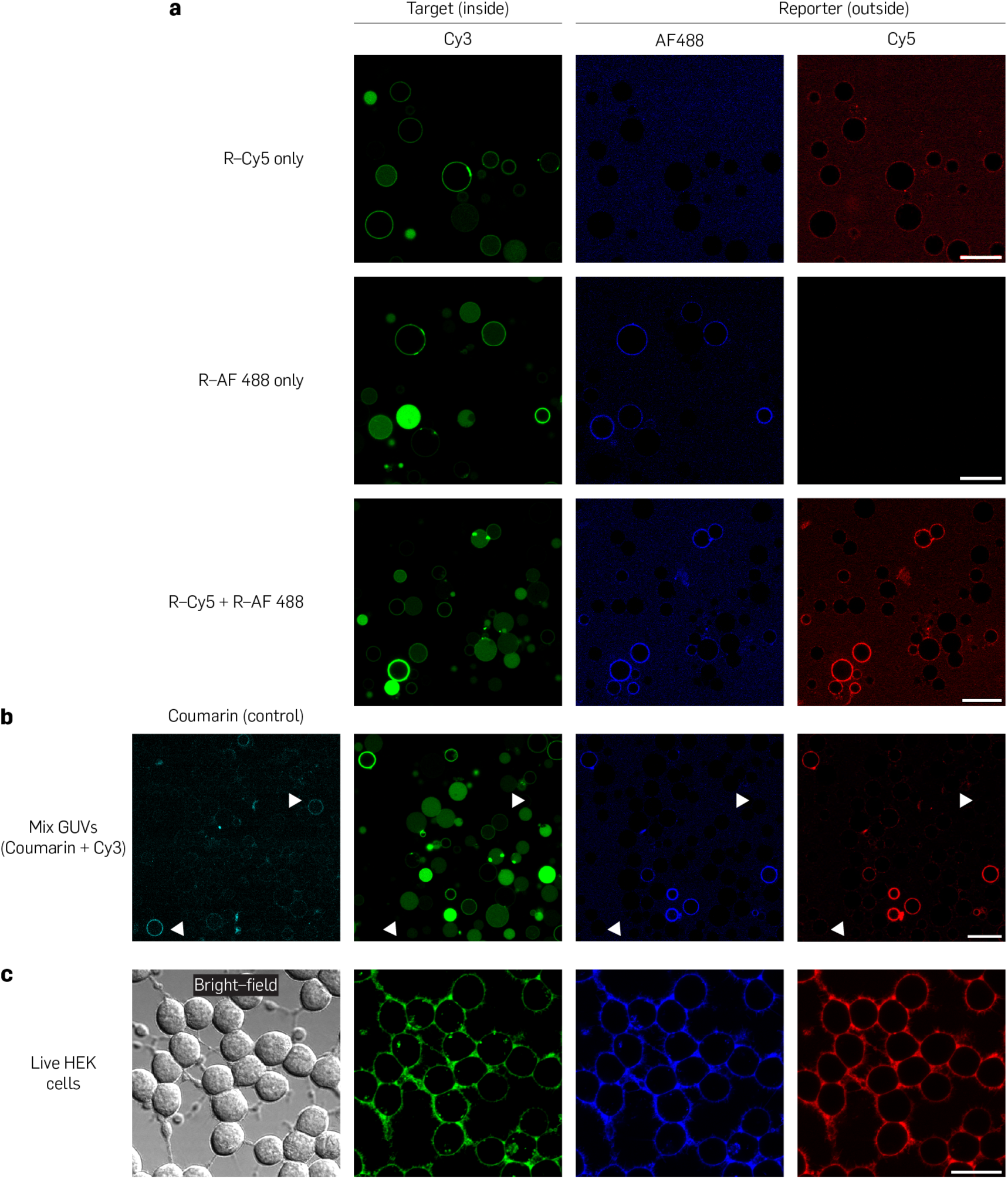
Additional dual reporter labeling images in GUVs and live-cell compatibility. (**a**) Extended-views of confocal images for dual reporter labeling in GUVs corresponding to Fig. 7c. When treated with R-Cy5 (100 nM) or R-AF488 (100 nM) individually, T′-Cy3 detected GUVs showed specific fluorescence on the respective Cy3 or AF488 channel; both were observed when both reporters were applied simultaneously. (**b**) Dual reporter labeling with a control coumarin dye (without target DNA) encapsulated GUVs (Cyan). Cy3 target GUVs display dual reporter labeling, while no reporter labeling is seen on coumarin GUVs, suggesting no non-specific signaling. Coumarin dye binds to lipid membrane showing cyan rings as control GUVs. (**c**) Live-cell compatibility of dual reporter probes. Live HEK293T cells were incubated with HALOS 31 (250 nM) in Opti-MEM containing 50 µM ATA and 12.5 mM MgCl_2_ for 15 min under live-cell culture environment. After washing with Opti-MEM, cells were sequentially treated with T-Cy3 (250 nM) and R-Cy5 + R-AF488 (250 nM each) with washing steps (3×) between each addition. Confocal imaging showed Cy3 (green), Cy5 (red) and AF488 (blue) fluorescence, demonstrating compatibility of HALOS-mediated signal transduction using dual reporter labeling on live-cell membrane. Scale bar: 20 µm (a–c).

**Fig. S20.**
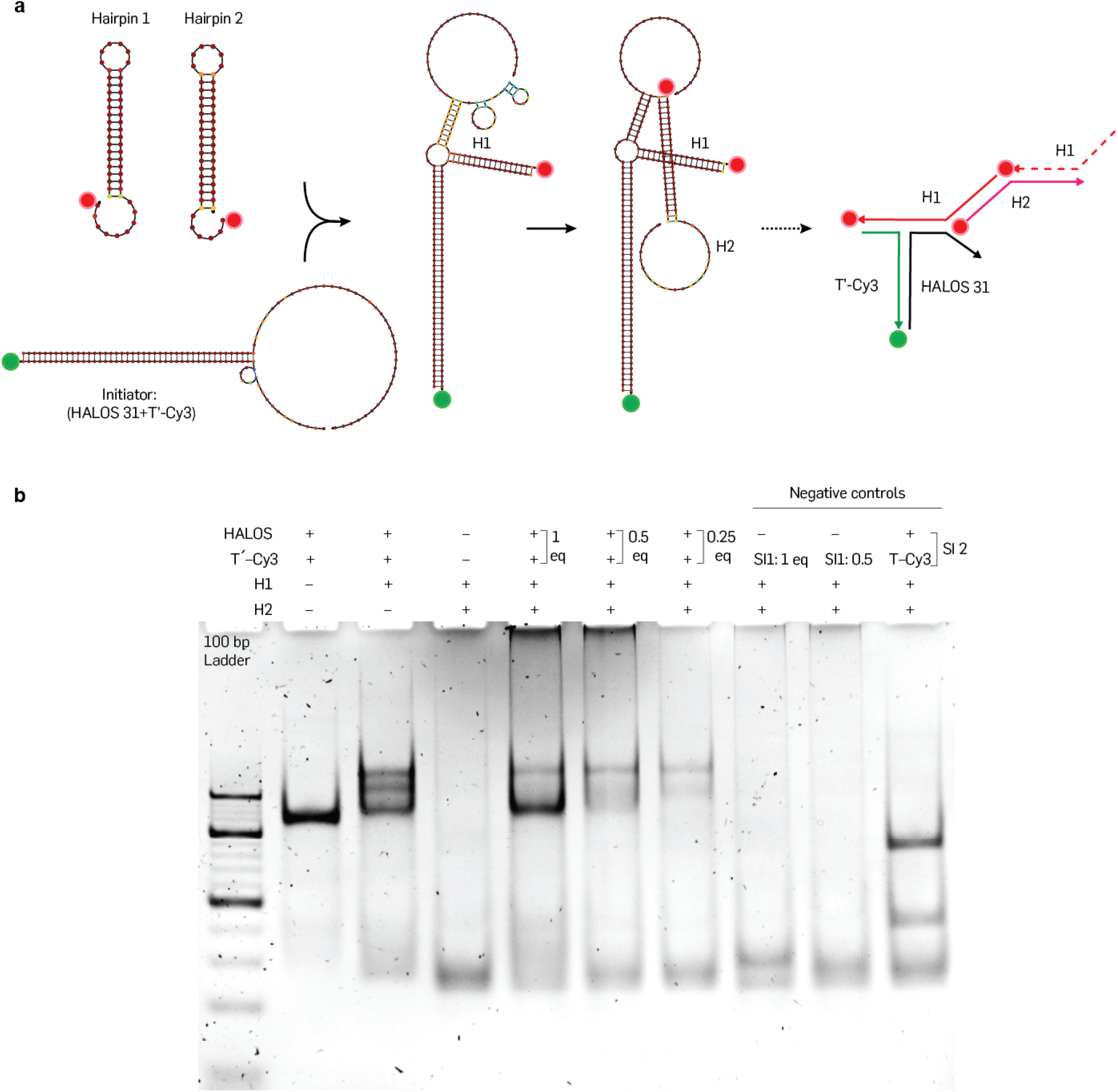
NUPACK simulation and native PAGE analysis of ψ-HCR-mediated signal amplification. (**a**) Extended target (T′-Cy3, 60-nt) triggered hairpin opening in HALOS and initiation of ψ-HCR by proximity induced split-initiator generation. NUPACK-simulated structure of the metastable hairpin monomers (H1, H2) coupled on HALOS, depicting spontaneous opening of the hairpin monomers in the presence of split-initiator and their thermodynamic stability at 25 *^◦^*C. Sequences of H1 and H2 were designed with maximum thermodynamic energy gain with no cross-reactivity. (**b**) Native PAGE of ψ-HCR in solution (1× TAE+150 mM NaCl+12.5 mM MgCl_2_) supports the initiation of HCR with initiator (HALOS 31+T′-Cy3). HCR progress was monitored with varied initiator ratios (1 to 0.25 equiv.), while no HCR was observed in the absence of T′-Cy3 and HALOS 31. Negative controls lacking either split initiator (SI) 1 (extended target domain) or 2 (open hairpin domain) resulted in no ψ-HCR product formation, confirming the requirement of both split initiators for the reaction. A 10% native PAGE was run in 1× TBE buffer with 12.5 mM MgCl_2_ at 100 V for 60 min at room temperature, post-stained with EtBr (0.5 µg/mL, 10 min) and imaged using UV-transillumination.

**Fig. S21.**
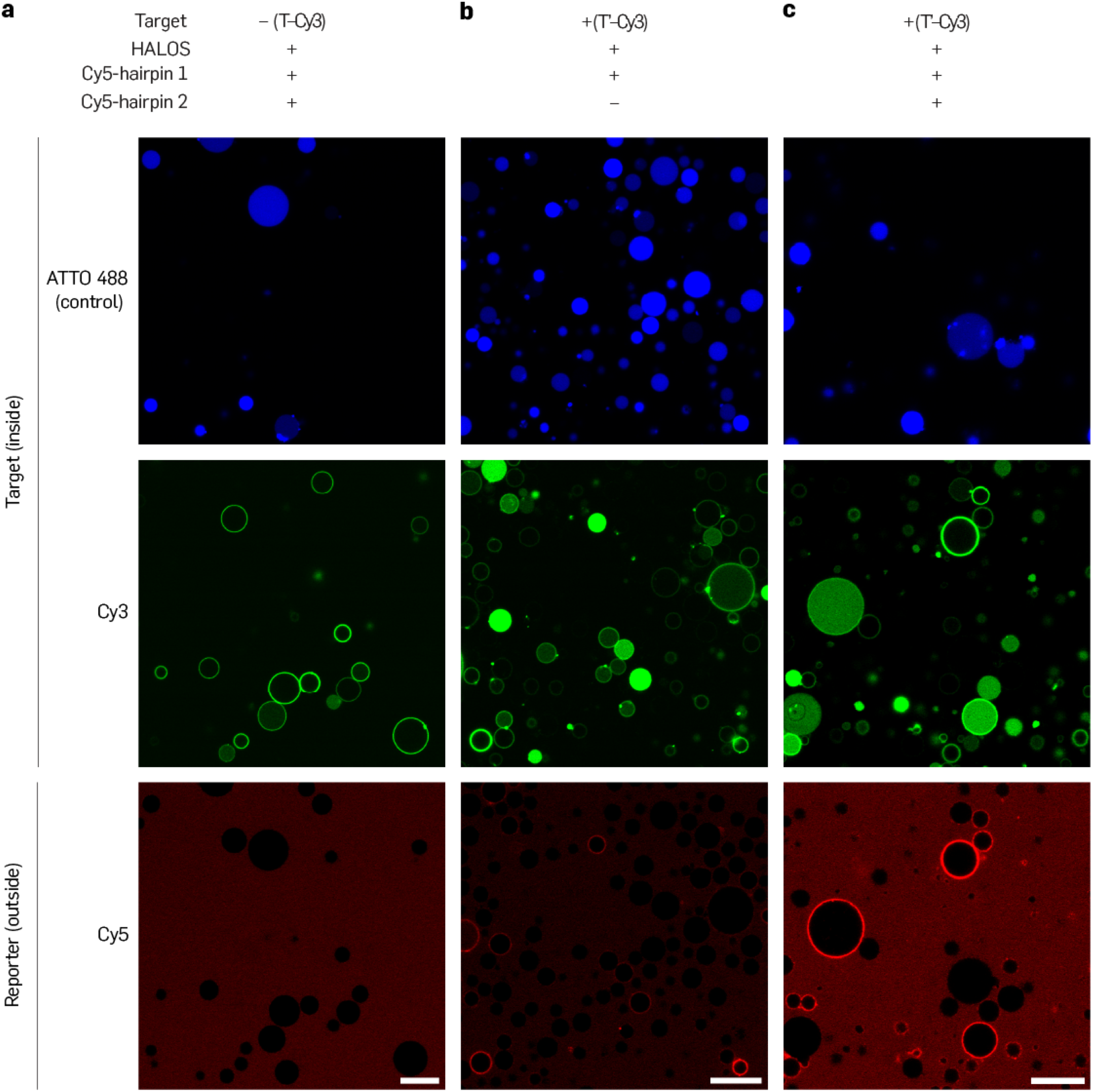
Signal amplification in GUVs confirms proximity-based initiation of ψ-HCR in GUVs. (**a**–**c**) Confocal images demonstrate ψ-HCR-based signal amplification from extended target-encapsulated (T′-Cy3) GUVs. Control GUVs (ATTO488) with no targets and in the absence of either initiator domain with non-extended target (T-Cy3, 41-nt) confirmed no progression of ψ-HCR (a). With extended target (T′-Cy3, 60-nt), reporter H1 produced low signal-to-noise, while ψ-HCR enabled pronounced signal amplification, confirming proximity induced initiation of HCR (b–c). Scale bar: 20 µm (a–c).

**Fig. S22.**
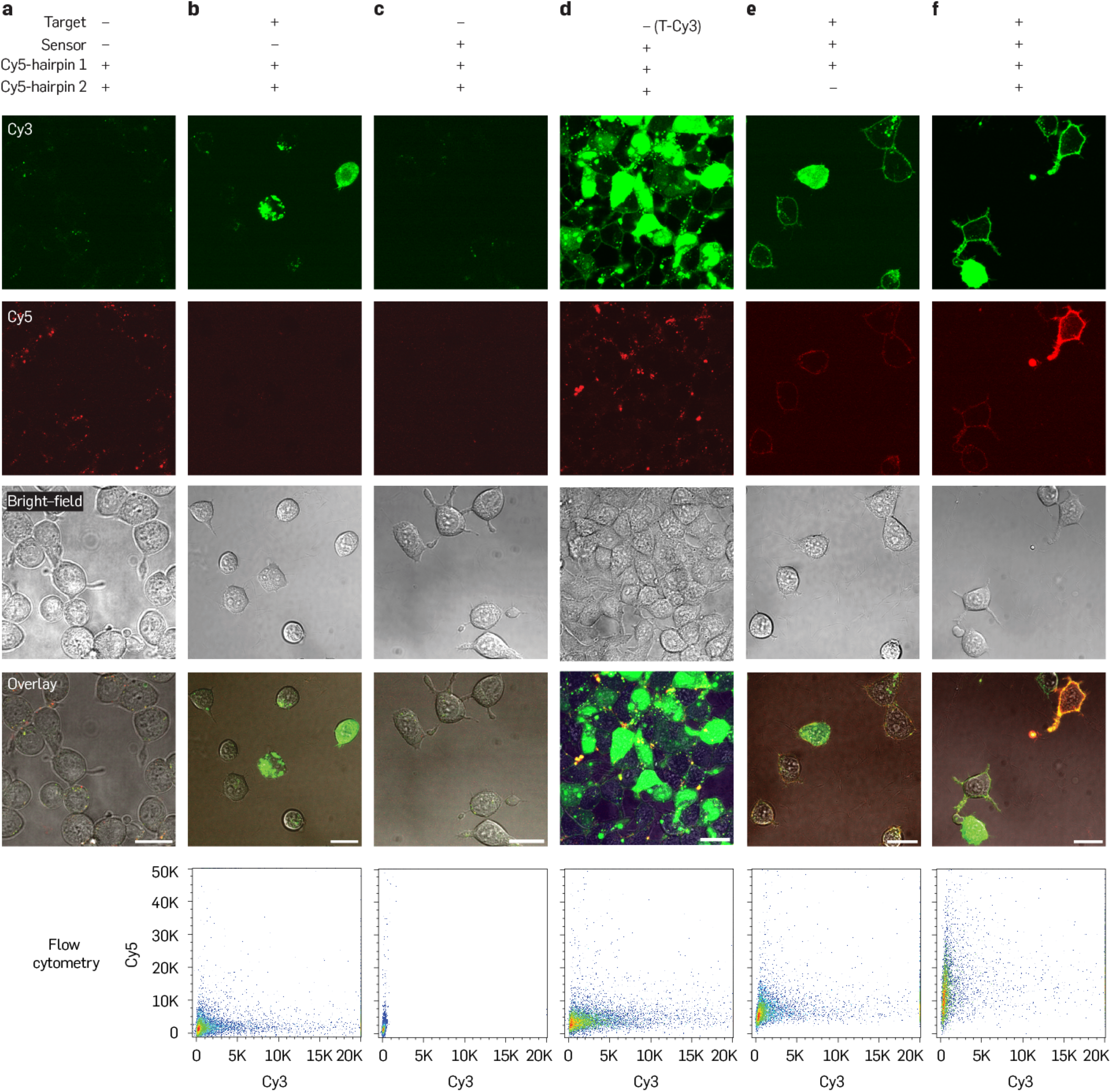
Signal amplification of HALOS-mediated RNA detection in live A549 cells using ψ-HCR. (**a**–**f**) Additional confocal microscopy images of signal amplification for HALOS-mediated RNA detection and amplification using ψ-HCR in live A549 cells (presented in Fig. 7g). Cells were transfected with 2′-OMe-RNA target (2′-OMe-T′-Cy3, 60-nt) using lipofectamine-based transfection. Negative controls lacking HALOS, target RNA, or both, displayed no membrane Cy3 and Cy5 fluorescence (a–c). Furthermore, in the absence of an extended domain in the target RNA (2′-OMe-T-Cy3, 41nt), target detection using membrane Cy3 labeling but without an amplified Cy5 signal indicates failure to initiate ψ-HCR (d). Extended target RNA (60-nt)-transfected cells treated with HALOS 31 exhibited a bright Cy5 signal upon performing ψ-HCR reaction (60 min in cell-culture environment at 37 *^◦^*C) with substantially higher Cy5 intensity for HCR versus H1 alone (e–f). Flow cytometry Cy3 vs. Cy5 scatter plots corroborated the amplified Cy5 surface signal for HCR compared to H1 and negative controls. Scale bar: 20 µm (a–f).

### Supplementary Movies

**Movie S1 All-atom molecular dynamics of HALOS 31 membrane anchoring and orientation.** One-microsecond all-atom MD trajectory of HALOS 31 in an aqueous solution and embedded within a lipid bilayer. In the absence of a membrane, the double-stranded stem remains largely intact, but the two cholesterol moieties stack and intercalate between the DNA bases to avoid the aqueous environment, producing pronounced helical distortion and a small inter-cholesterol angle (0–30*^◦^*). At the membrane, the small-angle lateral state similarly drives cholesterol stacking and stem distortion, whereas a large inter-cholesterol angle (100–210*^◦^*) stabilizes an axial orientation in which HALOS 31 sits perpendicular to the bilayer with an intact stem, positioning the toehold domain for target recognition. The DNA backbone is shown as a tube, and the nitrogenous bases are shown as spheres. The lipid volume is rendered as a transparent surface, with water and ions omitted for clarity.

**Movie S2 All-atom molecular dynamics of target-bound HALOS 31 supporting transmembrane signal transduction.** One-microsecond all-atom MD trajectory of target-bound HALOS 31 at the membrane in intermediate and open hairpin conformations. Target binding promotes the expulsion of the non-cholesterol stem arm (b*) from the bilayer into the outer aqueous solution, accompanied by an increased hydrogen-bond count (mean±SD= 87.9±9.5) and a large inter-cholesterol angle distribution (60–150*^◦^*), confirming stable membrane residence and the structural basis for signal transduction across the bilayer. Representation is identical to that in Movie S1.

